# A Potent Kalihinol Analogue Disrupts Apicoplast Function and Vesicular Trafficking in *P. falciparum* Malaria

**DOI:** 10.1101/2023.11.21.568162

**Authors:** Z Chahine, S Abel, T Hollin, JH Chung, GL Barnes, ME Daub, I Renard, JY Choi, V Pratap, A Pal, M Alba-Argomaniz, CAS Banks, J Kirkwood, A Saraf, I Camino, P Castaneda, MC Cuevas, J De Mercado-Arnanz, E Fernandez-Alvaro, A Garcia-Perez, N Ibarz, S Viera-Morilla, J Prudhomme, CJ Joyner, AK Bei, L Florens, C Ben Mamoun, CD Vanderwal, KG Le Roch

## Abstract

Here we report the discovery of MED6-189, a new analogue of the kalihinol family of isocyanoterpene (ICT) natural products. MED6-189 is effective against drug-sensitive and-resistant *P. falciparum* strains blocking both intraerythrocytic asexual replication and sexual differentiation. This compound was also effective against *P. knowlesi* and *P. cynomolgi*. In vivo efficacy studies using a humanized mouse model of malaria confirms strong efficacy of the compound in animals with no apparent hemolytic activity or apparent toxicity. Complementary chemical biology, molecular biology, genomics and cell biological analyses revealed that MED6-189 primarily targets the parasite apicoplast and acts by inhibiting lipid biogenesis and cellular trafficking. Genetic analyses in *P. falciparum* revealed that a mutation in *PfSec13*, which encodes a component of the parasite secretory machinery, reduced susceptibility to the drug. The high potency of MED6-189 *in vitro* and *in vivo*, its broad range of efficacy, excellent therapeutic profile, and unique mode of action make it an excellent addition to the antimalarial drug pipeline.

**Editor’s Summary:** Here we report the mode of action and mechanism of resistance of a pan-antimalarial agent, MED6-189, which disrupts apicoplast function and vesicular trafficking in *P. falciparum*.

## INTRODUCTION

In 2022, an estimated 247 million clinical cases and 619,000 global deaths were due to malaria worldwide, most of which were caused by *Plasmodium falciparum* (*1*). Because of widespread resistance to commonly used antimalarials, artemisinin-based combination therapies (ACTs) represent the last resort in the antimalarial armamentarium for management of drug resistance malaria (*2–5*). Unfortunately, there has been a gradual increase in resistance to ACTs, particularly in the Greater Mekong region of Southeast Asia (*6–8*). There are presently only a small number of antimalarials with efficacy comparable to that of ACTs, and current vaccines have only limited protective efficacy (*9, 10*). There is therefore an urgent need to identify novel therapeutics to combat the ever-growing threat of drug resistance and to suppress the spread of the disease. Ideally, such molecules should target novel pathways not previously targeted by approved antimalarials or those in clinical development.

The isocyanoterpene (ICT) family of sponge-derived natural products encompasses a large number of biosynthetically related, isonitrile-, isothiocyanate-, isocyanate-, and formamide-containing diterpenoids, many of which have been shown to have potent antibacterial, antifungal, and antimalarial activity (*11–13*). The presence of the isonitrile functional group is critical for potent antimalarial activity. Among the ICTs, the kalihinol subfamily counts some of the most active compounds, with kalihinols A (*11, 12, 14*) and B (*14–17*) exhibiting potent activity against drug-sensitive and-resistant *P. falciparum* isolates (*16, 17*). Our labs previously completed the first synthesis and testing of kalihinol B; in demonstrating that its activity was similar to that of kalihinol A, we learned that the oxygen heterocycle motif was likely not critical for activity. We therefore dramatically truncated this group, which ultimately led to the discovery of analogue MED6-189, whose simplified structure allowed for an easier synthesis.

Efforts to understand the mechanism of action of the ICT family go back more than two decades (*18–20*). Wright, Tilley, and co-workers showed that some ICTs can bind free heme, as well as inhibit the formation of its crystalline form, β-hematin (*19*). This study suggested that inhibition of heme detoxification is an important cause of toxicity to the malaria parasite (*19*). However, it focused predominantly on the tetracyclic cycloamphilectane and tricyclic amphilectane ICTs, and no kalihinols were investigated. More recently, another study showed that compounds in these same two structural classes inhibit both intraerythrocytic replication and liver stage development of malaria parasites (*21*), indicating that heme detoxification might not be the primary mechanism by which ICTs exert their antiplasmodial activity.

We show that MED6-189, one of our most accessible synthetic analogues of kalihinols A and B, and closely related chemical probe molecules, inhibits drug-sensitive and - resistant *P. falciparum* strains in vitro, blocks parasite sexual differentiation, and eliminates infection in a humanized mouse model of *P. falciparum* malaria. Using a systems biology strategy that integrates various multi-omics platforms, we gained an unbiased understanding of both the drug mechanism of action and possible routes of resistance. Our findings demonstrate that MED6-189 (and by extension, kalihinols and presumably other ICTs) disrupts the apicoplast, an organelle essential for the synthesis of parasite fatty acids and isoprenoids. Metabolomic and proteomic analyses show that the compound interferes with lipid metabolism and trafficking during the parasite asexual stage. Moreover, we found that mutations in the *PfSec13* gene (*PF3D7_1230700*), an important component of the protein secretory machinery, is associated with altered susceptibility to the drug. Finally, the compound was also found to be potent against other zoonotic/human *Plasmodium* parasites, *P. knowlesi* and *P. cynomolgi*, thus highlighting its potential as a pan-antimalarial drug.

## RESULTS

### Generation of synthetic kalihinol analogues

Because large-scale production of the ICT derived natural products is not sustainable, we synthesized kalihinol B (Fig. 1A, left), a tetrahydrofuran isomer of kalihinol A (a tetrahydropyran), which showed strong activity against *P. falciparum* Dd2 isolate with IC_50_ values of 4.6 nM (*16*). Because this compound was still challenging to generate on a sufficiently large scale for in-depth mechanism of action studies, several less complex synthetic analogues of kalihinols A and B were generated and their antiparasitic activity evaluated in vitro (See Supporting Information S1 and fig. S1)(*16*). Among these compounds, MED6-189 emerged as a highly potent, more accessible drug with IC_50_ values below 50 nM against several *P. falciparum* strains (Fig. 1A, right) (*16*). Through a robust sequence of ten chemical steps, the synthesis of MED6-189 was scaled to support the in-depth studies detailed in this work (See Supporting Information S1 and fig. S1)(*22, 23*).

**Figure 1.**
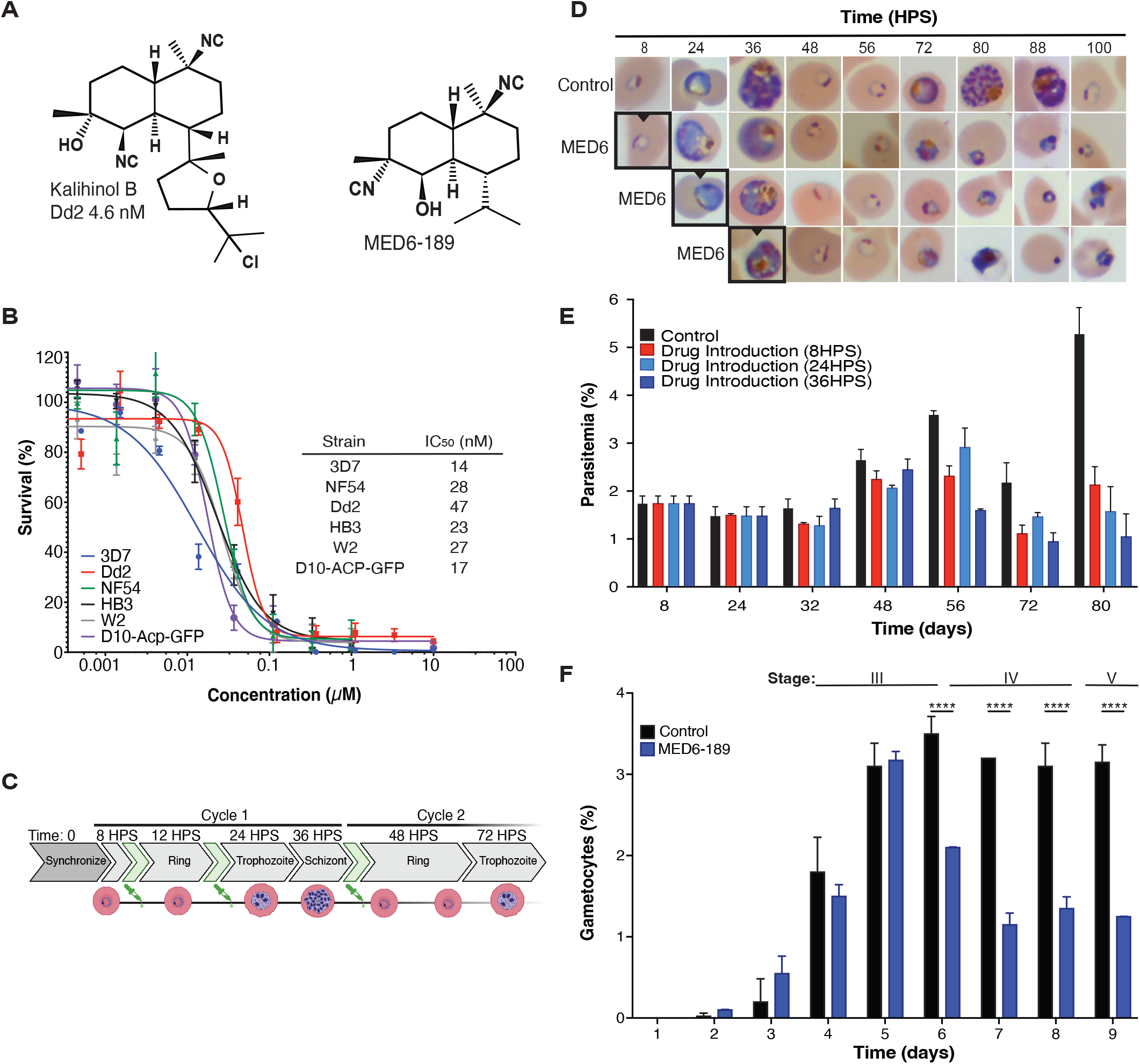
Effect of MED6-189 on *P. falciparum* intraerythrocytic development. **A.** Chemical structures of the natural product kalihinol B (left) and its analog MED6-189 (*16, 17*). **B.** SYBR Green-based dose response assays were conducted on early-ring stage parasites (6 hours post invasion). The parasites were exposed to serial dilutions of MED6-189 for 72 hours, after which parasite growth was assessed. 3D7 WT (blue), NF54 (green) and drug-resistant strains Dd2 (red), HB3 (black), W2 (grey) and D10-Acp-GFP (purple) lines (Sigmoidal, 4PL, X is concentration, n≥3, nonlinear regression, CI:95%). **C.** Schematic diagram of the development of *P. falciparum* following two consecutive erythrocytic cycles. The time points at which MED6-189 was introduced are depicted in green. **D.** Giemsa-stained images of synchronized 3D7 parasites that were incubated with either DMSO or MED6-189 (at its IC_80_ concentration). The images depict various developmental stages of the parasite’s intraerythrocytic life cycle. Bordered images represent timepoints when drug is first introduced (n=3, p<0.05) **E.** (%) parasitemia following exposure of 3D7 parasites to either DMSO (Control) or MED6-189 at various stages of the parasite life cycle within erythrocytes (p< 0.05, n=3, 2-way ANOVA, Tukey t-test). **F.** Inhibition of *P. falciparum* gametocyte development following MED6-189 treatment (blue) during early gametocytogenesis compared to the control (black) (p<0.05, n=3, 2-way ANOVA).

### MED6-189 affects *P. falciparum* intraerythrocytic life cycle

To assess the effectiveness of MED6-189, (Fig. 1A, right) on the intraerythrocytic development (IED) of *P. falciparum*, we performed growth inhibition assays on several isolates with established sensitivity or resistance to various classes of antimalarials (Fig. 1B). MED6-189 showed excellent activity against both drug-sensitive strains (3D7 and NF54), with IC_50_ values of 14 nM ± 2 nM and 28 nM ±5 nM, respectively, and pyrimethamine-and chloroquine-resistant strains (HB3, Dd2 and W2) with IC_50_ values of 23 nM ± 2 nM, 47 nM ± 7 nM, and 27 nM ± 5 nM, respectively (Sigmoidal, 4PL, X is concentration, n≥3, nonlinear regression, CI:95%) (Fig. 1B, table S1A). In order to identify the specific stage of the *P. falciparum* asexual intraerythrocytic cycle (IEC) that was predominantly inhibited by the compound, we carried out phenotypic analyses on synchronized cultures that were incubated with either MED6-189 or a control vehicle (DMSO) at different time intervals following synchronization (Fig. 1C). Whereas no significant impact of the compound on parasite development could be observed during the first IEC, significant changes including cell cycle arrest and cellular stress, were evident during the trophozoite stage of subsequent IEC (Fig. 1D). The treated parasites were unable to fully undergo schizogony or re-invasion (p< 0.05, n=3, 2-way ANOVA, Tukey t-test) (Fig. 1E). This phenotype is reminiscent of that of compounds that target the apicoplast, such as fosmidomycin (*24–26*). To assess whether MED6-189 could inhibit the parasite’s ability to undergo sexual differentiation, a critical step in malaria transmission, we examined *P. falciparum* gametocyte development in the absence or presence of the compound. MED6-189 inhibited early gametocyte development (Stage III-Days 4 to 6) (Fig. 1F) by ∼62% compared to controls (p<0.05, n=3, 2-way ANOVA). Furthermore, when exposed to high doses of MED6-189 (1µM) during the early stages of gametocyte development, parasite growth was abolished, and after 48h of drug treatment, no quantifiable gametocytes were observed (fig. S2A). In contrast, the compound had little to no effect on mature-stage gametocytes (Stage V) (Fig. 1F, S2A, table S1B).

### MED6-189 targets apicoplast metabolism

In order to gain deeper insights into the cellular metabolic target of MED6-189, we synthesized a fluorescently-labeled kalihinol analogue, MED6-131, featuring a JF549 fluorophore attachment (see MED6-131 structure in fig. S2B) and assessed its activity and cellular localization. Cell growth assays confirmed its potent action against *P. falciparum*, exhibiting an IC_50_ of 17.23 nM ± 0.95. Fluorescence microscopy analyses were conducted using D10-Acp-GFP transgenic parasites, which express a fusion of the Acyl Carrier Protein (Acp) with GFP in the parasite apicoplast (*27*). These analyses revealed the co-localization of MED6-131 with Acp-GFP during the trophozoite and schizont stages of the parasite life cycle (Fig. 2A). These findings affirm the compound’s localization within the apicoplast.

**Figure 2.**
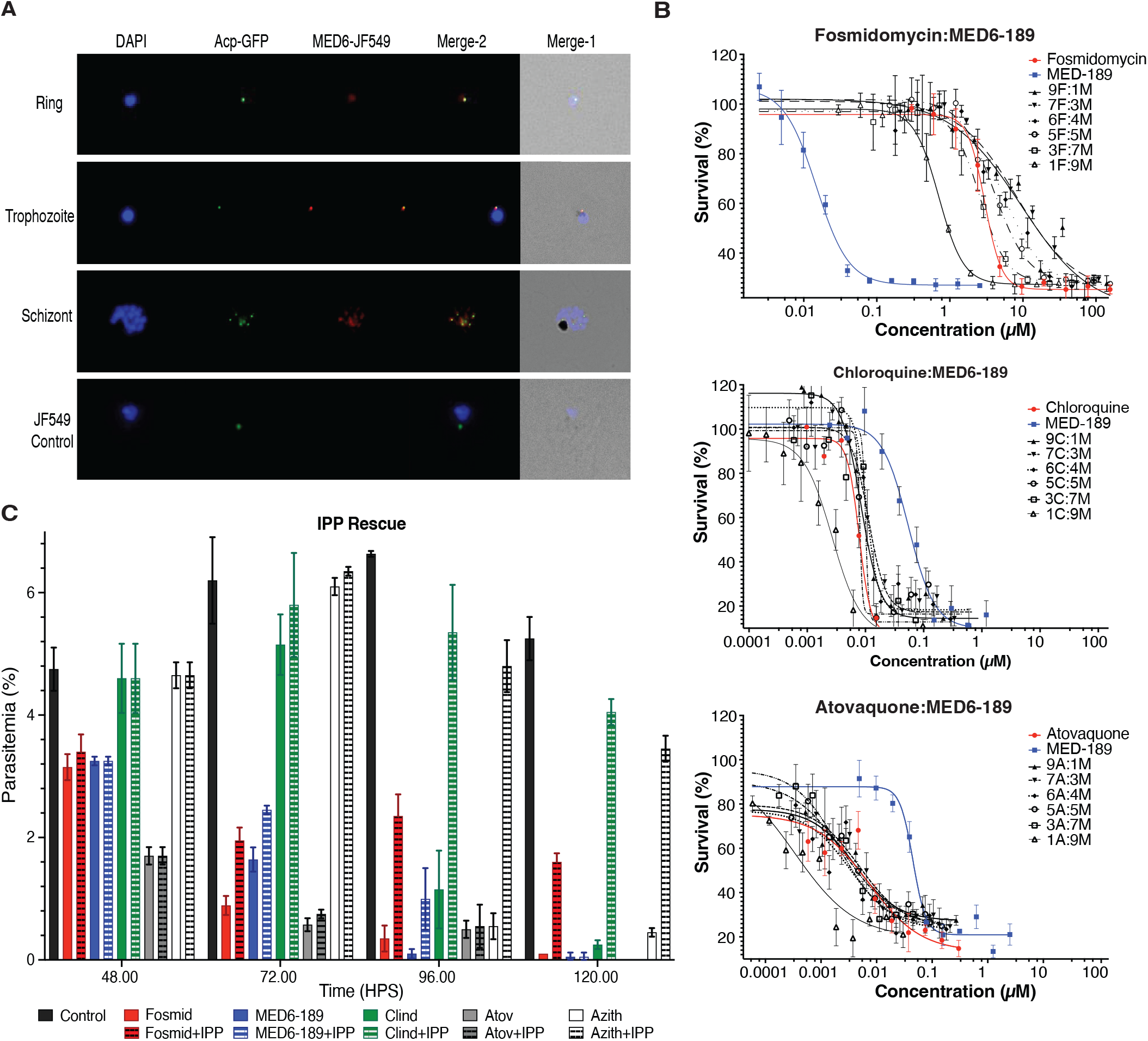
MED6-189’s localization and activity in combination with other antimalarials. **A.** Cellular localization of MED6-131 (Red) in D10-Acp-GFP *P. falciparum* transgenic parasites. Nuclei are stained with DAPI (Blue). Overlap between ACP-GFP (Green) and MED6-131 can be seen during the trophozoite and schizont stages of the cell cycle. **B.** Dose-dependent interactions between MED6-189 (Blue) and various antimalarials with known mechanisms of action (Red). The figures show logarithmic growth of parasites (Y-axis) as a function of drug concentrations for MED6-189 (M), Fosmidomycin (F), Chloroquine (C) or Atovaquone (A) (X-axis). The regression line represents a nonlinear regression (Variable slope with four parameters), with significant differences considered if p<0.05. Activity correlations between each compound and MED6-189 were analyzed using Pearson correlation (r) using GraphPad Prism 9 (GraphPad Software, Inc.), n=3 (See table S2, fig. S3). **C.** Rescue of 3D7 parasites exposed to DMSO (black), Fosmidomycin (Red), MED6-189 (Blue), along with several other known antimalarials supplemented with IPP 48 post-synchronization (dotted lines). The analysis was performed using a 2-way ANOVA, n=3, with p<0.05 significance (See table S2C).

To explore whether MED6-189 targets the apicoplast non-mevalonate MEP/DOXP pathway, which is essential for parasite survival (*26*), we initiated drug-drug interaction studies. Our data demonstrated that MED6-189 and fosmidomycin, a known inhibitor of the MEP/DOXP pathway, exhibited antagonistic effects, as indicated by an estimated Fractional Inhibitory Concentration (FIC) index of 2.4 ± 0.36 (Fig. 2B). Interactions of MED6-189 with chloroquine and atovaquone, drugs unrelated to apicoplast functions, were primarily additive (chloroquine (FIC score= 1.71 ± 0.4) and atovaquone (FIC score = 0.91 ± 0.2), (Fig. 2B, table S2A and B, fig. S3) *(28, 29)*.

To substantiate MED6-189’s apicoplast targeting, we investigated the effect of isopentenyl pyrophosphate (IPP) supplementation on MED6-189’s antimalarial activity. IPP is a product of the MEP/DOXP pathway, and its supplementation reduces susceptibility to MEP inhibitors. Synchronized parasites were treated with either vehicle control (DMSO), MED6-189, fosmidomycin or other known antimalarial agents (*30–32*) at IC_80_ concentrations for one intraerythrocytic cycle prior to IPP (200 µM) supplementation. In contrast to samples not supplemented with IPP, parasites supplemented with IPP survived the second IEC and initiated a third cycle in the presence of MED6-189 or fosmidomycin (Fig. 2C, table S2C). Our results demonstrate that MED6-189 inhibition can only be rescued by IPP for one additional cycle. Together, these findings indicate that, in addition to its inhibition of the apicoplast, MED6-189 likely hinders one or more additional targets within the parasite.

### Multi-omics approaches unveil the mechanism of action of MED6-189

To elucidate the cellular and metabolic pathways affected by MED6-189, we employed a comprehensive multi-omics strategy. First, we analyzed transcriptional changes by conducting RNA-seq on *P. falciparum* (3D7 strain) in the absence or presence of MED6-189 at various time points during the parasite IEC. Whereas no significant change in transcript levels were detected between the control and drug-treated samples during the first IEC (table S3A), significant changes were manifest during the late ring and trophozoite stages of the second IEC, consistent with the delayed inhibitory activity of MED6-189 on parasite development (log_2_ fold-change (FC) >0.5 or <-0.5) (table S3A, S3B). Gene Ontology (GO) enrichment analyses identified several classes of transcripts that were significantly downregulated following exposure to MED6-189, including those known to be involved in parasite invasion and egress, such as *SUB1/SUB2* genes (PF3D7_0507500/PF3D7_1136900). These findings suggest that the compound induces cell cycle arrest. Among the upregulated genes, we identified those involved in the isoprenoid catabolic process and apicoplast function, such as G3PAT, autophagy, and stress responses (Fig. 3A, table S3C, fig. S4) (*33*). Although most changes detected in transcript level in the second IEC were likely induced by the drug in a nonspecific manner (cell cycle arrest and stress induction), our results may also suggest a potential compensatory mechanism of the pathways affecting isoprenoid catabolic process and the apicoplast.

**Figure 3.**
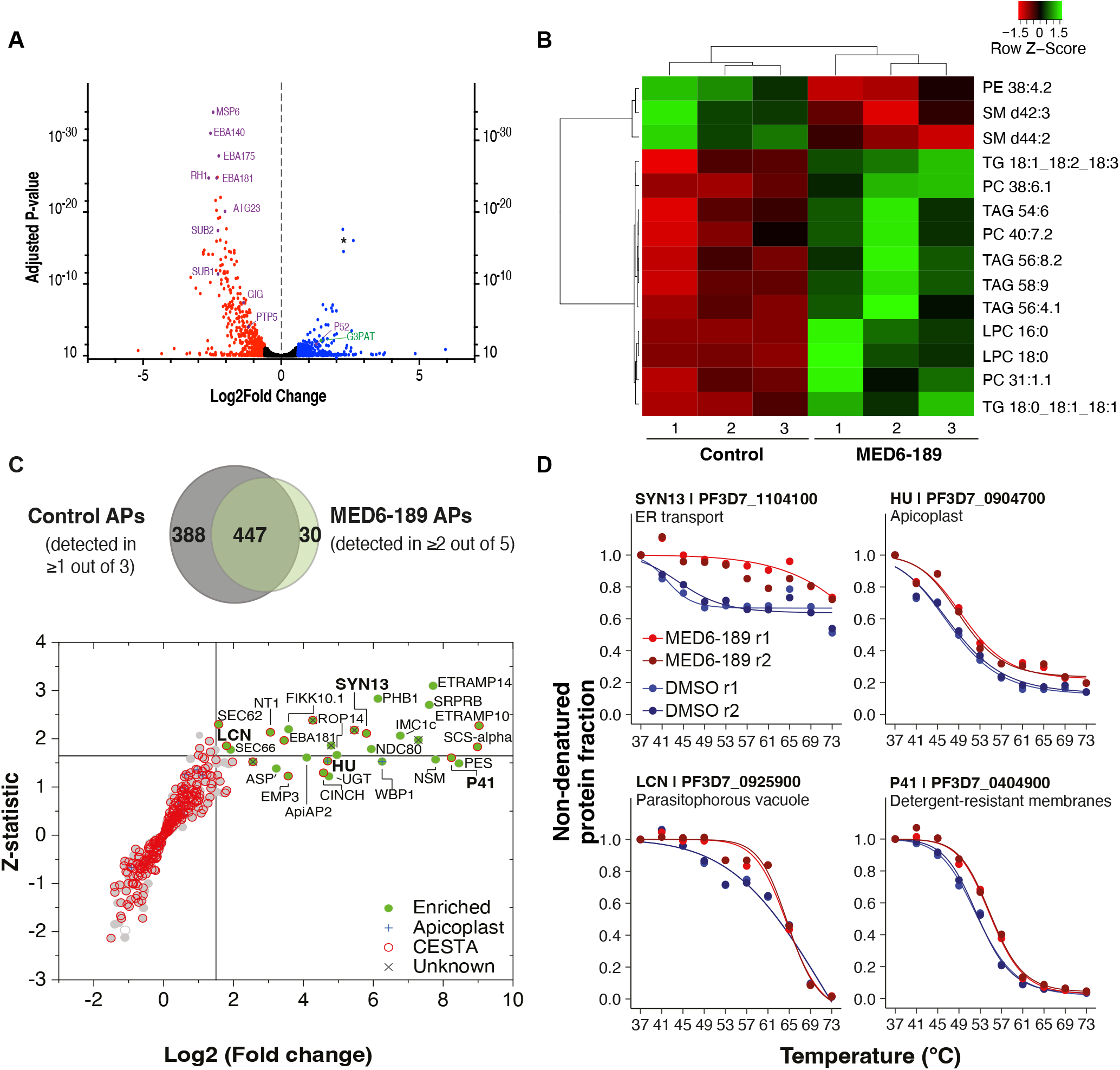
Omics-based profiling of MED6-189 treated parasites. **A.** Volcano plot representing transcriptomic changes induced by MED6-189 treatment. A total of 5712 transcripts were identified with an adjusted p-value cut-off of 0.05. Transcripts associated with invasion and stress responses are highlighted in purple and those related to apicoplast function in green (**\*** represent ncRNAs of unknown function). **B.** Heatmap depicting the regulation of lipid metabolism in response to MED6-189. Metabolites significantly upregulated in response to MED6-189 treatment are shown in green and those downregulated in red. We used a Log_2_ transformation to the data for the calculation of q-values (Benjamini-Hochberg adjusted p-values) and p-values using *Welch’s t-test* or *ANOVA*. **C.** Protein pulldown assays using biotinylated kalihinol analogue, MED6-118. The significance plot displays all proteins detected in at least two of the five independent MED6-118-based affinity purifications (APs). Scatter plots with gray dots depict QPROT-derived log_2_(FC) and Z-statistic values between MED6-APs and negative controls (table S4). Significantly enriched proteins with a Log_2_(FC)≥1.5 and a Z score ≥1.645 or those not detected in controls are highlighted in green. Proteins for which thermal profiles are available are shown in red (table S5 and Supporting Information S2). Proteins localized to the apicoplast are indicated with a blue cross, while proteins of unknown function are marked with a gray “X”. A Venn diagram shows the protein overlap between the MED6-118 APs and controls. **E.** CETSA melt-curve analysis of *P. falciparum* lysates treated with MED6-189. The thermal profiles for four *P. falciparum* proteins significantly enriched in the MED6-118-based pull-downs are shown. Stabilization is assessed on the relative amount of soluble protein remaining (Y-axis) after thermal challenge (X-axis). Sample replicates are color-coded in shades of red for MED6-189-treated samples and blue for DMSO controls.

To assess the effect of MED6-189 on cellular metabolism, highly synchronized parasite cultures were maintained in the absence or presence of the compound and their metabolomic profiles were examined during the second IEC (72hrs post-treatment). A total of 1178 metabolites were analyzed (see methods), of which 40 were significantly affected by MED6-189 treatment. These included 2 polar metabolites (thiamine pyrophosphate (TPP) and S-adenosyl methionine (SAM)) and 38 lipids, including components of the Kennedy pathway for the synthesis of phosphatidylcholine and phosphatidylethanolamine (PC and PE, respectively)*(34, 35)* (fig. S5, table S5). TPP is essential for the activity of several enzymes including the apicoplast-associated pyruvate dehydrogenase (PDH) and vitamin B1 biosynthesis (*36–38*), and SAM is a major precursor for the synthesis of PC from the SDPM pathway (*39*). This suggests an impact of MED6-189 on apicoplast biogenesis and membrane biosynthesis (*34, 39*).

### MED6-189 binds to *P. falciparum* proteins involved in membrane trafficking

To identify proteins that interact with MED6-189, a biotinylated derivative, MED6-118, (fig. S2B) with a similarly potent antimalarial activity (IC_50_= 92.4 nM ± 2.0), was synthesized and used in pull-down assays using *P. falciparum* protein extracts. The purified proteins were analyzed through Multidimensional Protein Identification Technology (MudPIT) and quantified using dNSAF spectral counting (table S4). Over 450 proteins were detected in at least two of the five biological replicates (Fig. 3D). Among them, 30 proteins were either absent in the negative controls or significantly enriched, as determined by Log_2_(FC) ≥1.5 and Z-statistic ≥1.645, calculated using QPROT (*40*). These enriched proteins are involved in vesicular trafficking, lipid biogenesis and signaling (Fig. 3E). Specifically, three proteins known to be involved in the endoplasmic reticulum (ER) COPII trafficking systems Sec62 (PF3D7_1438100), syntaxin (PF3D7_1104100), a member of the Qa-SNARE family, and a conserved protein recently annotated as a putative translocation protein SEC66 (PF3D7_0204200) were found to be significantly enriched in the pull-down experiments from the MED6-189-treated but not the vehicle control samples (*41–45*).

To further investigate protein interactions with MED6-189, we conducted Cellular Thermal Shift Assays (CETSA) (*46*) using whole protein extracts purified from either vehicle or MED6-189 treated parasite cultures (*46*). Following the addition of the ligand, the samples were incubated at temperatures ranging between 37°C and 73°C, and soluble proteins were then analyzed via liquid chromatography coupled to tandem mass spectrometry, as described previously (*47*). The data was processed using the mineCETSA R package (*47*), from which melting curves were generated for over 800 *P. falciparum* proteins (table S5 and Supporting Information S2)(*48, 49*). Among them, 30 proteins exhibited stabilization and displayed a pattern of temperature-mediated stabilization (melting curves) from which 14 were also detected as enriched in the affinity purification experiments (Fig. 3E and S6A). Notably, SYN13 exhibited the most significant shift in melting temperature (Fig. 3E), while the other three proteins, namely the apicoplast localized bacterial histone-like protein (HU | PF3D7_0904700), lipocalin (LCN | PF3D7_0925900), and 6-cysteine protein (P41 | PF3D7_0404900), displayed a 2-3°C difference between the untreated and MED6-189 treated samples.

Other proteins of interest with substantial thermal shifts upon MED6-189 treatment included UIS2 (PF3D7_1464600), a serine/threonine protein phosphatase localized to the parasitophorous vacuole, and PyKII (PF3D7_1037100) an apicoplast pyruvate kinase. While these proteins were detected in the affinity purification dataset, they did not meet the significance cut-off (fig. S6B). Finally, a few proteins were not detected in the MED6-189 pull-downs but exhibited significant stabilization (PF3D7_0724100, PF3D7_1466900), or destabilization in the case of BOP1 (PF3D7_1405800), a protein involved in ribosome biogenesis (fig. S6C). Collectively, these two complementary proteomics approaches suggest a possible mechanism of action of MED6-189 involving the association and potential disruption of the membrane trafficking apparatus between the ER-Golgi or ER-apicoplast systems.

### Selection of MED6-189 refractory parasites

To gain further insights into the cellular machineries impacted by MED6-189, we conducted extended *in vitro* resistance selection experiments (*50–52*). 3D7 parasites were first cloned through serial dilution (*53*) and isolated clones were split into controls or experimental lines, and cultured in the presence of MED6-189 at IC_50_ values before gradually escalating the drug dosage (Fig. 4A). After approximately 36 months of continuous in vitro propagation, parasites exhibiting noticeable drug tolerance emerged. Drug inhibition assays of these selected cloned lines revealed a 2 to 4-fold decrease in susceptibility to MED6-189 compared to wild type clones (3D7). The IC_50_ values for the drug-sensitive parental lines were approximately 14± 1 nM, while those for the resistant lines were approximately 60 ± 6 nM (Sigmoidal, 4PL, X is concentration, n=3, nonlinear regression, CI:95%) (Fig. 4A, table S1A). Whole genome sequencing (WGS) identified single-nucleotide polymorphisms (SNPs) or Indels in two coding genes: *PfSec13* (PF3D7_1230700) and endoplasmin (PF3D7_1222300) (table S6) in the mutant lines, but not in in the parental lines. Among these, *PfSec13* appeared to be the most promising target, owing to its established role in protein trafficking (fig. S6F) (*54, 55*). While SEC13 was not pulled down by MED6-189 affinity purification, both replicate thermal profiles were consistently stabilized in the presence of MED6-189 (fig. S6G). The role of the *PfSec13* gene in susceptibility to MED6-189 was confirmed in *Saccharomyces cerevisiae*. Since the yeast *SEC13* gene is essential for viability, we assessed the effect of MED6-189 following overexpression of Sec13p using a multi-copy plasmid or its repression using a tetracycline-regulated (Tet-off) promoter. Sec13p overexpression significantly increased resistance to the drug by 2.4-fold, whereas its repression was accompanied with a ∼5.5-fold increase in susceptibility to MED6-189 (Fig. 4F). Alteration of the expression levels of the endoplasmin in yeast did not result in discernible changes in susceptibility to MED6-189 (Data not shown).

**Figure 4.**
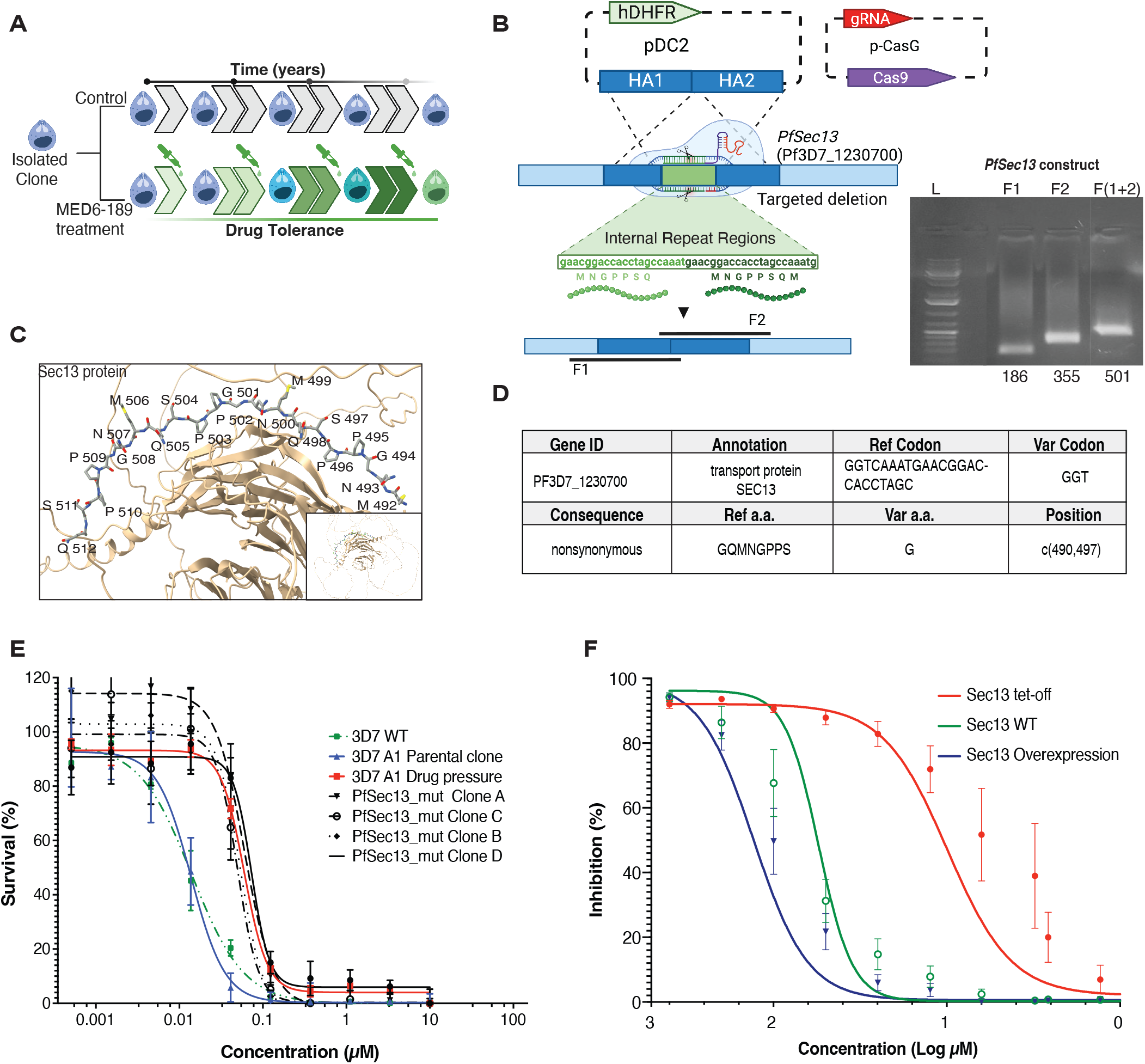
Evidence for a role of *Sec13* in susceptibility to MED6-189. Graphical illustration of the methodology employed to isolate MED6-189 resistant parasites. **B.** Predicted structure of the Sec13 protein with the seven amino acid tandem repeat region targeted for deletion (red). **C.** Schematic representation of the CRISPR-Cas9-based replacement strategy used to introduce a deletion of the targeted repeat regions in the *PfSec13* gene. The insertion was achieved through overlap extension PCR of fragments directly upstream and downstream of the target segment and subsequently formed by primer overlap extension PCR to replicate the desired deletion. The insertion was validated through whole genome sequencing. **D.** Results of the sequencing analysis, confirming the successful deletion of the tandem repeat region of *PfSec13* using CRISPR-Cas9 in an isolated clone. **E.** 3D7 WT and parental lines (blue), resistant lines (red) maintained in the presence of DMSO or MED6-189 and transgenic *PfSec13*-mut clones (dashed) were subjected to a parasite survival assay. The curves depict parasite survival (y-axis) in response to serial drug dilution of MED6-189 (x-axis). Data was analyzed using a Sigmoidal, 4PL (X represents concentration, n=3, nonlinear regression, CI:95%). **F.** Comparison of the growth rates of wild type *S. cerevisiae* and transgenic clones with either overexpressed or down-regulated Sec13p following treatment with a vehicle (DMSO) or increasing concentrations of MED6-189.

To validate the role of SEC13 in drug resistance in *P. falciparum*, we used the CRISPR-Cas9 gene editing tool. The seven amino acid deletion detected in the *PfSec13* gene in the resistant clones formed a tandem repeat sequence, which could potentially enhance protein structural integrity or be involved in protein-protein interactions (*56*). These tandem repeat sequences were targeted for deletion in the 3D7 wild type strain (Pf3D7_12_v3: 1261090-1261110 (+)) (Fig. 4B-C) using CRISPR-Cas9. The targeted deletion was verified through Sanger sequencing and WGS (Fig. 4D, table S6). Four *PfSec13-mut* transgenic clones were tested for their sensitivity to MED6-189. All *PfSec13-mut* clones exhibited tolerance levels to MED6-189 similar to those of the MED6-189 resistant lines obtained after drug selection (IC_50_ range: 50-73 nM, SEM ± 9 and 10 nM) compared to the wild-type strain (Fig. 4E, table S1A). Collectively, these genetic and pharmacological studies in both *P. falciparum* and *S. cerevisiae* underscore the crucial role of SEC13 in susceptibility to MED6-189.

### MED6-189 exhibits a favorable safety and tolerability profile

The strong potency of MED6-189 against *P. falciparum* in vitro coupled with a comprehensive understanding of its mechanism of action, prompted us to explore its efficacy in an animal model of *P. falciparum* malaria. The in vitro safety profile of MED6-189 was first determined by assessing possible effects on the growth and metabolic activity of five human cell lines (HeLa, THP1, HEK293, HepG2 and hTERT) at concentrations ranging between 48 nM and 100 µM. No inhibitory activity could be found, resulting in an estimated in vitro therapeutic index of > 500, surpassing that of several approved antimalarial drugs (table S7). MED6-189 was further evaluated in 15 in vitro assays using the Enhanced Cross Screen Panel (eXP), which provides pharmacological information and toxicology alerts for the target compound. Of these, hPXR (Human Pregnane-X-receptor) IC_50_ 7.4 µM & BSEP inhibition (Bile Salt Export Protein) IC_50_ 50 µM were highlighted as factors to be considered in future drug optimization studies aimed at identifying ideal partner drugs and mitigating potential hepatotoxicity. Given the susceptibility of isonitrile compounds to hydrolysis to formamides in acidic aqueous solution, we synthesized the formamide derivatives GB209-2 and GB209-3 by acidic hydrolysis of MED6-189 (see Supporting Information S1). These derivatives exhibited significantly decreased antiplasmodial activities (IC_50_ values of 3.8 and 3.9 µM, respectively), indicating that these potential degradation products are not responsible for the observed antimalarial effects of MED6-189 administration. We also demonstrated that MED6-189 had reasonable stability to aqueous acidic conditions that mimic the gut (See Supporting Information S1). Furthermore, MED6-189, GB209-2 and GB209-3 demonstrated no hemolytic activity at concentrations up to 10 µM (See Supporting information S1 and fig. S7, S8). To guide in vivo efficacy studies, we investigated the in vivo tolerability profile of MED6-189. Our data showed that animals treated with MED6-189 at doses up to 50 mg/kg did not exhibit any adverse events, and there were no significant changes in hematology, clinical chemistry analyses, nor necropsy observations. Importantly, the data also indicated improved exposure levels and compound absorption.

### *In vivo* MED6-189 efficacy in a humanized mouse model of *P. falciparum* malaria

Due to the highly favorable safety and tolerability profile exhibited by MED6-189, we chose a dosage of 50 mg/kg to assess the in vivo efficacy of the compound using a humanized mouse model of *P. falciparum* malaria (*57*). NOD scid gamma (NSG) mice were intravenously engrafted with human red blood cells (hRBCs) daily until the percentage of hRBCs reached ∼50%, at which point the mice were infected with 2×10^7^ *P. falciparum*-infected erythrocytes. At the third day post-infection, mice were administered either the vehicle alone or MED6-189 at 50 mg/kg orally, once a day for four days. Whereas the control group showed a rapid increase in parasite load with parasitemia reaching ∼6% by day 6 post-infection, mice treated with MED6-189 showed a rapid clearance of parasites achieving more than 90% reduction in parasitemia in the peripheral blood by the seventh day post-treatment (Fig. 5A, table S9).

**Figure 5.**
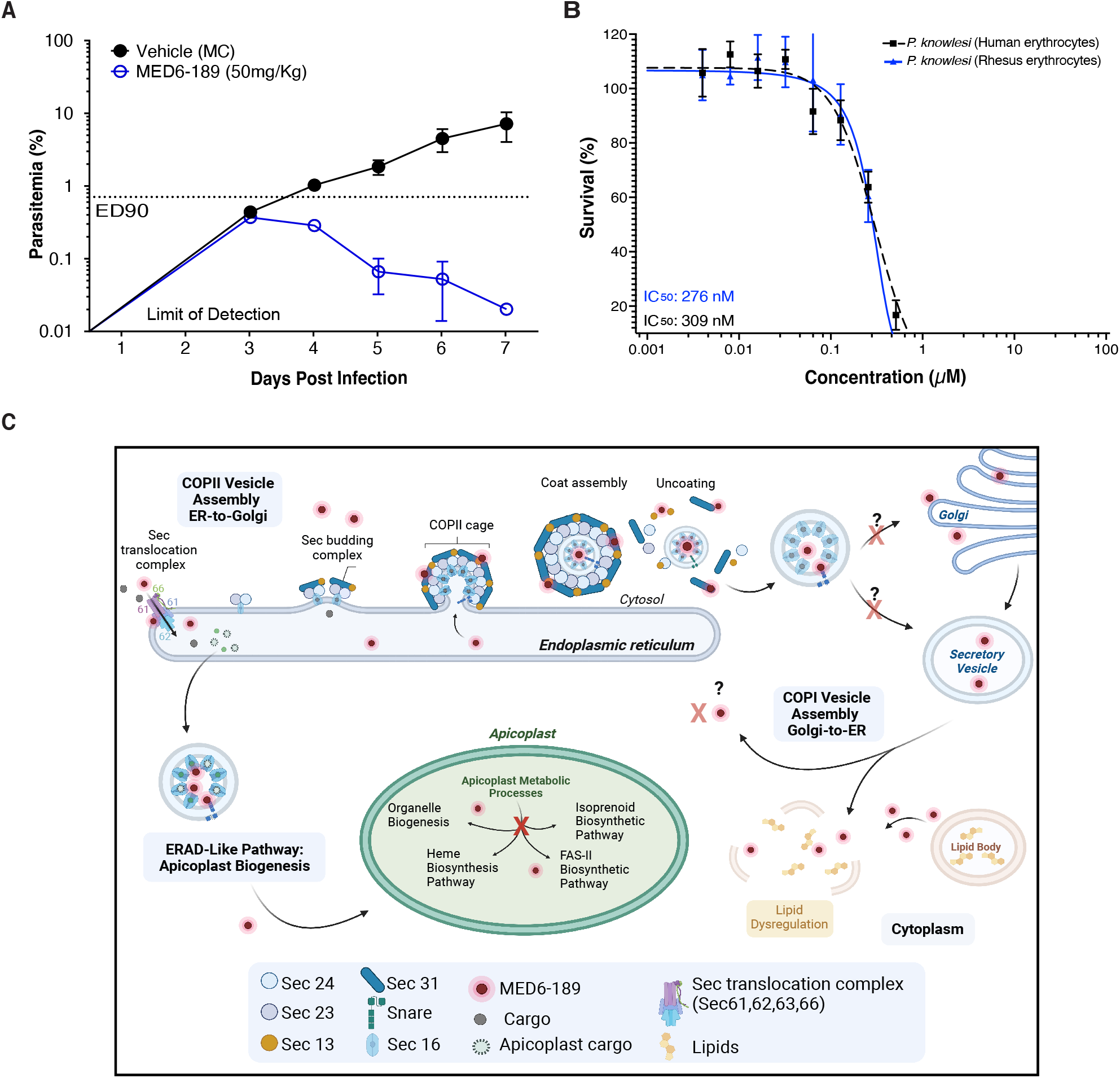
In vivo and broad-spectrum antimalarial efficacy of MED6-189. **A.** The in vivo efficacy of MED6-189 was evaluated in a humanized mouse model infected with *P. falciparum* (blue) compared to untreated controls (black). **B.** Dose-dependent response of MED6-189 on *P. knowlesi* YH1 human erythrocyte infecting (black) and rhesus erythrocyte infecting (blue) parasites. The graphs illustrate the logarithmic growth of parasites (Y-axis) in response to varying drug concentrations (X-axis). Error bars represent standard deviations from two independent experiments conducted in triplicate. The regression line is derived from a nonlinear regression analysis (Variable slope with four parameters, least squares fit). **C.** Proposed mode of action of MED6-189 in *P. falciparum*-infected erythrocytes. The compound is imported into the endoplasmic reticulum (ER) via the Sec translocation complex (SEC61,62,63,66), where it interacts with components of the ER transport machinery. The compounds translocated into the apicoplast where it directly interacts with proteins involved in crucial apicoplast function, ultimately disrupting this vital organelle. The interactions of MED6-189 with components of the apicoplast function and trafficking systems lead to dysregulation of lipids, resulting in the disruption of key biological processes within the *Plasmodium* parasite.

### Pan-antimalarial activity of MED6-189

Given the favorable biological activity of MED6-189 against *P. falciparum*, we also assessed its effectiveness against other *Plasmodium* species that infect humans, including *P. vivax*, the second most prevalent cause of human malaria (*1*). Since *P. vivax* cannot be propagated continuously in vitro, we examined the efficacy of MED6-189 against *P. knowlesi* and *P. cynomolgi*, which infect human erythrocytes and are commonly used as model systems for *P. vivax* infection (*58*). *P. knowlesi*, an Asian Old World monkey parasite with a robust in vitro culture system, is known to be zoonotic for humans. *P. cynomolgi*, a simian parasite capable of infecting humans experimentally (*54*), is phylogenetically closely related to *P. vivax* and shares similar biological and genetic properties with *P. vivax* (*58*). MED6-189 demonstrated significant activity against both parasites, with IC_50_ values of 309 nM ± 42 nM and 276 nM ± 23 nM against *P. knowlesi* in human and rhesus erythrocytes, respectively and (136 ± 56 nM, n=3) against *P. cynomolgi* (Fig. 5B and S9-S10) (Sigmoidal, 4PL, X is concentration, n=3, nonlinear regression, CI:95%). Together, the data validate MED6-189 as a potent antimalarial with a broad-spectrum activity that targets both *falciparum* and non-*falciparum* human malaria.

## DISCUSSION

Using a multifaceted approach, we have investigated the in vitro and in vivo efficacy, mode of action, resistance mechanisms, and preclinical safety profile of the kalihinol analogue MED6-189. Our findings establish MED6-189 as a promising lead in the fight against malaria, and it may serve as a representative of other kalihinol and ICT compounds with antiparasitic potential. Our studies further highlighted the pivotal role of the apicoplast in the biological activity of the drug. By employing targeted and untargeted analyses, we have confirmed the apicoplast as a major target of this drug’s action. Furthermore, our investigations have unveiled a slow-acting mode of operation for MED6-189, reminiscent of other small molecules that target the integrity and biogenesis of the apicoplast.

In order to localize MED6-189 within the parasites, we used a fluorescent analogue, which confirmed the apicoplast as a major site of MED6-189 accumulation. Consistent with this finding, the compound was found to have a slow killing mode of action, reminiscent of that of several classes of small molecules that target apicoplast integrity and biogenesis(*24, 25, 59*). Moreover, in vitro efficacy studies show that the activity of MED6-189 was antagonistic to that of fosmidomycin, a drug previously shown to disrupt the MEP pathway in the apicoplast. Interestingly, metabolic supplementation with IPP led to a partial rescue of parasite replication. Unlike fosmidomycin, MED6-189-treated parasites did not progress past the third intraerythrocytic cycle, suggesting that the compound may have multiple mechanisms of action, which may also account for the challenges in obtaining resistant parasites through standard drug selection methods.

Our metabolite profiling identified significant alterations in lysophosphatidylcholine (LPC), several triglycerides/triacylglycerides (TG/TAGs) and a limited subset of phosphatidylcholine (PC) forms in MED6-189-treated parasites. Notably, LPC exhibited the most pronounced change, with an estimated 3.5-fold increase in its steady-state levels. LPC, a precursor for the biosynthesis of PC, the most abundant phospholipid (50% of phospholipids in *P. falciparum* membranes), plays a key role in the regulation of sexual differentiation (*60*). Our data also indicated a significant reduction in the biosynthesis of sphingomyelin (*39*), sterol-esters and phosphatidylethanolamine (PE) in MED6-189-treated samples, suggesting a significant disruption in lipid metabolism and turnover caused by the compound. Our metabolomic analysis identified thiamine pyrophosphate (TPP) as a major altered metabolite in *P. falciparum* following treatment with MED6-189. TPP is essential for the activity of several enzymes and biosynthetic pathways, including the pyruvate dehydrogenase PDH complex found in the apicoplast (*38*). The apicoplast PDH converts pyruvate into acetyl-CoA, the major fatty acid precursor, whereas a second distinct PDH fuels the tricarboxylic acid cycle in the mitochondria (*61*). Altogether, our data suggest that MED6-189 operates by modulating *P. falciparum* lipid metabolism and apicoplast biogenesis (*62*).

Proteomic analyses further supported the link between MED6-189 activity and *P. falciparum* membrane biogenesis and trafficking pathways. Several components of the SEC and SNARE secretory machinery, namely SEC13 (PF3D7_1230700), SEC62 (PF3D7_1438100), SEC66 (PF3D7_0204200), and syntaxin SYN13 (PF3D7_1104100), were found either to be enriched in the drug pull-down experiments and/or showed differential melting profiles between MED6-189 treated versus vehicle-treated control samples. However, it is worth noting that not all targets identified in the pull-down assays were also identified by CETSA. With relatively fewer proteins of significance found stabilized in our CETSA dataset, one may hypothesize that MED6-189 may have greater impact on the ribosomal RNA, which is not captured in CETSA results. Other factors can account for the differences seen between CETSA and affinity proteomics findings including the level of abundance of specific proteins. For example, the SEC13 protein melting curves were not identified in our DMSO control samples yet found to be stabilized in our MED6-189 samples. The problem with low abundance target proteins not generating CETSA curves has been noted previously by Mateus and coworkers (*63*). The melting curves of SEC13 in the presence of DMSO and pyrimethamine have been previously established by Dziekan *et al* 2020 (*47*). Comparing these two sets of data seems to indicate that MED6-189 does indeed stabilize SEC13 beyond its previously observed melting temperature (fig. S6G).

SEC13 plays a major role in COPII-mediated vesicular transport between the nuclear pore complex, ER and Golgi membranes, whereas SEC62 and SEC66 (SEC71) are important components of protein translocation machinery in the ER *(44, 54, 55, 64, 65)*. The ER-associated degradation system (ERAD) is a quality control mechanism that retro-translocates misfolded secretory proteins across the ER. A similar system, the ERAD-like system, is believed to play a critical role in protein transport into the apicoplast(*66, 67*). Both the thermal proteomic profiling and pull-down assays identified significant interactions between MED6-189 and the SNARE protein family. Interestingly, syntaxin SYN13, a Qa-SNARE family protein, was also identified in our CETSA assay as *P. falciparum* proteins specifically stabilized by MED6-189. The role of these proteins in COPI and COPII vesicle trafficking has previously been reported (*68–71*). These proteins play a key role in membrane identification and mediate membrane fusions across various organelles including the endoplasmic reticulum, mitochondria, apicoplast, and other double-membrane-bound vesicles(*69, 72–74*). Most recently, SNARE proteins have been shown to play an important role in the biogenesis and maintenance of the apicoplast organelle in *Toxoplasma gondii* (*75*).

The thermal shift assay revealed that MED6-189 significantly affected the stabilization of proteins such as 2C-methyl-d-erythritol 2,4-cyclodiphosphate enzyme (MECP | PF3D7_0209300), ssDNA binding protein (SSB | PF3D7_0508800), and acetyl-coA binding proteins (PF3D7_0810000), all of which are integral to the biosynthesis of terpenoids(*76*), fatty acids, and apicoplast biogenesis systems (*66, 67, 76*).

Utilizing omics-based strategies, we have gained an impartial understanding of how MED6-189 is trafficked into the apicoplast and disrupts its biogenesis, ultimately leading to parasite cell death (Fig. 5C). These studies were further complemented by genetic approaches aimed at selecting parasite clones with reduced susceptibility to MED6-189. Unlike other antimalarials for which resistant parasites can be selected within a small number of intraerythrocytic cycles, parasites with enhanced tolerance to MED6-189 required an estimated 36 consecutive months of drug pressure. The IC_50_ for MED6-189 in these resistant parasites was determined to be 3-4-fold that of the parent 3D7 strain. Whole genome sequencing of the resistant clones identified SNP and Indel in *PfSec13* gene involved in vesicular trafficking. The importance of *SEC13* in MED6-189 cellular response was further validated using yeast as a model system. This finding led us to apply CRIPSR-cas-9 genetic editing tool in *P. falciparum* to validate the role of SEC13 in MED6-189 tolerance. *PfSec13-mut* lines were subsequently cloned and survival assays confirmed the role of SEC13 as an adaptive mechanism of resistance to MED6-189 treatment. Our findings through proteomics analyses, which suggest a direct interaction between SEC13 and MED6-189, indicate that both the mode of action and mechanism of resistance of the drug are most likely linked.

Altogether, our studies provide a comprehensive investigation into the activity and mode of action of MED6-189, a highly effective antimalarial compound targeting both the asexual and sexual stages of *P. falciparum*. The array of complex molecular pathways hindered by MED6-189 and presumably related ICT antimalarials along with the difficulty for the parasite to successfully develop resistance, makes it an effective candidate at preventing parasite progression and hindering transmission activity. The ability of MED6-189 to inhibit the growth of *P. falciparum in vitro* and in humanized mice as well as to inhibit the growth of *P. knowlesi* and *P. cynomolgi* makes it a promising pan-antimalarial lead drug.

## MATERIALS AND METHODS SUMMARY

Detailed information on materials and methods is available as Supplementary Materials and Methods. In brief, the anti-malarial activity of individual analogues were evaluated *in-vitro* against *P. falciparum* 3D7 (MRA-102, drug sensitive) W2 (MRA-157, Chloroquine resistant) strains (ATCC^®^ Manassas, VA), Dd2 (MRA-156), NF54 (MRA-1000), HB3 (MRA-155), and D10-Acp-GFP (MRA-568). *P. knowlesi* assays were quantitatively measured by flow cytometry using SYBR Green I, and Mitrotracker Deep Red as previously described (*77, 78*) on a Beckman Coulter CytoFLEX. Nonhuman primate infections were required to generate *P. cynomolgi* M/B strain parasites for in vitro testing. **RNA-sequencing:** differential expression analysis was done by use of R package DESeq2 with an adjusted P-value cutoff of 0.05. Volcano plots were made using R package Enhanced Volcano or GraphPad Prism 9 (GraphPad Software, Inc.). **LC-MS Metabolomics-lipids:** LC-MS metabolomics analysis was performed on a Synapt G2-Si quadrupole time-of-flight mass spectrometer (Waters) coupled to an I-class UPLC system (Waters). **LC-MS Metabolomics-polar metabolites:** Targeted metabolomics of polar, primary metabolites was performed on a TQ-XS triple quadrupole mass spectrometer (Waters) coupled to an I-class UPLC system (Waters). **Metabolomic data processing and analysis:** Untargeted data processing (peak picking, alignment, deconvolution, integration, and spectral matching) was performed in Progenesis Qi software (Nonlinear Dynamics). Data were normalized to total ion count. Similar metabolites features were assigned a cluster ID using RAMClust (*79*). An extension of the metabolomics standard initiative guidelines was used to assign annotation level confidence(*80, 81*). Targeted data processing was performed in Skyline software(*82*). **CETSA mass spectrometry data** was analyzed using mineCETSA R-language package as described previously (*47*). **Proteomics data processing and analysis:** Tandem mass (MS/MS) spectra were interpreted using ProluCID v.1.3.3(*83*). DTASelect v.1.9(*84*) and swallow v.0.0.1, an in-house developed software (*85*) were used to control FDRs at less than 1.2%. All datasets were contrasted against their merged data set, respectively, using Contrast v1.9(*84*) and in-house developed sandmartin. An in-house developed software, NSAF7 v.0.0.1, was used to generate spectral count-based label free quantitation results (*86*). QSPEC/QPROT (*40*) was used to calculate log_2_ Fold-Changes and Z-statistics. **Yeast strain Sec13p expression analysis:** Overexpression of *S. cerevisiae* Sec13p, was generated by an episomal *E. coli*/yeast shuttle vector, transformed into the BY4741 yeast strain. *SEC13* gene repression utilized the tet-off system. **Tolerability studies**: used Male CD1 (20-22g Envigo). **Toxicological Profiles:** Blood samples for analysis were carried out in the Abacus5 junior Vet → (Practice CVM S.L.L). Clinical chemistry was performed by mean analysis of whole blood in Vetscan→ (ABAXIS) & F560 (Menarini→) analyzer and by mean analysis of plasma. **In vivo Efficacy studies:** MED6-189 oral efficacy was tested in female NOD-*scid IL-2Rγ^null^* (NSG) mice engrafted with human erythrocytes and infected *P. falciparum*-infected erythrocytes. Peripheral blood from *P. falciparum*-infected mice were stained with TER-119-Phycoerythrine (marker of murine erythrocytes) and SYTO-16 (nucleic acid dye) and then analyzed by flow cytometry (FACSCalibur, BD). **Bioanalysis and pharmacokinetics analysis: LC-MS analysis:** An Acquity Ultra-Performance liquid chromatography (UPLC) system (Waters Corp., Milford, MA, USA) couple to a triple quadrupole mass spectrometer (API 4000™, AB Sciex, Foster City, CA, USA) was used for the analysis. **Pharmacokinetic analysis:** Blood concentration time data were analyzed by Non-Compartmental PK analysis using Phoenix WinNonlin software (Certara NY, US) to calculate PK parameters.

### Statistical analyses

Parasitemia and proportion of asexual stages were analyzed using a two-way ANOVA with Tukey’s test for multiple comparisons. Significant differences were indicated as following: * for p< 0.05; ** for p< 0.01, *** for p<0.001 and **** for p<0.0001. Statistical tests were performed with GraphPad Prism. Figures were generated with GraphPad Prism 9, Biorender, Adobe illustratorv27.3.1. Putative Sec13 protein was created by Alphafold monomer V2.0 prediction for protein transport protein SEC13 (*Q8I5B3*) with PDB reference *AF-Q8I5B3-F1-model_v4 (1).pdb* and protein structure was formed through ChimeraX.

### Funding

This work was supported by the National Institutes of Allergy and Infectious Diseases of the National Institutes of Health (Grant R01 AI-138139 to C.D.V., C.B.M, and K.L.R) and the University of California, Riverside (NIFA-NIFAHatch-225935 to K.G.L.R.). C.B.M. research is supported by NIH grants AI138139, AI123321, AI152220 and AI153100 and AI136118; the Steven and Alexandra Cohen Foundation (Lyme 62 2020); the Global Lyme Alliance and The Blavatnik Fund for Innovation. A.K.B. is supported by an International Research Scientist Development Award (K01 TW010496) from the Fogarty International Center of the National Institutes of Health. *P. k.* studies were supported by the Emory National Primate Research Center (Grant No. ORIP/OD P51OD011132) and Emory National Primate Research Center (Grant No. U42PDP11023). This publication includes data generated at the UC San Diego IGM Genomics Center utilizing an Illumina NovaSeq 6000 that was purchased with funding from a National Institutes of Health SIG grant (#S10 OD026929). The additional support of Tres Cantos Open Lab Foundation is gratefully acknowledged.

### Author Contributions

K.L.R., C.B.M., and C.D.V. conceived and designed all experiments. Z.C. performed all experiments with *P. falciparum* and S.A. contributed to the bioinformatics data analyses. M.E.D. and J.H.C. synthesized the kalihinol derivatives, including the biotinylated and fluorescent analogues. G.L.B. performed hydrolytic stability studies on MED6-189 and synthesized and characterized the formamide hydrolysis products. J.S.K. performed the metabolomic experiments while A.S., C.B and L.F. performed the proteomics mass-spectrometry analyses. J.Y.C. conducted the studies in yeast. I.R., M.C.C. and S.V-M. evaluated the in vivo efficacy of MED6-189 and MED6-159 in murine malaria models. I.C., P. C., J. D. M-A., N. I. and A. G-P. evaluated the safety, pharmacokinetic and tolerability profiles of MED6-189. A.K.B. evaluated the in vitro efficacy of MED6-189 and MED6-159 in *P. knowlesi* malaria parasites. C.J.J and M.A.A conducted the studies in *P. cynomolgi.* Z.C, K.L.R., C.B.M., C.D.V., J.C. and P.V. contributed to the writing of the manuscript. All authors reviewed, edited, and approved the final manuscript.

### Data availability

The MED6-189 purifications and CETSA mass spectrometry datasets have been deposited to the ProteomeXChange (with PXD038457 and PXD038053 accessions, respectively) via the MassIVE repository (MSV000090812 and MSV000090667; with password: kalihinol) and may also be accessed from the Stowers Original Data Repository (http://www.stowers.org/research/-publications/libpb-1759). WGS and RNA-seq datasets generated in this study have been deposited in the NCBI BioProject database under SubmissionID: SUB12241156 with BioProject ID: PRJNA930408.Metabolomics datasets generated in this study are available at Panorama database and can be accessed through the following link (https://panoramaweb.org/qpjX5B.url)usingaccession codes.

### Code availability

The entire in-house software suite (Kite) used for the MudPIT mass spectrometry analysis is available in Zenodo: (https://zenodo.org/record/5914885#.Y9q0juzMKjg).

## Acknowledgements

We thank Matthias Göhl and Ryan Kozlowski for early efforts to scale up the synthesis of MED6-189. Alphafold monomer V2.0 prediction for protein transport protein SEC13 (Q8I5B3) and protein structure was formed through ChimeraX.

