## Supplemental Figures for "A Potent Kalihinol Analogue Disrupts Apicoplast Function and Vesicular Trafficking in *P. falciparum* Malaria": MED6-189_Supplemental Figures_11_21_23.pdf

**The PDF file includes:**

- Materials and Methods
- Supplementary Figures
  - Figs. S1 to S10

**Other Supplementary Materials for this manuscript include the following:**

- Supplementary Tables
  - Tables S1-S9
- Supporting Information S1
- Supporting Information S2

### Materials and Methods

**Asexual parasite Culture and maintenance--** *P. falciparum* 3D7 (MRA-102) parasites were propagated in 5% of human $O^+$ erythrocytes and 10 mL of RPMI-1640 medium containing 0.5% Albumax II (Invitrogen), 2 mM L-glutamine, 50 mg/L hypoxanthine, 25 mM HEPES, 0.225% $NaHCO_3$ and 10 mg/mL gentamycin. They were maintained at 37 °C and gassed with a sterile mixture of 5% $O_2$, 5% $CO_2$ and 90% N2 (*75, 76*). Parasite synchronization was achieved using two 5% D-sorbitol treatments 8 hours apart (*77*).

*P. knowlesi* strain YH1 (provided courtesy of Manoj Duraisingh) parasites were routinely cultured in 4% rhesus erythrocytes (Yerkes primate research center) and 10 mL of RPMI-1640 medium containing 0.5% Albumax II (Invitrogen), 2 mM L-glutamine, 50 mg/L hypoxanthine, 25 mM HEPES, 0.225% $NaHCO_3$ and 10 mg/mL gentamycin. They were maintained at 37 °C and gassed with a sterile mixture of 1% $O_2$, 5% $CO_2$ and 94% N2. Parasite synchronization was achieved using two 5% D-sorbitol treatments, spaced one per re-invasion cycle, 24 hours apart.

**Animal use--** Nonhuman primate infections were required to generate *P. cynomolgi* M/B strain parasites for in vitro testing because these parasites are not adapted to continuous culture. Infections were performed at the University of Georgia, an Association for Assessment and Accreditation of Laboratory Animal Care (AAALAC) accredited institution. The UGA Institutional Animal Care and Use Committee reviewed and approved all procedures including blood collections, infections with malaria parasites, clinical interventions, etc. During the experimental procedures, the animals were housed in compliance with Animal Welfare Act regulations as well as the Guide for the Care and Use of Laboratory Animals. This includes multiple enrichment activities provided daily, including physical manipulanda and environmental enrichment, and routine health checks performed by facility animal resources staff and veterinarians.

***P. cynomolgi* M/B strain parasite generation and cryopreservation--** Captive bred Japanese macaques (*Macaca fuscata*) were obtained from Oregon National Primate Research Center. These animals were infected with *P. cynomolgi* M/B strain blood stage parasites that were cryopreserved. After infection, parasitemia was monitored daily by thick and thin blood films as previously described in Joyner *et al.* 2019 (*78*). When parasitemia reached about 1-3% ring stages, blood was collected from the infected animal and cryopreserved in Glycerolyte 57 solution as previously described (*79*).

***P. cynomolgi* M/B strain Drug Assay--** *P. cynomolgi* M/B strain parasites preserved in Glycerolyte 57 were thawed as previously described (*79*). After thawing, the parasitemia was enumerated and adjusted to approximately 1% rings by diluting with freshly isolated Japanese macaques RBCs. The 1% parasitemia mixture was then plated into a 96 well round bottom culture plate at 5% hematocrit. The culture medium was comprised of IMDM supplemented with GlutaMAX (Gibco #31980030) containing 30 mM HEPES (Sigma-Aldrich), 17 mM D-glucose (Fisher Chemical) and 440 µM hypoxanthine (Sigma-Aldrich) and 1% Albumax II (Gibco) and 20% (v/v) heat-inactivated rhesus or Japanese macaque serum. Med6-189 was then added to each well to obtain the needed final concentrations of drug. After plating the parasites were placed into a StemCell hypoxia chamber with a gas mixture of 95% $N_2$ and 5% $CO_2$. Cultures were then placed into a CellExpert Eppendorf Incubator and maintained at 37°C and samples harvested at the indicated time points. Culture media was replaced daily, and drug re-added as described above to ensure that fresh compound was always present. During the drug assay thin blood smears were made and stained with Giemsa to determine the phenotypic changes with the parasites after exposure to the compound. For determining the $IC_{50,}$ the parasites were harvested at 60 hours after the initial drug dosing. To determine the parasitemia in each well, the wells were resuspended in 30 ul of 10 µg/ml of Hoechst 33342 and incubated at for 30 minutes at 37°C. The stained culture wells were pelleted by centrifugation and resuspended in PBS followed by analysis on a Novocyte Quanteon Flow Cytometer. The experiment was repeated twice, and each concentration was assessed in triplicate.

**Data analysis--** The percentage of parasitized erythrocytes was determined by enumerating the number of Hoechst positive events out of the total erythrocytes after excluding non-single cells by forward and side scatter. The percent survival was calculated by averaging the parasitemia in the triplicates for each condition and then dividing by the parasitemia observed in the wells that did not receive drug treatment. The $IC_{50}$ was then calculated using GraphPad Prism 9 (GraphPad Software, Inc.).

**Localization of fluorescently labeled MED6-189 analogue--** 3D7, NF54 and D10-Acp-GFP parasite lines were washed in incomplete medium then incubated with 1 µM of JF549-labelled kalihinol analogue (MED6-131) for 30 min-2 hrs. at 37°C, were washed in incomplete medium and fixed onto coverslip with 4% paraformaldehyde for 20 min at RT under darkness. After 3-5 PBS washing, coverslips were mounted in Vectashield Antifade Mounting Medium with DAPI (H-1200). Images were acquired using Keyence BZ-X810 and processed with ImageJ program.

**Parasitemia and asexual stage phenotype analyses--** To understand the phenotypic effects of MED6-189 on *P. falciparum*, 3D7 lines were synchronized, grown for one cycle post-synchronization, and treated with either the vehicle (DMSO) or MED6-189 at $IC_{80}$ values for 48 hrs. Culture media were replaced daily with reintroduction of the drug. Parasitemia and proportion of the different asexual blood stages were determined by counting Giemsa-stained blood smears by light microscopy at every 12-hour time point.

**Parasitemia and sexual stage phenotype analyses--** To understand the phenotypic effects of MED6-189 on *P. falciparum* sexual stage development, NF54 lines were synchronized, and exposed to stress induced gametocyte induction followed by treatment with *N*-acetyl glucosamine (NAG; Sigma-Aldrich) supplemented media for 6-7 days to eliminate remaining asexual parasites. Upon first signs of gametocyte formation observed through giemsa stained smears (5-7 post stress induction), samples were treated with either the vehicle (DMSO) or MED6-189 at 3x $IC_{80}$ values with 3 biological replicates per condition. Culture media were replaced daily with reintroduction of the drug. Gametocyte parasitemia and proportion of the different sexual blood stages were determined by counting Giemsa-stained blood smears by light microscopy at every 24-hour time point (*80*).

***P. falciparum* in vitro SYBR green fluorescence-based assays--** The anti-malarial activity of individual analogues were evaluated *in-vitro* against *P. falciparum* 3D7 (MRA-102, drug sensitive) and W2 (MRA-157, Chloroquine resistant) strains (ATCC® Manassas, VA), Dd2 (MRA-156), NF54 (MRA-1000), HB3 (MRA-155), and D10-Acp-GFP (MRA-568) via SYBR green-I based fluorescence assay. Parasite growth rate and stages of development were determined via microscopy through Giemsa-stained smears of the cultures. Serial dilutions of the compounds ranging from 10uM-0.5nM were prepared in 96-well plates (Corning, Costar 3904) followed by the introduction of synchronized ring stage parasites cultures at 0.5% parasitemia and 2.5%

hematocrit added to each well. Eight wells were treated with vehicle control (DMSO) as control. Plates were incubated in a modular incubation chamber (Billups-Rothenberg, Del Mar, CA) maintained at 37 °C for 72 hrs. in a low oxygen environment (96% $N_2$, 3% $CO_2$, 1% $O_2$). After 72 h, plates were removed from incubation and stored at -80˚C for 24 hrs. Plates were subsequently thawed followed by incubation with 1X volume of lytic buffer (20 mM Tris-HCl, pH 7.5, 5mM EDTA, 0.008% Saponin, 0.08% Triton X-100) containing SYBR Green 1X for 4–6 h at 37˚C in darkness. Plates were read with a Molecular Devices SpectraMAX Gemini EM at Ex. 495 nm, Em. 525 nm. Assessment of anti-malarial activity of compounds was made on the basis of fifty percent inhibitory concentration values ($IC_{50}$) determined by DNA content of the parasite using SigmaPlot (10 (Systat)) or GraphPad Prism v 9.0.0 under program Sigmoidal, 4PL, X is concentration, nonlinear regression, CI:95%)(*75-77*).

***P. knowlesi* In vitro $IC_{50}$ Assays--** Parasite lines were double synchronized with 5% D-sorbitol, grown for one cycle post-synchronization, and treated with a range of concentrations of either the vehicle (DMSO) or MED6-189, plated in triplicate. *P. knowlesi* parasites were allowed to re-invade (24 hours post-assay) and assays were harvested at 36 hours post-reinvasion. Parasitemia was quantitatively measured by flow cytometry using SYBR Green I, and Mitrotracker Deep Red as previously described (*81, 82*) on a Beckman Coulter CytoFLEX (fig. S9). Data from 100,000 erythrocytes were acquired per sample and the percentage of double positive SYBR Green I positive, viable parasitized cells was calculated as parasitemia.

**RNA-sequencing--** Two independent biological replicates were generated for each time point at the ring, trophozoite or schizont stage, for every culture condition and line. Total RNA was extracted with TRIzol® LS Reagent (Invitrogen) followed by incubation for 1 hr. with 4 units of DNase I (NEB)at 37°C. RNA samples were visualized by RNA electrophoresis and quantified on Synergy™ HT (BioTek). mRNAs were then purified using NEBNext® Poly(A) mRNA Magnetic Isolation Module (NEB) according to the manufacturer's instructions. Libraries were prepared using NEBNext® Ultra™ Directional RNA Library Prep Kit (NEB) and amplified by PCR with KAPA HiFi HotStart Ready Mix (KAPA Biosystems). PCR conditions consisted of 15 min at 37°C followed by 12 cycles of [98°C (30 s), 55°C (10 s) and (62°C 1 min 15)] and 5 min at 62°C. The quantity and quality of the final libraries were assessed using a Bioanalyzer (Agilent Technology Inc). Libraries were sequenced using a NovaSeq 6000 DNA sequencer (Illumina), producing paired-end 100-bp reads. FastQC (https://www.bioinformatics.babraham.ac.uk/projects/fastqc/) was used to analyze raw read quality. Based on the quality data, the first 11 bp of each read and any adapter sequences were removed using Trimmomatic (http://www.usadellab.org/cms/?page=trimmomatic). Bases were trimmed from reads using Sickle with a Phred quality threshold of 25 (https://github.com/najoshi/sickle) and reads shorter than 18 bp were removed. The resulting reads were mapped against the *P. falciparum* 3D7 genome (v46) using HISAT2 (version 2-2.2.1) with default parameters. Uniquely mapped, properly paired reads with mapping quality of 40 or higher were retained using SAMtools (http://samtools.sourceforge.net/). PCR duplicates were removed using PicardTools MarkDuplicates (Broad Institute). Genome browser tracks were generated and viewed using the Integrative Genomic Viewer (IGV) (Broad Institute). Raw read counts were determined for each gene in the *P. falciparum* genome using BedTools (https://bedtools.readthedocs.io/en/latest/#) to intersect the aligned reads with the genome annotation. Differential expression analysis was done by use of R package DESeq2 to call up- and down-regulated genes with an adjusted P-value cutoff of 0.05. Volcano plots were made using R package Enhanced Volcano or GraphPad Prism 9 (GraphPad Software, Inc.).

**Metabolomic profiling: Sample Preparation--** Tightly synchronized parasites ($9 \times 10^8$ parasites at 72hrs post synchronization) were lysed with saponin, flash frozen and stored at −80 °C in triplicate. Lipids and polar metabolites were extracted from parasite pellets using a biphasic approach. To each sample, 1 mL of ice cold 3:2 methyl tert-butyl ether:80% methanol was added. To break up parasite pellets, samples were vortexed 2 min, sonicated for 15 min, vortexed for 2 min, sonicated for 15 min, then vortexed for 30 min at 4° C. All sonication was performed in an ice bath. 200 µL of water was added to induce phase separation followed by a 5 min vortex. After centrifugation for 15 min at 4° C at 16,000 x g, 200 µL of the top, nonpolar layer was transferred to a 2 mL glass vial and the bottom, polar layer, was transferred to a new 2 mL glass vial then analyzed by LC-MS. The nonpolar fraction was dried under a gentle stream of nitrogen at room temperature then resuspended in 400 µL of 9:1 methanol: toluene and analyzed by LC-MS.

**LC-MS Metabolomics-lipids--** LC-MS metabolomics analysis was performed on a Synapt G2-Si quadrupole time-of-flight mass spectrometer (Waters) coupled to an I-class UPLC system

(Waters). Separations were carried out on a CSH phenyl-hexyl column (2.1 x 100 mm, 1.7 μM) (Waters). The mobile phases were (A) water with 0.1% formic acid and (B) acetonitrile with 0.1% formic acid. The flow rate was 250 μL/min, and the column was held at 40° C. The injection volume was 1 μL. The gradient was as follows: 0 min, 1% B; 1 min, 1% B; 8 min, 40% B; 24 min, 100% B; 26.5 min, 100% B; 27 min, 1% B. The MS was operated in positive ion mode (50 to 1200 m/z) with a 100 ms scan time. Source and desolvation temperatures were 150° C and 600° C, respectively. Desolvation gas was set to 1100 L/hr and cone gas to 150 L/hr. All gases were nitrogen except the collision gas, which was argon. Capillary voltage was 1 kV in positive ion mode. A quality control sample, generated by pooling equal aliquots of each sample, was analyzed every 3 injections to monitor system stability and performance. Samples were analyzed in random order. Leucine enkephalin was infused and used for mass correction.

**LC-MS Metabolomics-polar metabolites--** Targeted metabolomics of polar, primary metabolites was performed on a TQ-XS triple quadrupole mass spectrometer (Waters) coupled to an I-class UPLC system (Waters). Separations were carried out on a ZIC-pHILIC column (2.1 x 150 mm, 5 μM) (EMD Millipore). The mobile phases were (A) water with 15 mM ammonium bicarbonate adjusted to pH 9.6 with ammonium hydroxide and (B) acetonitrile. The flow rate was 200 μL/min, and the column was held at 50° C. The injection volume was 2 μL. The gradient was as follows: 0 min, 90% B; 1.5 min, 90% B; 16 min, 20% B; 18 min, 20% B; 20 min, 90% B; 28 min, 90% B. The MS was operated in selected reaction monitoring mode. Source and desolvation temperatures were 150° C and 600° C, respectively. Desolvation gas was set to 1100 L/hr and cone gas to 150 L/hr. Collision gas was set to 0.15 mL/min. All gases were nitrogen except the collision gas, which was argon. Capillary voltage was 1 kV in positive ion mode and 2 kV in negative ion mode. System stability was monitored by analyzing a quality control sample (generated by pooling together equal volumes of all sample extracts) every 3 injections. Samples were analyzed in random order.

**Metabolomic data processing and analysis--** Untargeted data processing (peak picking, alignment, deconvolution, integration, and spectral matching) was performed in Progenesis Qi software (Nonlinear Dynamics). Data were normalized to total ion count. To group features belonging to the same metabolite, features were assigned a cluster ID using RAMClust (*83*). An extension of the metabolomics standard initiative guidelines was used to assign annotation level confidence(*84, 85*). Several metabolite databases were screened including Metlin, Lipidblast, Mass Bank of North America, and an in-house database. Targeted data processing (manual peak integration) was performed in Skyline software(*86*).

**Preparation of Cellular Extract Thermal Shift Assay (CETSA) samples for mass spectrometry analysis--** Acetone precipitated protein samples were resuspended in 100 µl of 100 mM triethylammonium bicarbonate (TEAB) and digested overnight at 37°C with 2.5 µg trypsin (Promega, Madison, WI). After digestion, approximately 0.5 *p*moles of LysC/trypsin digested porcine albumin was added to each sample as an internal control to monitor sample integrity. Peptide concentration was measured using the Pierce quantitative fluorometric peptide assay (Thermo Scientific, San Jose, CA). A sample volume equivalent to 30 µg in the lowest temperature sample (37°C) was used for subsequent labeling with TMT10plex reagents (Thermo Scientific, San Jose, CA). Samples were labeled with approximately 0.27 mg of TMT10plex reagent according to the manufacturer's instructions. Labels were matched to thermal challenge temperatures in order ($TMT^{10}$-126, 37°C to $TMT^{10}$-131, 73°C). Labeled samples for each treatment (DMSO, MED6-189) were pooled, buffer was removed using a vacuum centrifuge, and samples fractionated using the Pierce high pH fractionation kit according to the manufacturer's instructions. After fractionation, buffer was removed using a vacuum centrifuge and pellets resuspended in buffer A containing 95% HPLC grade water, 5% acetonitrile, and 0.1% formic acid (v/v/v), pH 2.6.

**Mass spectrometry analysis of CETSA samples--**TMT labeled samples were resolved for mass spectrometry analysis using a Dionex UltiMate 3000 RSLCnano liquid chromatography system. Peptides were first transferred from the autosampler to an Acclaim PepMap 100 C18 LC Trap Cartridge (inside diameter 0.3 mm, length 5 mm). Peptides were then resolved using an analytical column (75 µm inside diameter, 250 mm length) packed with 1.9 µM ReproSil®-Pur C18-aQ resin. Chromatographic separations used different combinations of buffer A (95% HPLC grade water, 5% acetonitrile, and 0.1% formic acid (v/v/v), pH 2.6), and buffer B (20% HPLC grade water, 80% acetonitrile, and 0.1% formic acid (v/v/v), pH 2.6). Chromatography steps were performed with a flow rate of 180 nl/minute as follows: (1) 2% B for 35 minutes (column equilibration); (2) a linear gradient from 2% B to 10% B over 3 minutes; (3) a linear gradient from

10% B to 40% B over 40 minutes; (4) a linear gradient from 40% B to 80% B over 10 minutes; (5) 80% B for 10 minutes (column wash); (6) a linear gradient from 80 % B to 2% B over 0.1 minutes; (7) 2% B for 12 minutes (column re-equilibration).

Peptides were analyzed using an Orbitrap Eclipse Tribrid mass spectrometer (Thermo Scientific, San Jose, CA) fitted with a FAIMS Pro™ interface. FAIMS compensation voltages (CV values) of 40 V and 60 V were used. For peptide identification and quantitation, an MS3 method was used: Full MS scans were performed using the Orbitrap mass analyzer (120,000 m/z resolution, 400-1600 m/z scan range); Peptides were isolated using a window of 0.7 m/z for MS2 fragmentation (35% CID energy) and subsequent detection with the ion trap mass analyzer. Synchronous precursor selection was used to select up to 10 peptide fragments identified during the MS2 step for MS3 analysis; Peptides for MS3 analysis were isolated using an MS1/MS2 isolation windows of 2 m/z; TMT reporter ions were removed from peptide fragments using HCD (collision energy 55%) and detected using the Orbitrap mass analyzer (50,000 m/z resolution, 100-500 m/z scan range).

The resulting .raw files for two technical replicates of each condition were processed using Proteome Discoverer 2.5 (Thermo Scientific, San Jose, CA) using the common workflow templates "PWF_Tribid_TMT_Quan_SPS_MS3_sequestHT_Percolator" (processing step) and "CWF_Comprehensive_Enhanced Annotation_Reporter_Quan" (consensus step). The "Normalization Mode" of the "Reporter Ions Quantifier" node in the consensus step was set to "none". Data were searched using the database published by Dziekan *et al* (*45*) with the addition of the sequence of porcine albumin.

**Downstream Analysis of CETSA mass spectrometry data--** Protein quantitation data were exported to .txt files (table S5) for analysis using the mineCETSA R-language package as described previously (*45*). Exported files were pre-processed to retain only high-confidence P. falciparum proteins for which abundance ratios had been generated. The resulting CETSA curves are provided as Supporting Information S2.

**Immunoprecipitation and MudPIT mass spectrometry--** Mid- to late-stage asexual parasites were collected following saponin treatment and purified. Samples were then resuspended in 50 mM-Tris-HCl pH 7.5, 300 mM NaCl, 0.5 mM EDTA, 0.5 mM EGTA, 2 mM AEBSF 0.5% Triton X-100, 20mM N-ethymaleimide and 1 mM AEBSF and EDTA-free protease inhibitor cocktail (*87*). Post cell lysis solution was homogenized via sonication for 6-9 rounds. The soluble extracts were centrifuged at 13,000g for 15 min at 4°C. The lysates were precleared with Dynabeads MyOne TM Streptavidin T1 (Invitrogen) for 1h at 4°C. 100uM of biotinylated kalihinol analogue (MED6-118) was added to each sample with the exception of DMSO added to a negative control and incubated for 1 hr at 4˚C. Following drug incubation Fresh Dynabeads MyOne TM Streptavidin T1 beads were added to each sample and incubated overnight at 4°C. Dynabeads were used to precipitate biotinylated drug-protein complexes and were washed with buffer (PBS, 0.05% Tween20). Proteins were eluted using 50mM Tris-HCl pH 6.7, 100mM DTT and 2% SDS. The eluent was subsequently precipitated on ice in 20% TCA overnight followed by cold acetone washes. The urea-denatured, reduced, alkylated, and digested proteins were analyzed by Multidimensional Protein Identification Technology (MudPIT) on a linear ion trap tandem mass spectrometer coupled to an Agilent 1100 series HPLC as described previously (*88*).

**Proteomics data processing and analysis--** Tandem mass (MS/MS) spectra were interpreted using ProluCID v.1.3.3(*89*) against a database consisting of 5527 non-redundant (NR) Plasmodium falciparum 3D7 proteins (PlasmoDB-42 release), 36661 NR human proteins (NCBI, 2018-03-30 release), 419 usual contaminants (human keratins, IgGs, and proteolytic enzymes). DTASelect v.1.9(*90*) and swallow v.0.0.1, an in-house developed software (https://github.com/tzw-wen/kite)(*91*) were used to control FDRs at less than 1.2%. All datasets were contrasted against their merged data set, respectively, using Contrast v1.9(*90*) and in-house developed sandmartin v.0.0.1 (https://github.com/tzw-wen/kite/tree/master/ kitelinux). Our in-house developed software, NSAF7 v.0.0.1 (https://github.com/ tzw-wen/kite/tree/master/windowsapp/NSAF7x64), was used to generate spectral count-based label free quantitation results(*92*). QSPEC/QPROT (*38*) was used to calculate $\log_2$ Fold-Changes and Z-statistics to compare five replicate MED6-189 affinity purifications to three negative controls. Proteins detected in at least 2 of the 5 replicate analyses and not in the controls and/or with a $\log_2$ FC ≥1.5 and Z-statistic ≥1.645 were considered significantly enriched in the MED6-189 affinity purifications.

**Generation of transgenic *P. falciparum PfSec13-mut* strains--**Gene editing of the *P. falciparum Sec13* gene *(PF3D7_1230700)* spanning position (Ch12:1258922 - 1262608 (+)) on chromosome

12 was performed using a two-plasmid-based strategy. The plasmid pCasG-Cas9-sgRNA plasmid vector (gifted by the Prigge lab contains the site to express the sgRNA, along with the yDHODH gene as the positive selection marker. The sgRNA was selected from the database generated by Desai *et al. (93)* and cloned into pCasG-Cas9 plasmid vector at the BsaI restriction site (table S8A, B). The homology directed repair plasmid (modified pDC2-donor-*bsd* without eGFP) was designed to remove the 21 nt target region (Pf3D7_12_v3:1261089-1261110) with PCR insert of interest through homology arms using a BamHI restriction enzyme digest followed by overnight ligation for donor insertion. Selection was performed though selectable markers, blasticidin S-deaminase (bsd)(*94, 95*) present in the pDC2-donor plasmid construct along with DSM-1, in which the resistance gene yDHODH is present in the pCASG plasmid. The 541bp Pf*Sec13* target specifying homology arm sequences were isolated through PCR amplification and gel purification using primer set A and B respectively amplified at 60°C and 54°C annealing temperatures respectively. The final donor vectors were confirmed by restriction digest and Sanger sequencing. Plasmids were isolated from 250 mL cultures of *Escherichia coli* (XL10-Gold Ultracompetent Cells, Agilent Cat. 200314) and 60 µg of each plasmid was used to transfect ring stage parasites. 24-hrs before transfection, mature parasite cultures (6-8% parasitemia) were magnetically separated using magnetic columns (MACS LD columns, Miltenyi Biotec) and diluted to 1% parasitemia containing 0.5 mL fresh erythrocytes (*96*). The next day, ~3% ring stage parasites were pelleted and washed in 4 mL of cytomix (*97*). 200 µl of the infected erythrocytes were resuspended with the two plasmids in cytomix to a total volume of 400 µl in a 0.2 cm cuvette. Electroporation was performed with a single pulse at 0.310 kV and 950 µF using the Biorad GenePulser electroporator. Cells were immediately transferred to a flask containing 12 mL media and 400 µl erythrocytes. The culture medium was exchanged five hours post electroporation with 12 mL of fresh media. The following day, fresh culture media were added and supplemented with 1.5 µM DSM-1 (Calbiochem) and 2.5 µg/mL Blasticidin (RPI Corp B12150-0.1). Media and drug selection was replenished every 24 hours for 7 consecutive days after which drug selection was halted and fresh media was replenished weekly. After 14 days, the culture was split into two flasks and 50 µl of erythrocytes were added every two weeks. Once parasites were detected by microscopy, integration of the insert was confirmed by gDNA extraction, PCR amplification of fragment regions upstream of 5’ arm and downstream of 3’arm found in plasmid construct using primer sets indicated in (Sup table 8b) and through genomic sequencing.

**Isolation of *PfSec13-mut* Clones**-- To generate a genetically homogenous parasite lines, the transfected parasites were serially diluted to approximately 0.5 parasite/well into 96 well plates. 200µl final volume of cultured parasites were incubated with Blasticidin S for 4 weeks with a weekly change of media and addition of fresh red blood cells during the first 2 weeks followed by media changes every 2 days until parasite recovery is observed through microscopic analysis of Giemsa-stained blood smears.

**Molecular analysis of the *PfSec13-mut* line--** Genomic DNA (gDNA) was extracted and purified using DNeasy Blood & Tissue kit (Qiagen) following instructions from the manufacturer. The diagnostic and genotyping PCR analysis was used to genotype the transfected lines using the primer pairs (Set A: Sec13-insert-fwd, Sec13-insert-rev; Set B:Sec13-pam-mut-fwd, Sec13-pam-mut-R). The PCR amplification was conducted using 2xKAPPA master mix for thirty cycles with an annealing temperature of 54°C and 60°C for Sec13 extension, respectively, and extension temperature of 57°C. The PCR amplicons were analyzed by gel electrophoresis and sequenced. For whole genome sequencing, genomic DNAs were fragmented using a Covaris S220 ultrasonicator and libraries were generated using KAPA LTP Library Preparation Kit (Roche, KK8230). To verify that the insertion was present in the genome at the correct location in both transfected lines, reads were mapped using Bowtie2 (version 2.4.4) to the *P. falciparum* 3D7 reference genome (v48, PlasmoDB), edited to include the insertion sequence in the intended location. IGV (Broad Institute) was used to verify that reads aligned to the insertion sequence.

**Analysis of MED6-189 effect on the growth of yeast strains with repressed or over-expressed Sec13p--** For overexpression of *S. cerevisiae* Sec13p, a DNA fragment consisting of 860 bp upstream, 894 bp *SEC13* coding sequence, and 297 bp 3' untranslated sequence was generated by PCR amplification and introduced into an episomal *E. coli*/yeast shuttle vector, pRS425, by using an in-fusion cloning kit (Takara Bio USA, Inc) and the resulting pRS425-SEC13 plasmid was transformed into the BY4741 yeast strain. To swiftly repress the *SEC13* gene, we utilized the tet-off system. This involved substituting the native *SEC13* gene promoter with a tetracycline (tet)-regulatable counterpart, while leaving the original *SEC13* open reading frame intact. The yeast strain with the *SEC13* gene under the control of Tet off promoter (BY4741 kanR-tetO7-TATA-SEC13, URA::CMV-tTA) was obtained from Horizon Discovery Ltd and the repression of the gene on the strain was achieved by the addition of doxycycline to the yeast culture. To determine the effect of MED6-189 on the growth of the yeast strains, the cells were grown in a synthetic minimal growth medium to mid log phase, diluted and inoculated in 96-well plates at the cell destiny of $10^3$ cells/mL in 100 µL culture volume also containing MED6-189 (12 sequential 2-fold dilutions at a range of 0 to 400 µM). Cell growth was monitored every 12 hours by measuring optical density at 600 nm using a BioTek SynergyMx microplate reader. Prizm software was used to analyze the data.

**Variant analysis by genome-wide sequencing**-- To call variants (SNPs/indels) in the transfected lines compared to a previously sequenced control 3D7 line, genomic DNA reads were first trimmed to adapters and aligned to the *Homo sapiens* genome (assembly GRCh38) to remove human-mapped reads. Remaining reads were aligned to the *P. falciparum* 3D7 genome using bwa (version 0.7.17) and PCR duplicates were removed using PicardTools (Broad Institute). GATK HaplotypeCaller (https://gatk.broadinstitute.org/hc/en-us) was used to call variants between the sample and the 3D7 reference genome for both the transfected lines and the 3D7 control. Only variants that were present in both transfected lines but not the 3D7 control line were kept. We examined only coding-region variants and removed those that were synonymous variants or were located in *var*, *rifin*, or *stevor* genes. Quality control of variants was done by hard filtering using GATK guidelines.

**In vitro safety profile--** The Enhanced Cross Screen Panel (X-Screen, eXP) consists of *in vitro* assays designed to measure the effect of test compounds on receptor sites, ion channels, transporters, and enzymes, and in phenotypic assays. These assays are generally run for small-molecule compounds (NCEs) to help define selectivity and/or to identify potential safety concerns. These assays can also aid mode of action identification of an in vivo effect produced by a test agent.

**Tolerability study--** Male CD1 (20-22g Envigo) were used in tolerability studies.
All animal studies were ethically reviewed and carried out in accordance with European Directive 2010/63/EEC and the GSK Policy on the Care, Welfare and Treatment of Animals. The animals were housed in cages (Tecniplast®). The cages contained autoclaved dust free corncob bedding (Rehofix MK2000, 1.7-2.2 mm particle size) and carry filters covers. Animal maintenance

standard g-irradiated standard rodent diet (Envigo Teklad 2914) and ultra-filtered water was provided ad libitum. Environmental conditions: Photoperiods, simulated dawn/dusk facility, were adjusted to 12h of light (300 lux) and 12h darkness daily. The temperature range was 22± 2ºC. Relative humidity was 55% ± 10 and the air room was changed 15-20 times per hour. Environmental enrichment included nesting material (nesting cups, LBS), a red-tinted igloo (Datesand), a hanging clear tunnel (Datesand, and two wood blocks (Datensand).

**Protocol design--** Male CD1 mouse (n=1) was dosed with an initial dose at 100 mg/kg (1-day, oral route, once a day, 10ml/kg). The animals were observed during the first four hours after administration and after 24h. The neurobehavioral assessment Functional Observational Battery (FOB) was performed. Adverse effects were found at 100 m/kg and new dose was selected used the modified symptom-based Up-&-Down method and adjusted a constant factor (1.6). Other male CD1 mouse was dosed at 62.5mg/kg and showed adverse effects. Two mice were treated at 50 mg/kg and at this dose didn´t show adverse effects. To confirm the data, 3 animals per dose (100mg/kg and 63.4 mg/kg) and vehicle group with 1% Methylcellulose + 0.5% Tween (n=2) were dosed. Systemic exposure levels at different times were taken (0.5h, 2h, 4h, 8, 24, 48h) of treated animals (n=3 per group) and a toxicological profile was performed after 7 days of treatment (hematology, clinical chemistry, and necropsy). Additionally in the same animals with the same scheduling sampling, blood samples of 15 uL were taken from lateral tail vein and recovered in a vial with the same quantity of water, mix and freeze until bioanalytical quantification for the determination of blood level exposures and pharmacokinetic analysis.

**Toxicological Profile--** The bodyweight of animals was recorded before dosage, 24 hours after and every day in animals kept for 7 days. Any sign of toxicity was recorded for 4h and 24h hours after drug administration (Functional Observational Battery). All animals were sacrificed by $CO_2$ inhalation. Blood samples for analysis were then taken by intracardiac puncture and fractionated to blood recount and serum analysis. The blood recount was carried out in the Abacus5 junior Vet → (Practice CVM S.L.L). Clinical chemistry was performed by mean analysis of whole blood in Vetscan→ (ABAXIS) & F560 (Menarini→) analyzer and by mean analysis of plasma, obtained after centrifugation at 1800g rpm for 10 min in S-Microvette tubes (Sarstedt→) with Heparin-Lithium. The plasma parameters tested were: glucose, ALP, AST, ALT, GGT, LDH, AMY, CK

Calcium, Phosphate, Sodium, Potassium, total bilirrubin, total protein, albumin, blood ureic nitrogen (BUN), creatinine, triglycerides, cholesterol. After necropsy, several organs (liver, kidney, spleen, Caecum, and stomach) were removed and weighed. Liver, kidney, and spleen were fixed in formalin (formaldehyde 10%) to further histopathology study (Hematoxilin/Eosin stain).

**In vivo Efficacy--** MED6-189 oral efficacy was evaluated using a standard 4-day test in female NOD-*scid IL-2R*$\gamma^{null}$ (NSG) mice engrafted with human erythrocytes and infected with $20\times10^6$ *Plasmodium falciparum*-infected erythrocytes (*P. falciparum Pf*3D7$^{0087/N9}$). The experiment was conducted by GlaxoSmithKline (Tres Cantos) and was ethically reviewed and carried out in accordance with European Directive 2010/63/EEC and the GSK Policy on the Care, Welfare and Treatment of Animals. The mice were obtained from Charles River Labs, bred specifically for research and certified pathogen free.

The human biological samples were sourced ethically, and their research use was in accord with the terms of the informed consent under an IRB/EC approved protocol. Erythrocyte concentrates from malaria-negative donors were provided by *Biobancos de Castilla y Leon and Centro de Transfusiones de Madrid, Spain.* Research was conducted according to POL-GSKF-410 and was in accord with the terms of the informed consent of each donor.

NSG mice were engrafted daily with human red blood cells intravenously injected in a final volume of 0.7 ml (50% hematocrit). Mice were engrafted during two weeks prior to *P. falciparum* infection, mice achieve around 60% of human red blood cells in mice peripheral blood.

*P. falciparum* infections were performed by intravenous inoculation. All mice were randomly assigned to their corresponding treatment. MED6-189 was given by oral gavage at doses of 50 mg/kg per day for four consecutive days. Treatment started at day 3 and finished at day 6 after infection. In all cases, parasitemia was assessed in samples from peripheral blood obtained at days 3, 4, 5, 6, and 7 after infection. Fresh samples of peripheral blood from *P. falciparum*-infected mice were stained with TER-119-Phycoerythrine (marker of murine erythrocytes) and SYTO-16 (nucleic acid dye) and then analyzed by flow cytometry (FACSCalibur, BD). Efficacy was determined using flow cytometry data as the percentage of Parasitemia reduction compared with Vehicle-treated group. MED6-189 levels in blood after oral administration were measured in serial blood samples obtained during the 23 h period after the first dose in all mice. Peripheral blood samples (15 µL) were taken at frequent intervals, mixed with 30 µL of $H_2O$ milli Q and immediately frozen on dry ice. The frozen samples were stored at −80 ˚C until analysis. Vehicle-treated mice experienced the same blood-sampling regimen (table S9).

**Bioanalysis and pharmacokinetics analysis: LC-MS--**

**Bioanalysis of in vivo samples: Sample pretreatment--** Before the analysis 15ul of blood samples were defrost at ambient temperature and mixture with 200ul of ACN:MeOH (80:20). After protein precipitation samples were filtered by 0.45um filter plate (Multiscreen Solvinert 0.45um FTPE, Millipore). The 2ul TFA (Trifluoroacetic acid, sigmaAldrich) were added to each filtered sample in order to derivatize the compound to formamide analogue. This process occurs spontaneously.

**LC-MS analysis--** An Acquity Ultra-Performance liquid chromatography (UPLC) system (Waters Corp., Milford, MA, USA) couple to a triple quadrupole mass spectrometer (API 4000™, AB Sciex, Foster City, CA, USA) was used for the analysis. The MS/MS transition was 311.340 / 203.100.

The chromatographic separation was carried out at 0.4ml/min in a Acquity UPLC™ BEH C18 column (50x2.1 mm i.d., 1.7 mm; Waters Corp.) at 40ºC with Acetonitrile (sigmaAldrich) and 0.1% formic acid as eluents.

**Pharmacokinetic analysis--** Blood concentration time data were analyzed by Non-Compartmental PK analysis using Phoenix WinNonlin software (Certara NY, US) to calculate PK parameters. The maximum observed concentration (Cmax) and the time to reach it (Tmax), were determined. The area under concentration-time curve from dosing at time zero to the last measured concentration time (AUClast) and the total area under concentration-time curve from time zero extrapolated to infinity (AUCinf), were calculated using a linear trapezoidal rule.

### Supplemental Figures

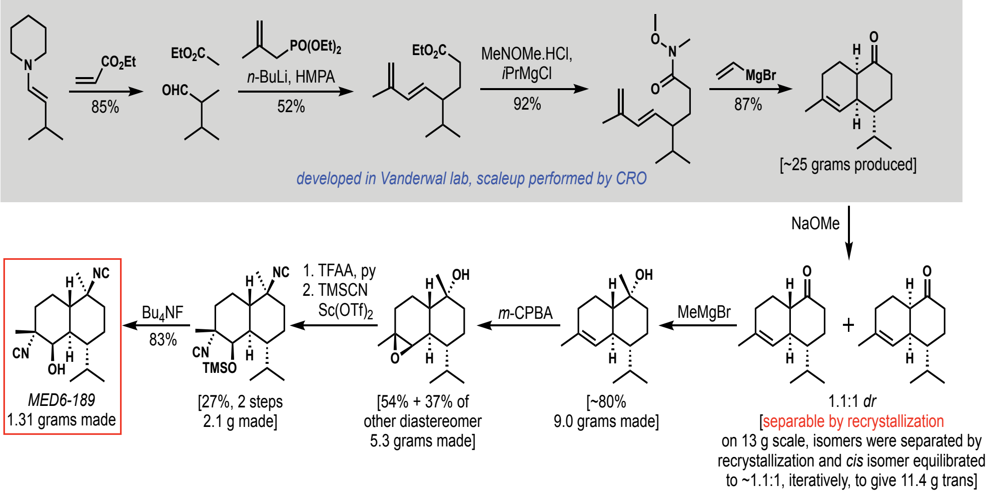


**Figure S1.** Route for the synthesis of MED6-189 on gram scale (See Supporting Information S1).

A

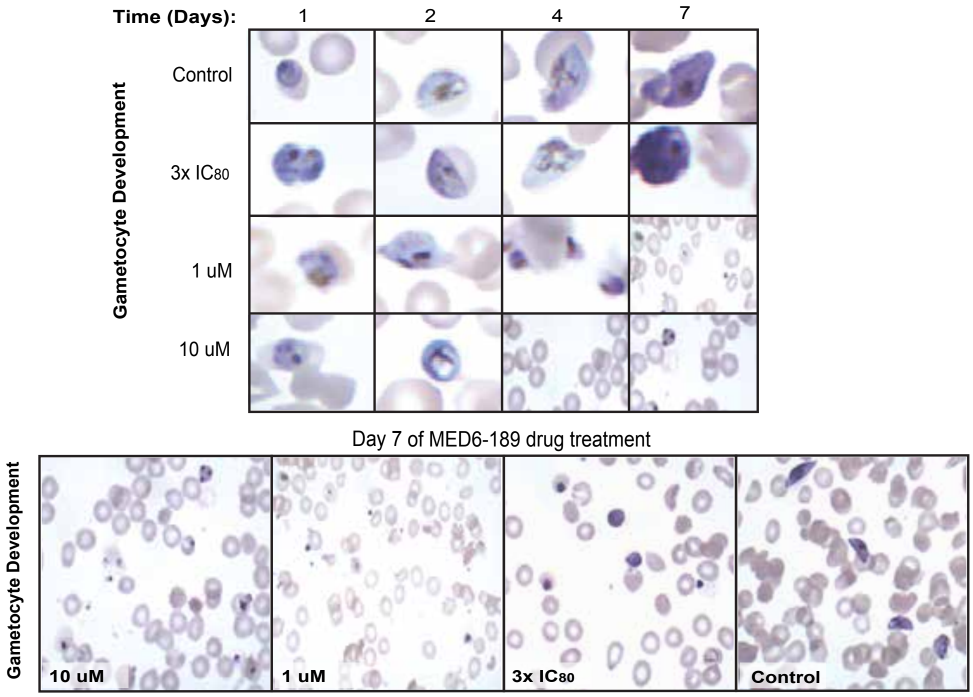


B

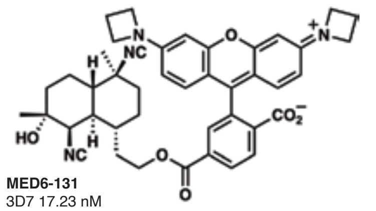


C

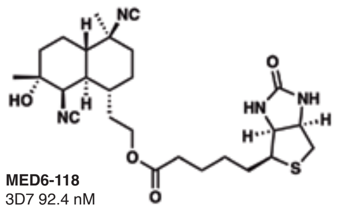


**Figure S2A.** Representative chemical profile of JF549-labelled kalihinol analogue. **B.** Representative chemical profile of the biotinylated kalihinol analogue. **C.** Sexual stage developmental impact of *P. falciparum* NF54 parasites incubated with MED6-189at 3xIC80.

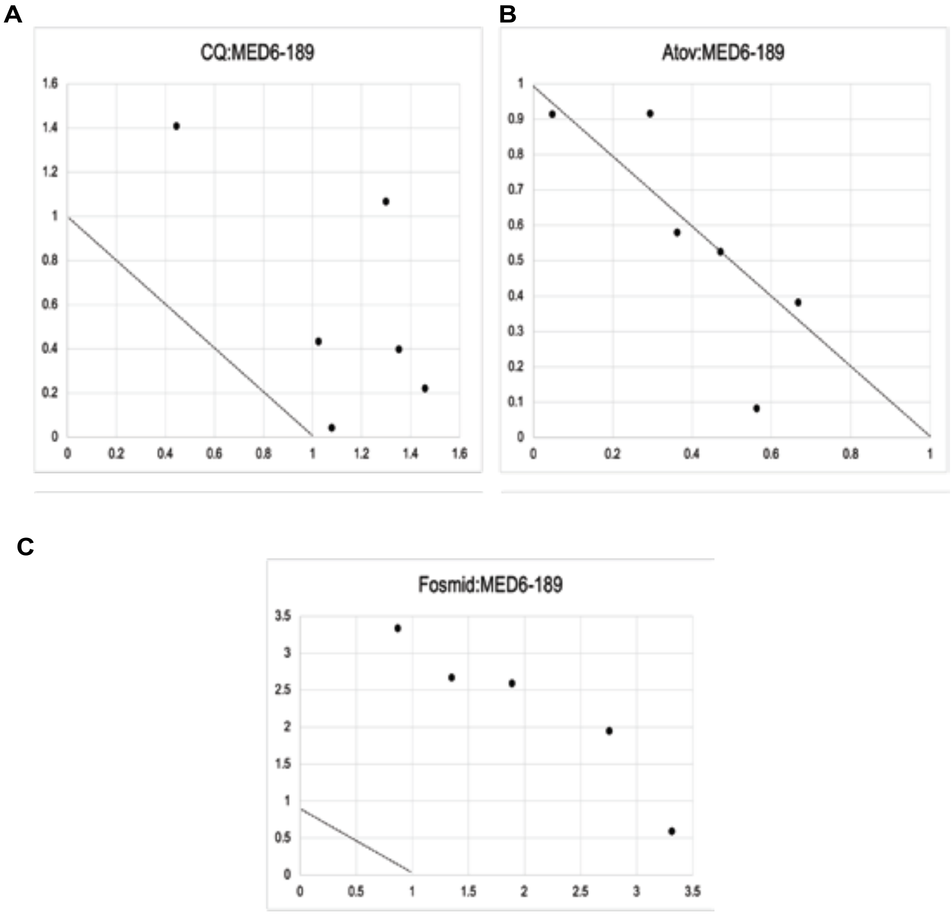


**Figure S3.** Normalized isobolograms demonstrating drug interaction (synergism, indifference, antagonism) between apicoplast inhibitors and MED6-189.

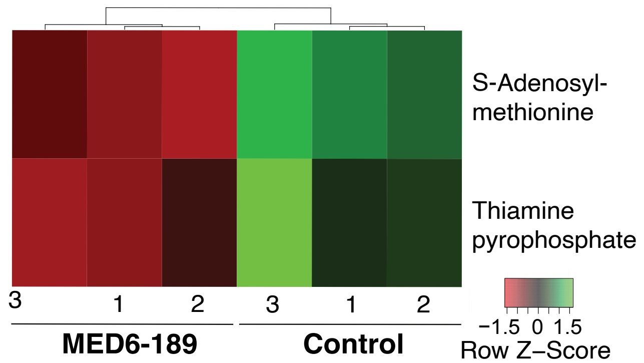


**Figure S4.** Heatmap visual representation of polar metabolites significantly upregulated (green) or downregulated (red) in response to MED6-189 treatment. $Log_2$ transformation was applied to the data to compute q-value (BH adjusted p-value) and p-value from Welch's t-test or ANOVA test.

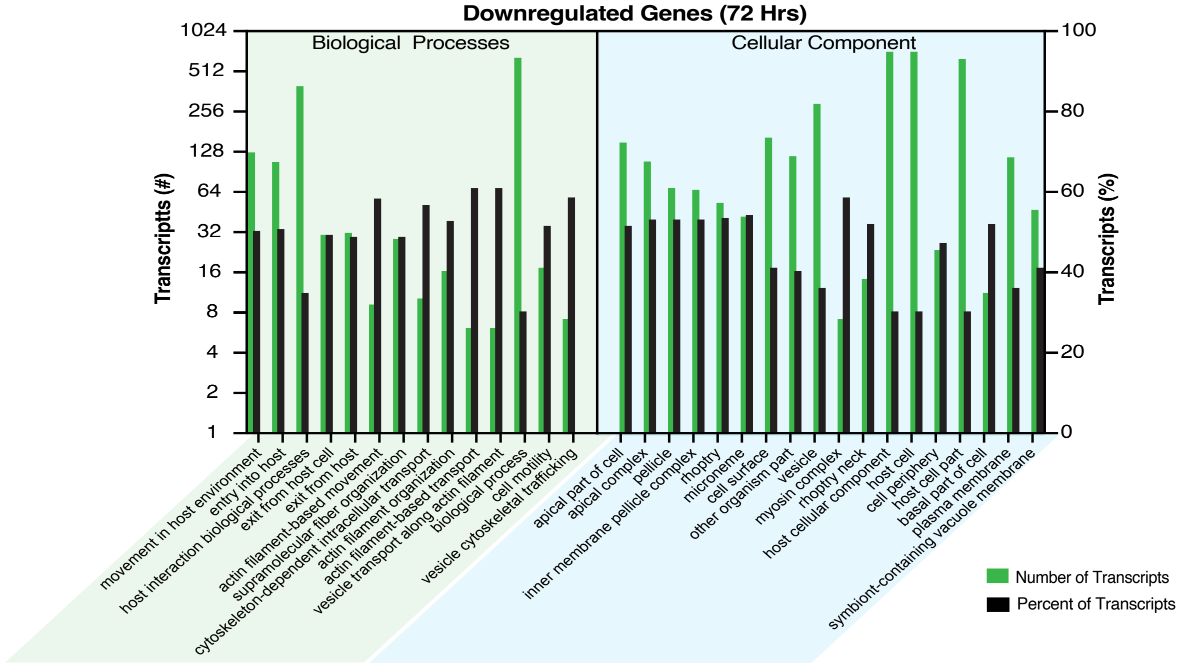
Downregulated Genes (72 Hrs)
Biological Processes
Cellular Component
Transcriptts (#)
Transcripts (%)
1024
512
256
128
64
32
16
8
4
2
1
100
80
60
40
20
0
movement in host environment
entry into host
host interaction biological processes
exit from host cell
exit from host
actin filament-based movement
supramolecular fiber organization
cytoskeleton-dependent intracellular transport
actin filament organization
actin filament-based transport
vesicle transport along actin filament
biological process
cell motility
vesicle cytoskeletal trafficking
apical part of cell
apical complex
inner membrane pellicle complex
pellicle
rhoptry
microneme
cell surface
other organism part
vesicle
myosin complex
rhoptry neck
host cellular component
host cell
cell periphery
host cell part
basal part of cell
plasma membrane
symbiont-containing vacuole membrane
Number of Transcripts
Percent of Transcripts

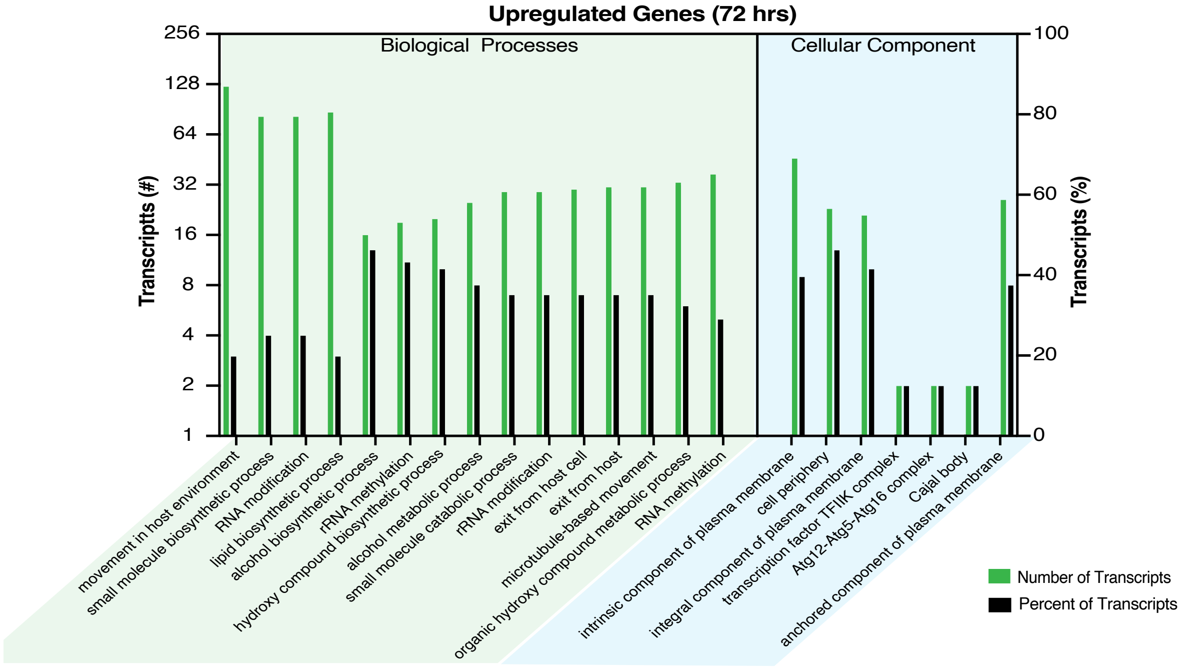
Upregulated Genes (72 hrs)
Biological Processes
Cellular Component
Transcriptts (#)
Transcripts (%)
256
128
64
32
16
8
4
2
1
100
80
60
40
20
0
movement in host environment
small molecule biosynthetic process
RNA modification
lipid biosynthetic process
alcohol biosynthetic process
rRNA methylation
hydroxy compound biosynthetic process
alcohol metabolic process
small molecule catabolic process
rRNA modification
exit from host cell
exit from host
microtubule-based movement
organic hydroxy compound metabolic process
RNA methylation
intrinsic component of plasma membrane
cell periphery
integral component of plasma membrane
transcription factor TFIIK complex
Atg12-Atg5-Atg16 complex
Cajal body
anchored component of plasma membrane
Number of Transcripts
Percent of Transcripts

**Figure S5.** Gene ontology bar graphs representing parasite transcriptomic responses to MED6-189. Gene ontology bar graph representing the top downregulated (top) and upregulated (bottom) transcripts from parasite samples cultured with either DMSO or MED6-189 collected at 72 HPS. Represented transcripts are filtered by highest number and percentage among most significantly expressed GO terms (Adjusted p-value $\leq 0.05$).

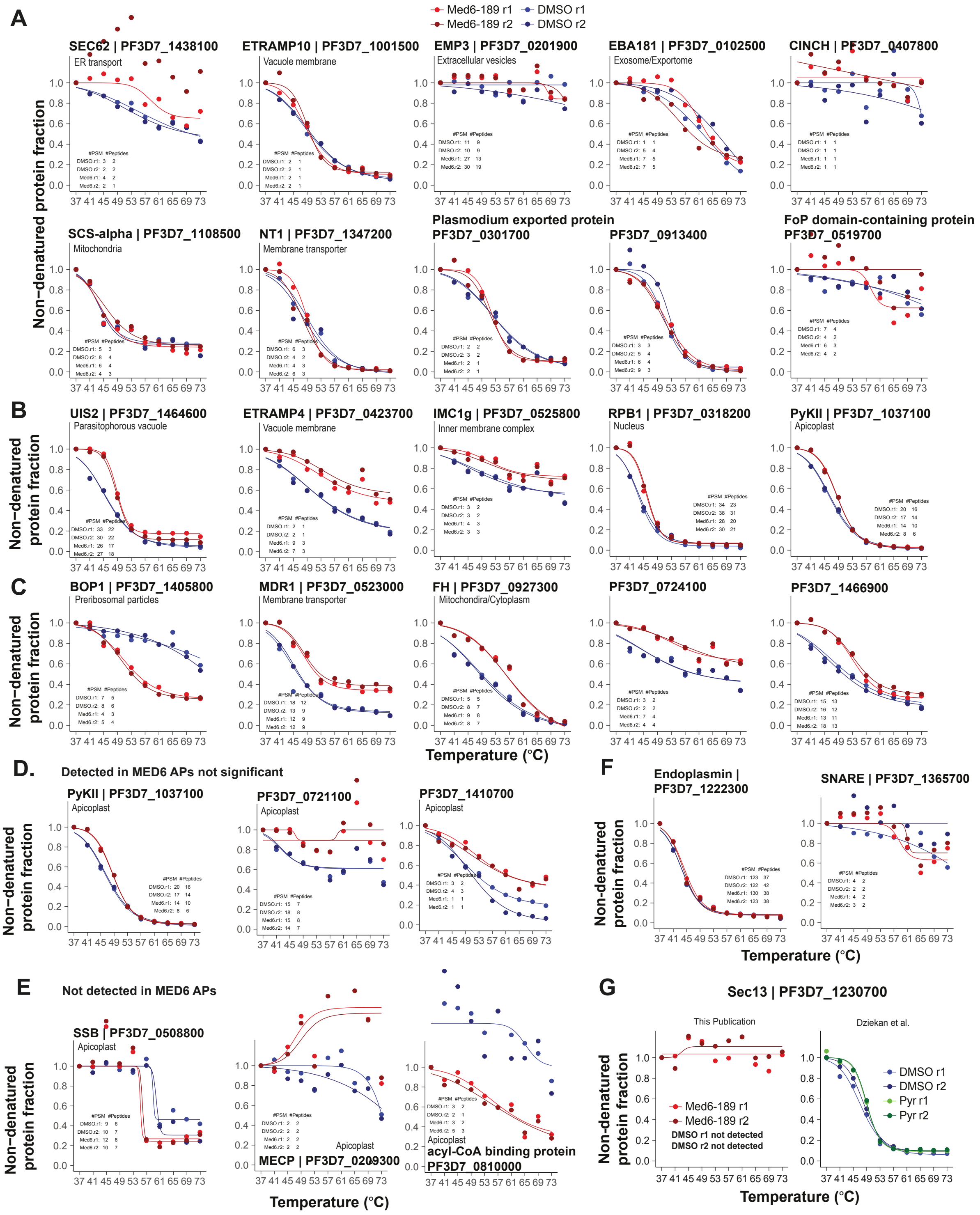
A
Med6-189 r1
Med6-189 r2
DMSO r1
DMSO r2
SEC62 | PF3D7_1438100
ER transport
ETRAMP10 | PF3D7_1001500
Vacuole membrane
EMP3 | PF3D7_0201900
Extracellular vesicles
EBA181 | PF3D7_0102500
Exosome/Exportome
CINCH | PF3D7_0407800
Non−denatured protein fraction
SCS-alpha | PF3D7_1108500
Mitochondria
NT1 | PF3D7_1347200
Membrane transporter
Plasmodium exported protein
PF3D7_0301700
PF3D7_0913400
FoP domain-containing protein
PF3D7_0519700
B
UIS2 | PF3D7_1464600
Parasitophorous vacuole
ETRAMP4 | PF3D7_0423700
Vacuole membrane
IMC1g | PF3D7_0525800
Inner membrane complex
RPB1 | PF3D7_0318200
Nucleus
PyKII | PF3D7_1037100
Apicoplast
C
BOP1 | PF3D7_1405800
Preribosomal particles
MDR1 | PF3D7_0523000
Membrane transporter
FH | PF3D7_0927300
Mitochondria/Cytoplasm
PF3D7_0724100
PF3D7_1466900
Temperature (°C)
D.
Detected in MED6 APs not significant
PyKII | PF3D7_1037100
Apicoplast
PF3D7_0721100
Apicoplast
PF3D7_1410700
Apicoplast
F
Endoplasmin |
PF3D7_1222300
SNARE | PF3D7_1365700
Temperature (°C)
E
Not detected in MED6 APs
SSB | PF3D7_0508800
Apicoplast
Apicoplast
MECP | PF3D7_0209300
Apicoplast
acyl-CoA binding protein
PF3D7_0810000
Temperature (°C)
G
Sec13 | PF3D7_1230700
This Publication
Dziekan et al.
Med6-189 r1
Med6-189 r2
DMSO r1 not detected
DMSO r2 not detected
DMSO r1
DMSO r2
Pyr r1
Pyr r2
Temperature (°C)

**Figure S6.** CETSA melt-curve analysis of *P. falciparum* lysates treated with MED6-189. Protein stabilization curves for *P. falciparum* proteins identified in lysates treated with or without 100 μM Med6-189 (MED6). Stabilization is assessed from the relative amount of soluble protein remaining (y axis) after thermal challenge (x axis). Sample replicates are plotted in shades of red (MED6-189 treated) or blue (DMSO controls). **A.** Proteins enriched but not showing a distinct temperature-mediated stabilization profile in the presence of MED6-189. **B.** Proteins detected in affinity purification dataset but did not pass significance cut-off. **C.** Proteins showing significant stabilization yet not detected in the MED6-189 protein affinity purification datasets. **D.** Proteins showing significant stabilization in thermal profiling datasets as well as detected in the MED6-189 affinity purification datasets but did not reach threshold cutoffs. **E.** Proteins showing significant stabilization yet not detected in the MED6-189 protein affinity purification datasets. **F.** Thermal profiling curves for Endoplasmin and SNARE protein candidates that were shown to be significantly detected in MED6-189 affinity profiles but not shown to have significant stabilization in CETSA assays. **G.** Thermal profiling curves for SEC13 in samples incubated with MED6-189, Pyrimethamine and DMSO controls.

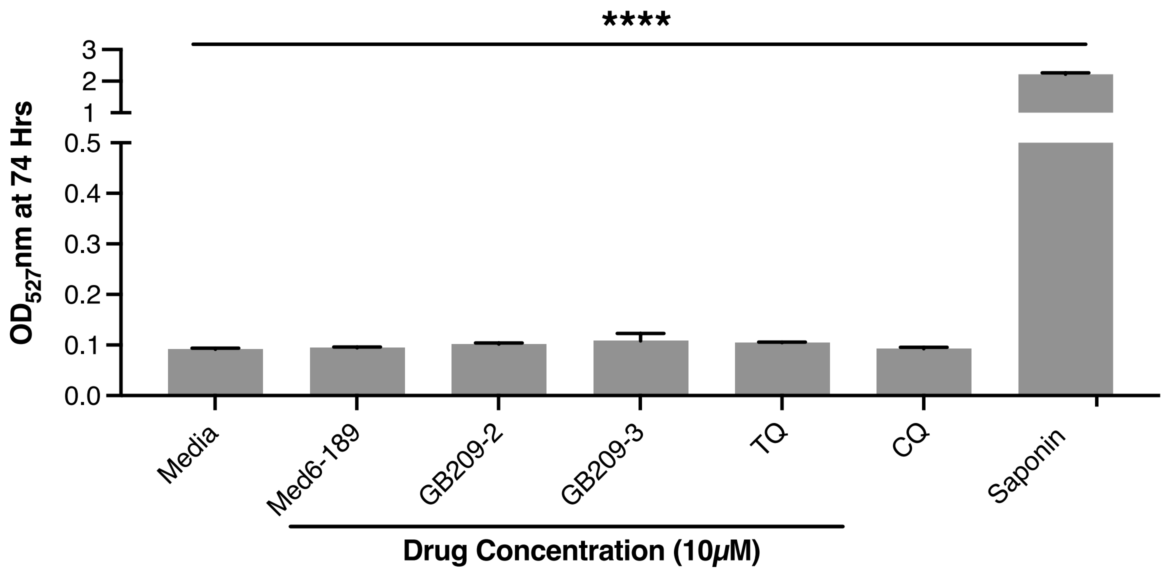


**Figure S7.** Enhanced Cross Screen Panel (eXP) graph. Pharmacological and toxicology values among compounds of interest

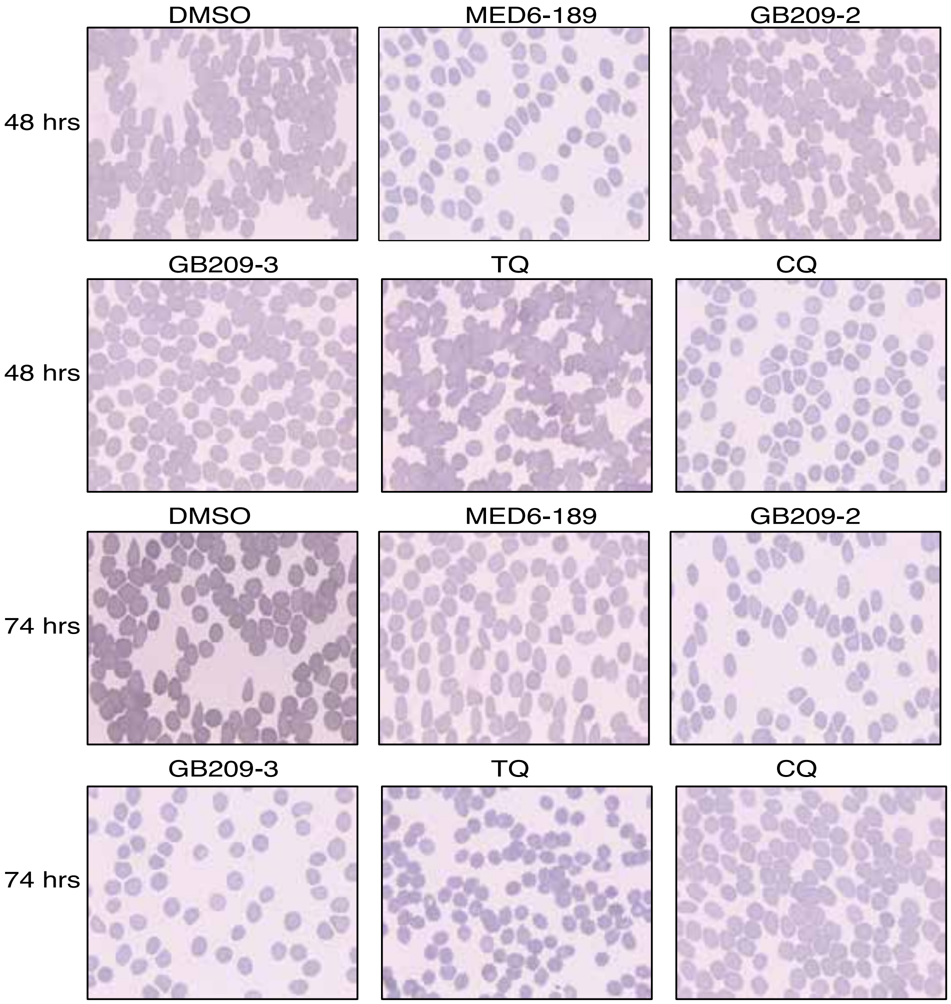


**Figure S8.** Giemsa smeared culture images depicting the hemolytic activity assays for MED6-189 and two formamide derivatives. No hepatoxicity was observed in response to MED6-189, GB209-2 and GB209-3 compounds tested after 48Hr and 72Hr drug incubation.

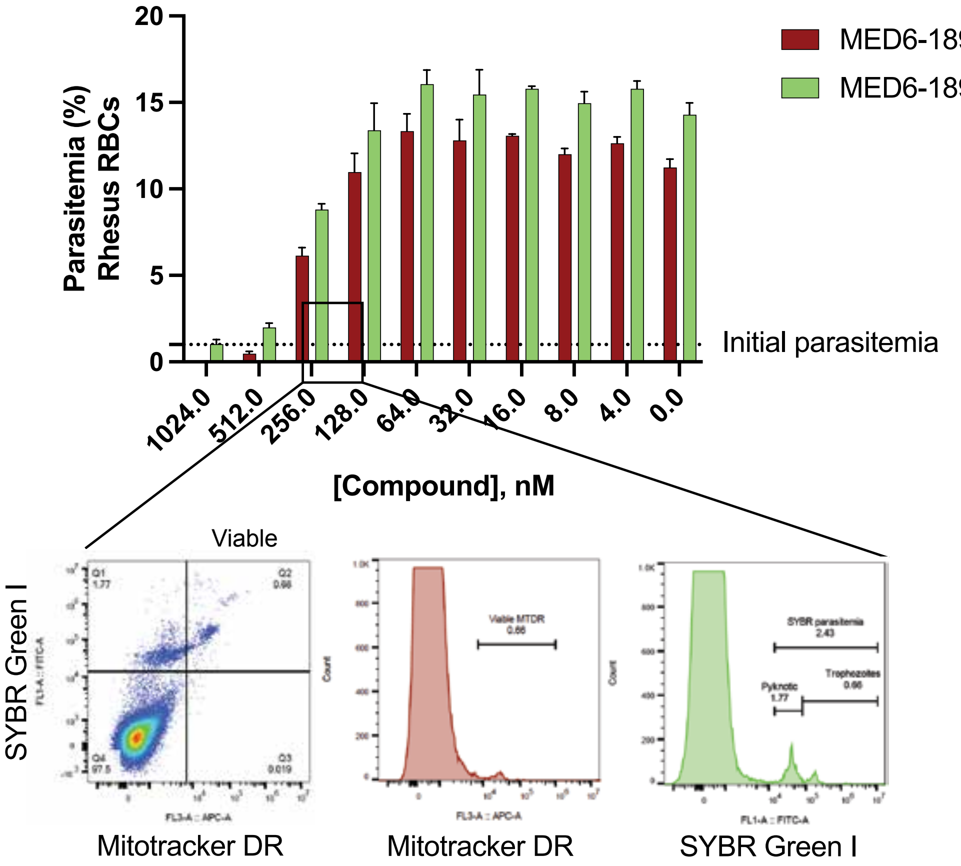


**Figure S9.**Flow cytometry panels illustrating results of SYBR green staining as well as Mitotracker Deep Red to determine viable parasites. Data for the IC50 plots are result of double positive staining (SYBR+, MTDR+) - representing viable parasites.

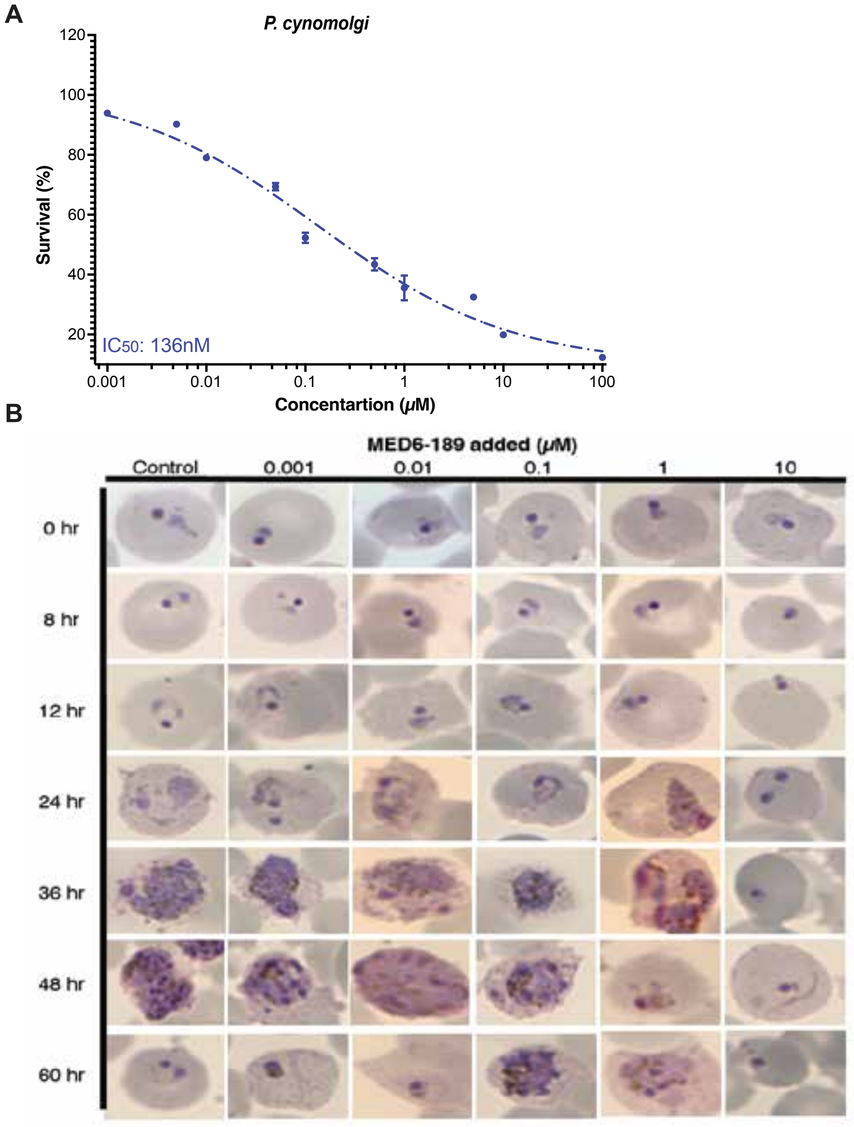


**Figure S10A.** Dose-dependent response of MED6-189 on *P. cynomolgi* M/B strain parasites. The figures exhibit the logarithmic growth of parasites (Y-axis) corresponding to varying drug concentrations (X-axis). The error bars represent the standard deviation obtained from two independent experiments conducted in triplicate. The regression line represents a nonlinear regression analysis (Variable slope with four parameters, least squares fit). **B.** Giemsa-stained images of *P. cynomolgi* M/B strain parasites incubated with MED6-189 examined by light microscopy at different developmental stages of the parasite cell cycle.
