## Supporting Information S1 for "A Potent Kalihinol Analogue Disrupts Apicoplast Function and Vesicular Trafficking in *P. falciparum* Malaria": Supporting Information_S1_ICTs_11_6_23.docx

**General experimental procedures**:

**S1.** Supplemental Information for the synthesis of MED6-189 on gram scale ( See Fig. S1).

Tetrahydrofuran (THF, Fisher, HPLC Grade) was dried by percolation through a column packed with neutral alumina and a column packed with Q5 reactant, a supported copper catalyst for scavenging oxygen, under a positive pressure of argon. Solvents for workup and chromatography include hexanes (Fisher, ACS Grade), ethyl acetate (EtOAc, Fisher, ACS Grade), dichloromethane (DCM, Fisher, ACS Grade), chloroform (CHCl_3_, Fisher, ACS Grade), methanol (MeOH, Fisher, ACS grade) and were used as obtained commercially unless otherwise stated. Column chromatography was performed using EMD Millipore 60 Å (0.040–0.063 mm) silica gel (SiO_2_). Analytical thin-layer chromatography was performed on Merck silica gel 60 F254 TLC plates. Visualization was accomplished with UV (254 or 210 nm), p-anisaldehyde, potassium permanganate, iodine/silica, and/or heat as developing agents. Chloroform-d (CDCl_3_, D 99.8%, DLM-7) was purchased from Cambridge Isotope Laboratories. Methanol-d_4_ (D_3_COD, 99.8%) was purchased from Sigma–Aldrich. Proton and carbon magnetic resonance spectra (^1^H NMR and ^13^C NMR) were recorded at 298K on a Bruker CRYO500 (500 MHz, ^1^H; 125 MHz, ^13^C) or a Bruker AVANCE600 (600 MHz, ^1^H; 151 MHz, ^13^C) spectrometer with solvent or residual solvent resonance as the internal standard (^1^H NMR: [CDCl_3_], CHCl_3_ at 7.26 ppm, [D_3_COD], HD_2_COD at 3.31 ppm, ^13^C NMR: CDCl_3_ at 77.16 ppm). ^1^H NMR data are reported as follows: chemical shift, multiplicity (s = singlet, d = doublet, t = triplet, q = quartet, dd = doublet of doublets, ddd = doublet of doublet of doublets, td = triplet of doublets, tdd = triplet of doublet of doublets, qd = quartet of doublets, m = multiplet, br s = broad singlet), coupling constants (Hz), and integration. Mass spectrometry data were obtained from the University of California, Irvine Mass Spectrometry Facility. High resolution mass spectra (HRMS) were recorded on a Waters LCT Premier spectrometer using ES+ and data are reported in the form of (m/z).

**Synthetic preparation of MED6-189**: Following modified procedures of Taber  *et al* (1979), Shenvi *et al* (2013), and Vanderwal *et al* (2017), 1.31 g of racemic MED6-189 was prepared according to the route shown in figure S1 below.

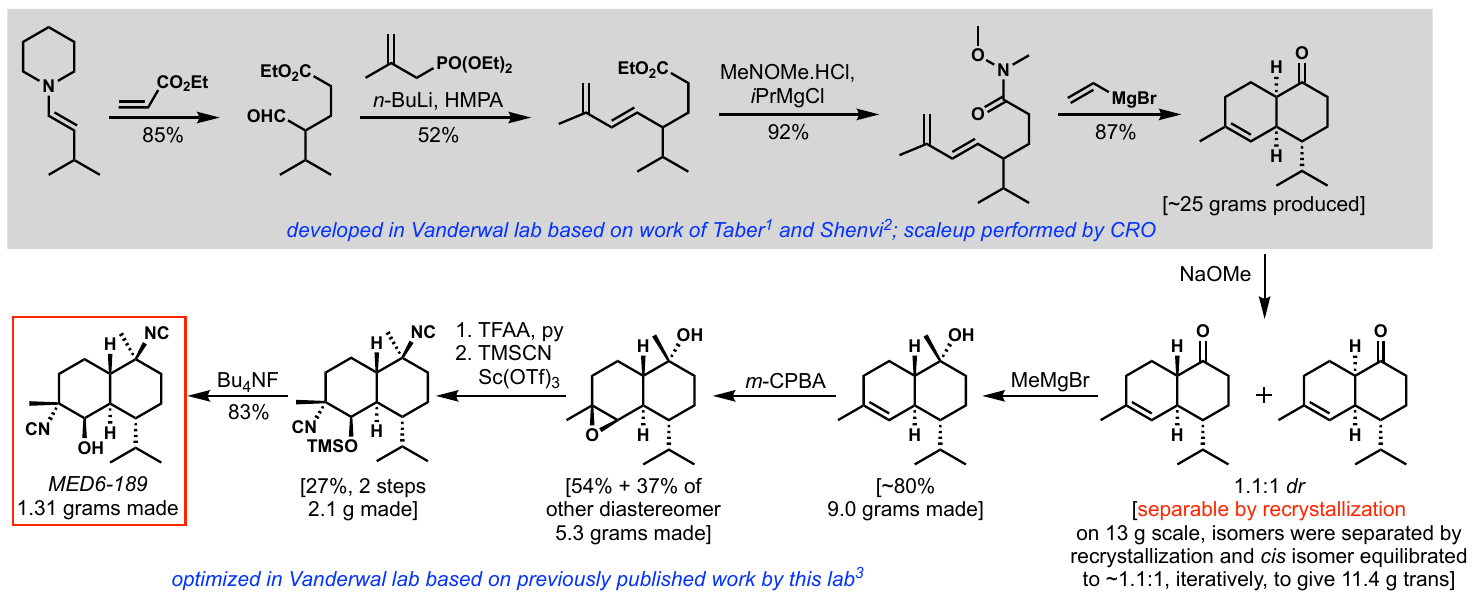

**Fig. S1.** Route for the synthesis of MED6-189 on gram scale.

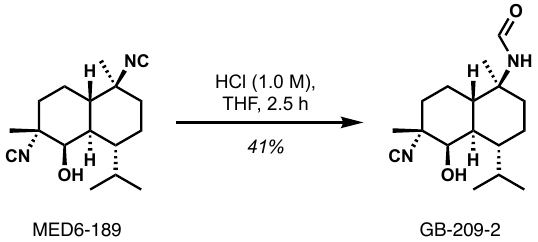

**Synthetic preparation of GB-209-2**: To a 1-dram vial containing MED6-189 (4.8 mg, 17 µmol) and a magnetic stir bar was added THF (tetrahydrofuran, 1.7 mL) and hydrochloric acid (0.87 mL, 1.0 M, 870 µmol). The mixture was stirred at room temperature for 2.5 h. The reaction was quenched by the addition of sodium bicarbonate (saturated aqueous solution, 1.0 mL). The organic phase was extracted using CH­_2_Cl_2_ (3 x 1 mL) and the combined extracts were dried using anhydrous magnesium sulfate. The volatiles were removed under reduced pressure in a rotary evaporator then under high vacuum to afford a colorless residue that was further purified using flash column chromatography (eluent: EtOAc:hexanes = 40:60, both distilled). GB-209-2 was obtained as a colorless oil that solidified upon standing (2.1 mg, 41% yield). The material will precipitate from DCM upon addition of hexanes to obtain the product in powdered form.

**Note:** GB-209-2 was observed as two different amide rotameric species (R^1^, R^2^ ratio = 55:45, respectively) under ambient conditions, complicating NMR spectroscopic analysis.

**^1^H NMR** (500 MHz, CDCl_3_, residual CHCl_3_ at 7.26 ppm): δ 8.28 (R^1^: [O=C**H**], d, *J* = 12.3 Hz, 1H), 8.05 (R^2^: [O=C**H**], d, *J* = 2.1 Hz, 1H), 5.52 (R^1^: [N**H**], d, *J* = 12.2 Hz, 1H), 5.09 (R^2^: [N**H**], s, 1H), 3.81 (R^1^: [HO–C**H**], apparent s, 1H), 3.79 (R^2^: [HO–C**H**], apparent s, 1H), 2.32 (R^1^: [HO–CH–C**H**], td, *J* = 12.1, 3.3 Hz, 1H), 2.26 (R^2^: [HO–CH–C**H**], td, *J* = 12.1, 3.3 Hz, 1H), 2.0–1.92 (m, 1H), 1.91–1.79 (m, 2H), 1.78–1.60 (m, 5H), 1.54–1.40 (m, 6H), 1.32–1.18 (m, 4H), 0.97–0.92 (m, 3H), 0.95 (R^1^: [CH(C**H_3_**)_2_], d, *J* = 7.1 Hz, 3H), 0.94 (R^2^: [CH(C**H_3_**)_2_], *J* = 7.1 Hz, 3H), 0.84 (R^1^: [CH(C**H_3_**)_2_], d, *J* = 7.0 Hz, 3H), 0.82 (R^2^: [CH(C**H_3_**)_2_], d, *J* = 7.0 Hz, 3H). **^13^C NMR** (126 MHz, CDCl_3_ at 77.16 ppm): δ 162.8, 160.5, 156.9, 156.8, 156.8, 156.4, 156.4, 156.3, 71.4, 71.2, 61.3, 61.2, 61.2, 61.1, 61.1, 61.0, 57.7, 55.8, 42.8, 41.8, 41.8, 41.8, 38.8, 38.7, 37.9, 36.9, 31.4, 31.3, 29.8, 26.7, 26.6, 25.9, 25.8, 21.5, 21.5, 21.3, 20.3, 20.1, 20.0, 19.4, 15.4. **HRMS (ES+)**: 315.2068 (*m/z* calc’d for C_17_H_28_N_2_O_2_ [M+Na]^+^:315.2048).

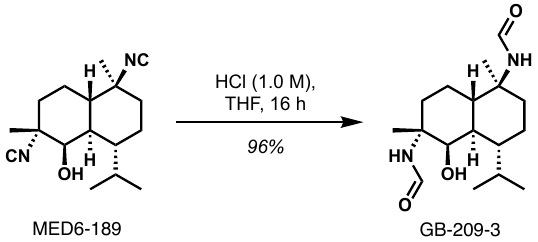

**Synthetic preparation of GB-209-3**: To a 1-dram vial containing MED6-189 (4.8 mg, 17 µmol) and a magnetic stir bar was added THF (tetrahydrofuran, 1.7 mL) and hydrochloric acid (0.87 mL, 1.0 M, 870 µmol). The mixture was stirred at room temperature for 16 h. The volatiles were removed under reduced pressure in a rotary evaporator then under high vacuum to afford bis-formamide GB-209-3 as a colorless oil that solidified upon standing (5.2 mg, 96% yield).

*Note*: GB-209-3 was observed as four different amide rotameric species (estimated ratio = 54:35:7:4) under ambient conditions, complicating NMR spectroscopic analysis:

**^1^H NMR** (500 MHz, D_3_COD, residual HD_2_COD at 3.31 ppm): δ 8.29–7.87 (2H, m), 4.54–3.70 (1H, m), 2.13–1.94 (2H, m), 1.85–1.76 (1H, m), 1.76–1.66 (1H, m), 1.65–1.46 (6H, m), 1.41 (3H, s), 1.39–1.32 (1H, m), 1.32–1.25 (2H, m), 1.25–1.11 (2H, m), 0.97–0.89 (3H, m), 0.79–0.70 (3H, m). **^13^C NMR** (126 MHz, D_3_COD at 49.00 ppm): 166.4, 163.7, 163.6, 163.6, 68.2, 68.1, 59.3, 59.1, 58.4, 58.4, 58.3, 58.2, 43.0, 43.0, 42.9, 41.7, 41.6, 40.9, 39.6, 39.4, 38.5, 38.4, 32.3, 32.1, 28.8, 26.4, 26.4, 26.3, 24.3, 24.3, 24.2, 22.2, 22.1, 22.0, 22.0, 21.9, 21.6, 20.8, 20.8, 20.4, 19.2, 19.1, 18.2, 18.1, 18.1, 15.5. **HRMS (ES+)**: 333.2143 (*m/z* calc’d for C_17_H_30_N_2_O_3_ [M+Na]^+^: 333.2154).

**Time course stability of MED6-189 to conditions of varying pH**:

Isonitriles are known to be somewhat reactive functional groups. While previous studies demonstrated that a closely related bis(isonitrile) compound had reasonable liver microsomal stability,^3^ we also wanted to account for the potential stomach-acid-catalyzed hydrolysis of the isonitriles in the course of in vivo experiments. We developed a mass spectrometric assay to understand the acid stability of the isonitriles with respect to hydrolysis to the corresponding formamides. For this assay, we generated authentic samples of the hydrolysis products GB-209-2 (faster reacting isonitrile converted to mono-formamide) and GB-209-3 (both isonitriles hydrolyzed to form bis-formamide). Each of these compounds were tested against Plasmodium falciparum strain 3D7 (triplicate) and found to have very little antiplasmodial activity (IC_50_ of 3.8 and 3.9 µM, respectively). These data, coupled with the reasonable stability of MED6-189 toward acid hydrolysis down to pH 4, strongly supports that a significant quantity of MED6-189 survives passage through the gut, and it is the parent compound that is responsible for the antiplasmodial activity.

Pure samples of MED6-189, GB-209-2, and GB-209-3 were calibrated using a Micromass® Quattro Premier XE mass spectrometer with triple quadrupole (ES+) ionization parameters described in the table below. MED6-189 was calibrated using both a parent mass transition (275 – 221 Da) and a fragment mass transition (248 – 221 Da) due to its low parent ion count. With these parameters, calibration curves were generated by analyzing pure samples of each compound at concentrations (nm) shown in the Calibration curves below in acetonitrile:water = 20:80.

| **a.** | Compound | M/Z Transition (g*mol^–1^) | Dwell (s) | Cone Voltage (V) | Collision Energy (eV) | Delay (s) | t_R_ (min) |
| --- | --- | --- | --- | --- | --- | --- | --- |
|  | MED6-189 | 275 – 221 | 0.100 | 30.0 | 10.0 | 0.010 | 2.15 |
|  | MED6-189 | 248 – 221 | 0.100 | 30.0 | 10.0 | 0.020 | 2.15 |
|  | GB-209-2 | 293 – 203 | 0.100 | 20.0 | 10.0 | 0.010 | 1.70 |
|  | GB-209-3 | 311 – 203 | 0.100 | 20.0 | 10.0 | 0.010 | 1.25 |

| **b*.*** | Time (min) | Flow Rate (mL*min^-1^) | %A | %B | Curve |
| --- | --- | --- | --- | --- | --- |
|  | Initial | 0.3 | 90 | 10 |  |
|  | 2.00 | 0.3 | 10 | 90 | 6 |
|  | 3.00 | 0.3 | 90 | 10 | 11 |
|  | 4.00 | 0.3 | 90 | 10 | 11 |

**Supporting Table 1. a.** Ionization (ES+) and ***b.*** inlet parameters for quantification of MED6-189 and formamides GB-209-2, and GB-209-3. Column Type: ACQUITY CSHTM Fluoro-Phenyl 1.7μm. Solvents A = formic acid (0.1% aq.) and B = acetonitrile.

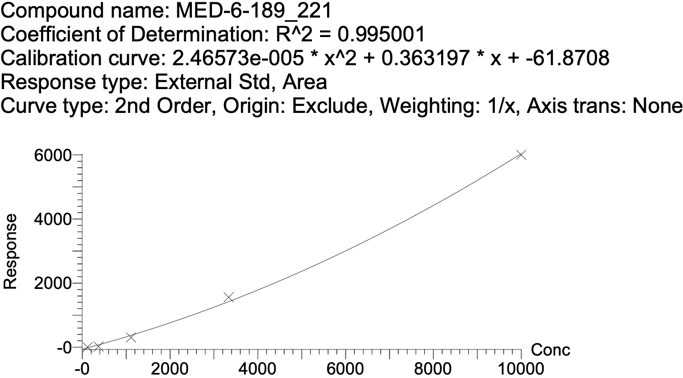

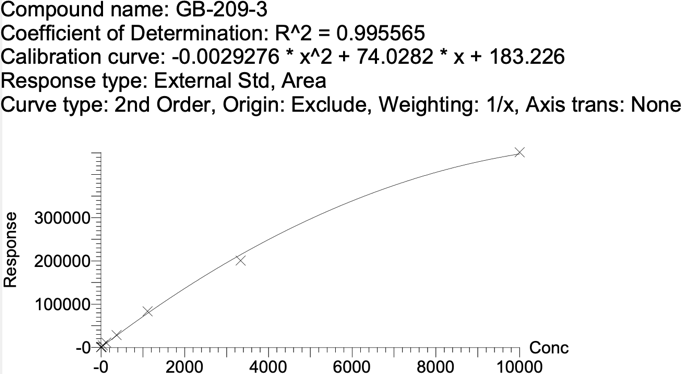

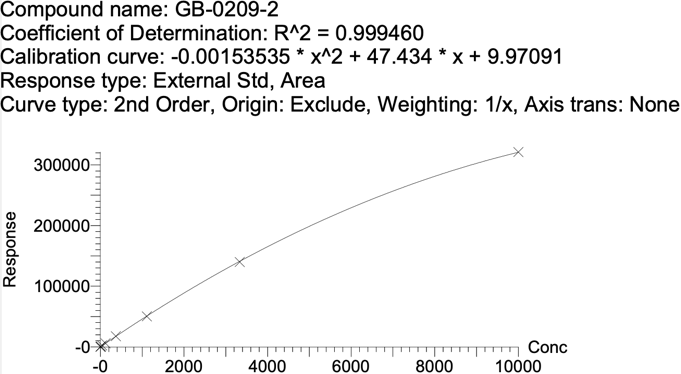

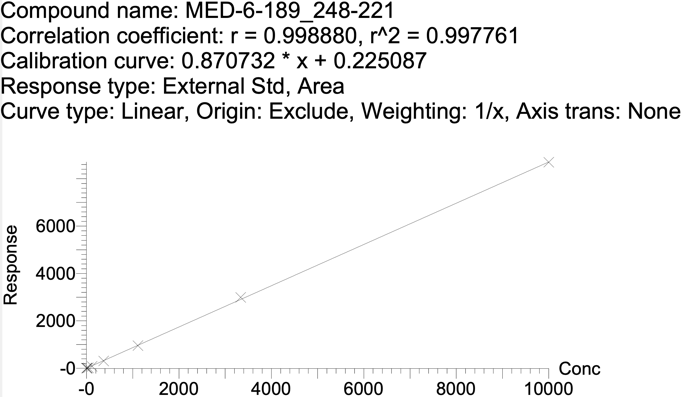

**Calibration curves for MED6-189, GB-209-2, and GB-209-3.**

A stock solution of MED6-189 (5.0 mM, 1.5 mg in 1.1 mL acetonitrile) was prepared. Aqueous buffer solutions of varying pH (400 μL, 0.1 M, KCl/HCl [pH = 2], sodium citrate/citric acid [pH = 3, 4, 5, 6], ammonium bicarbonate [pH = 7]) were charged with MED6-189 solution (100 μL) at room temperature so that the substrate concentration was 1.0 mM. At various time points (10 min, 30 min, 1 h, 2 h, 4 h, 8 h, 16 h) aliquots (10.0 μL) from each reaction mixture were diluted 300–fold using two serial 17.3-fold dilutions in neutral buffer solution (acetonitrile:ammonium bicarbonate solution [0.1 M, aqueous] = 1:4) so that the total concentration of substrate and products was approximately 3.3 μM. Diluted aliquots were immediately frozen at –20 ºC until analysis the following day. Using the calibration curves above, the concentrations of MED6-189, GB-209-2 and GB-209-3 were determined for each aliquot and plotted over time for respective pH values in the figure below describing the time course stability of MED6-189 to conditions of varying pH over time via quantitative analysis of formamide products GB-209-2 and GB-209-3.

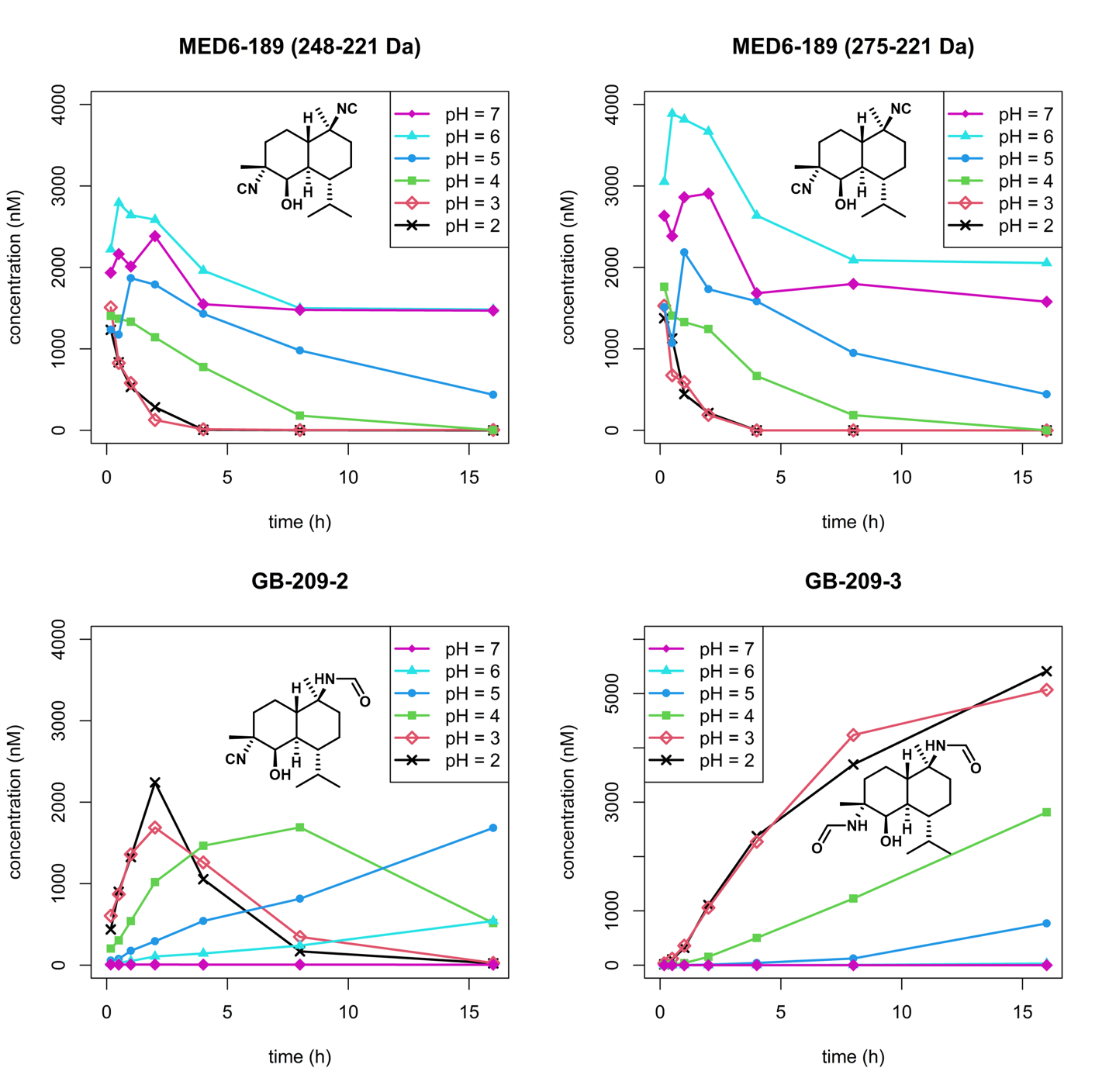

Time course stability of MED6-189 to conditions of varying pH over time via quantitative analysis of formamide products GB-209-2 and GB-209-3.

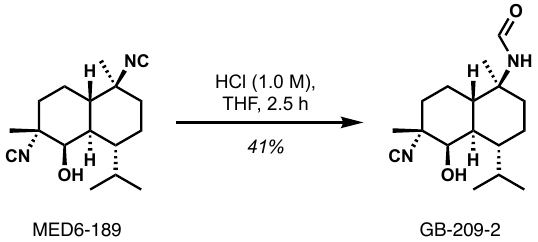

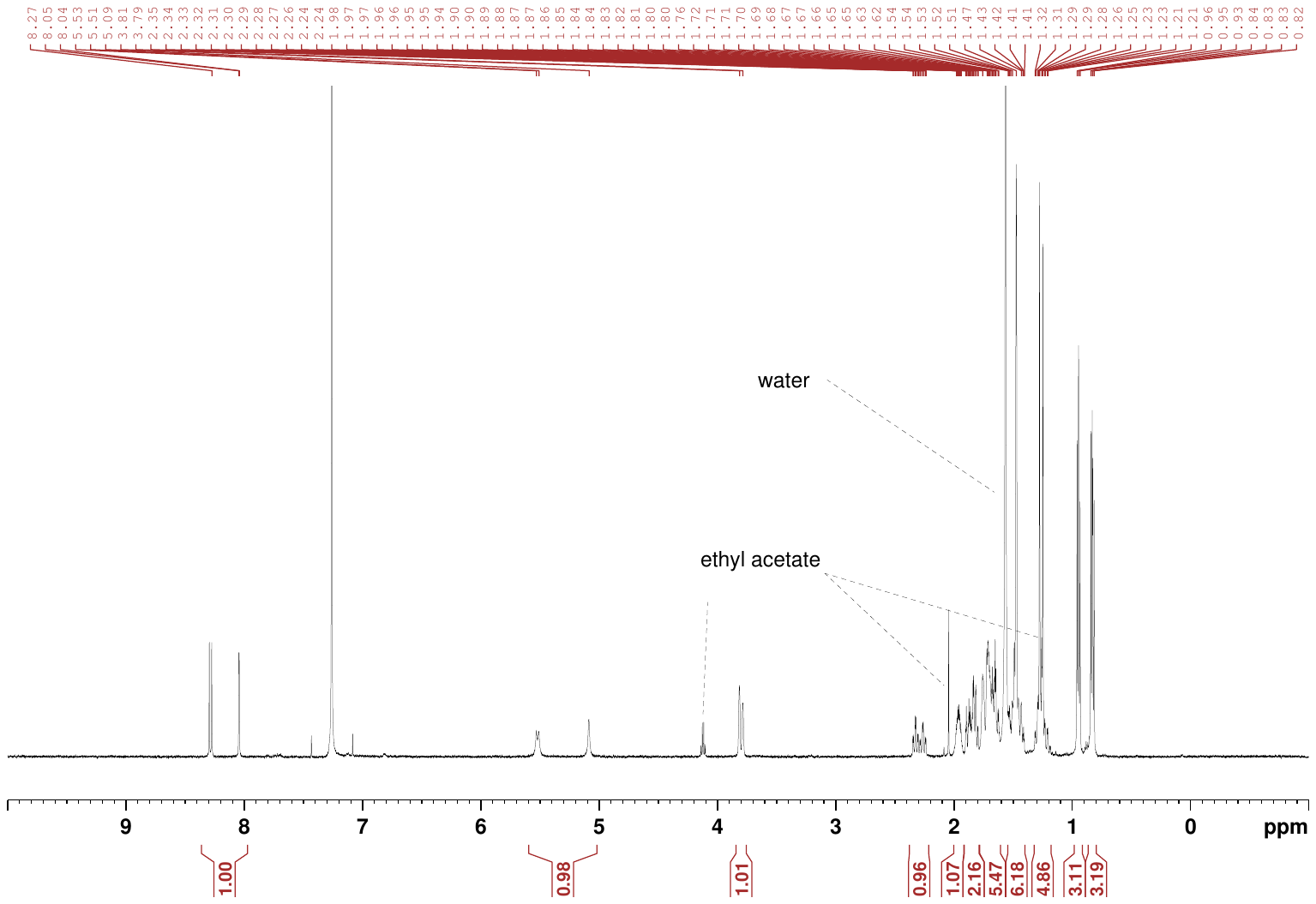

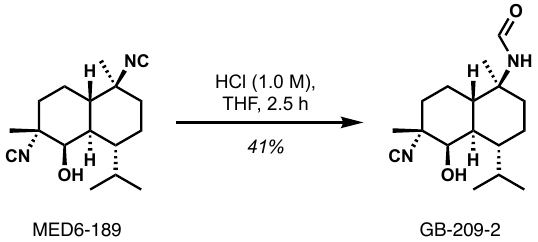

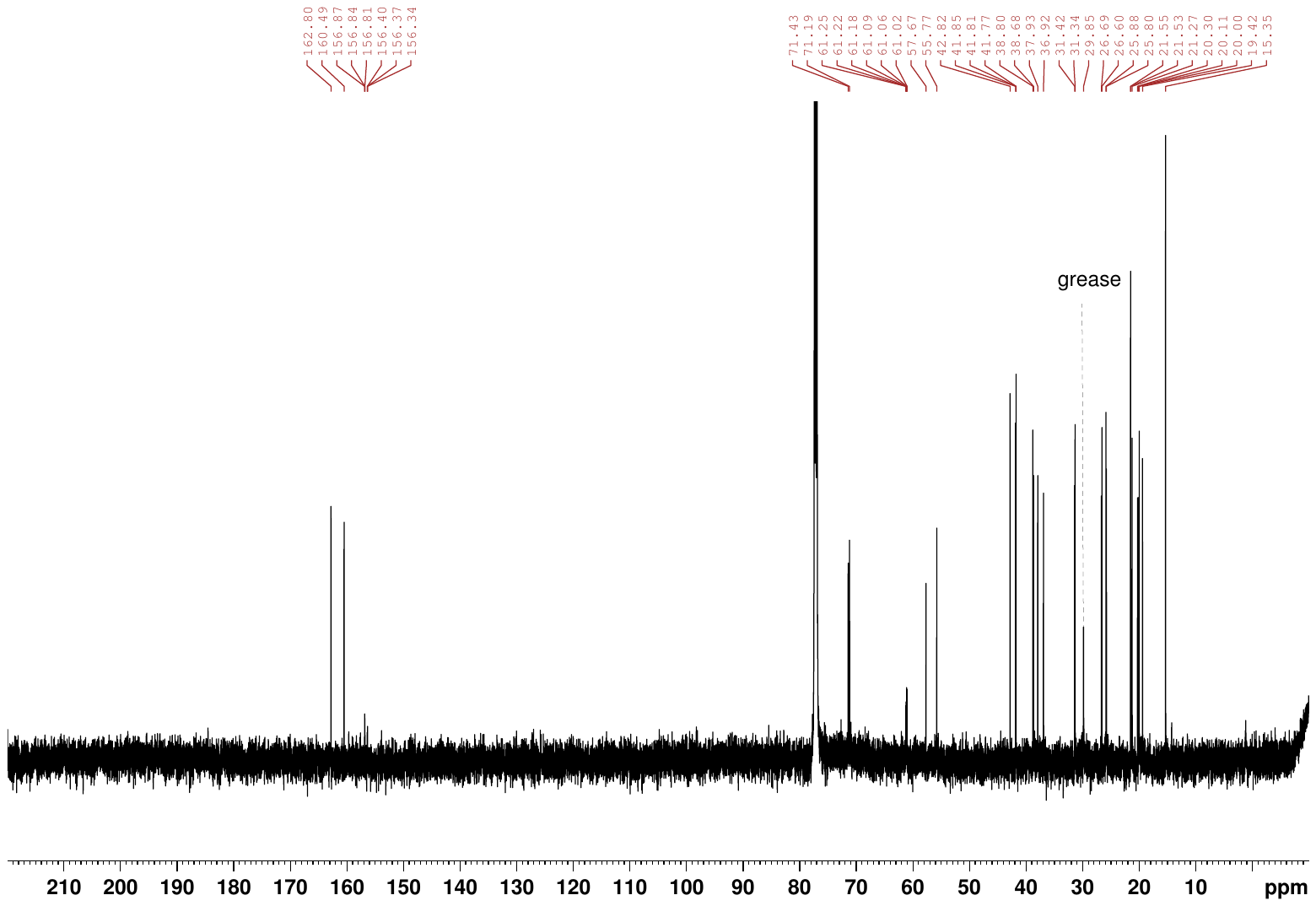

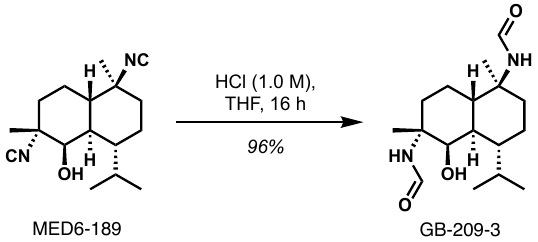

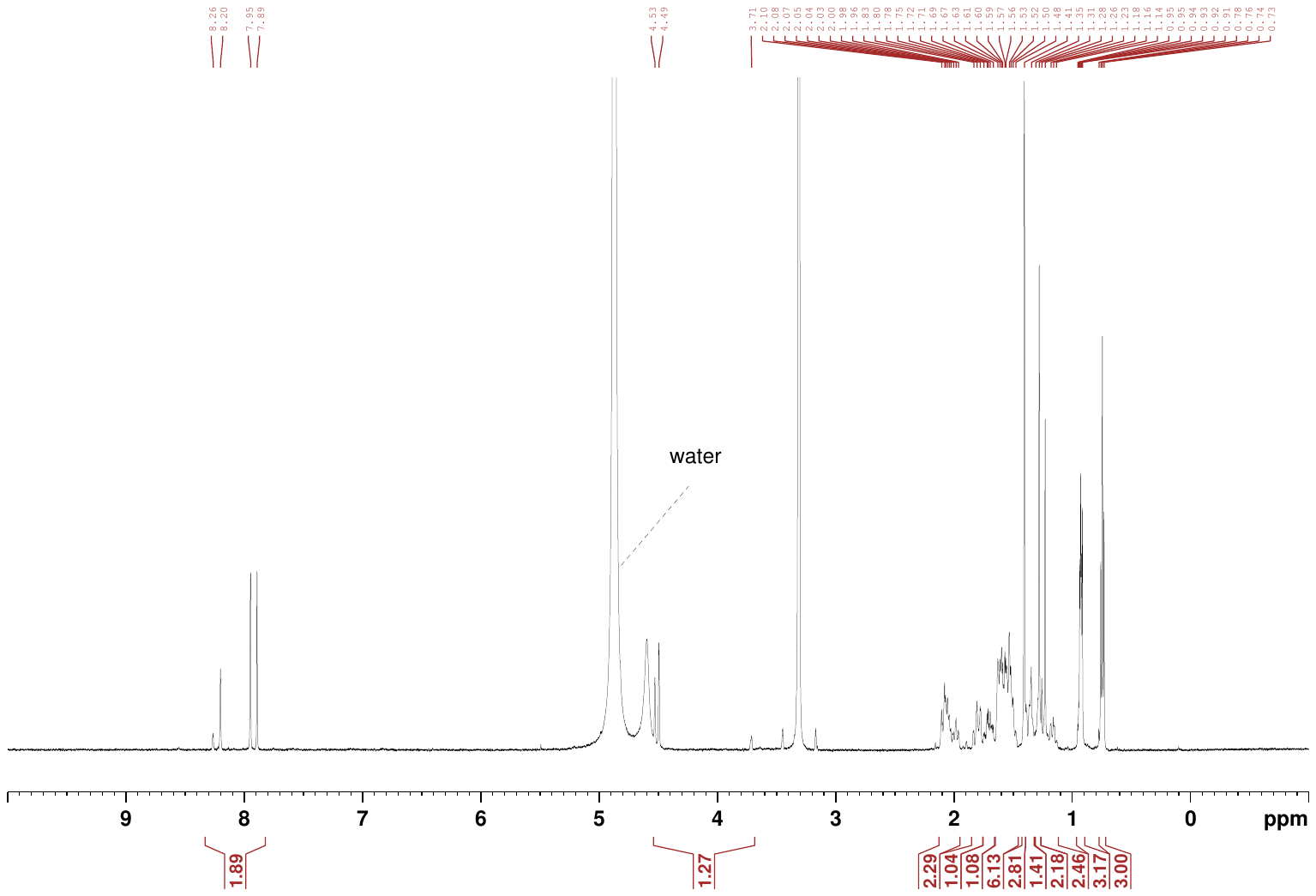

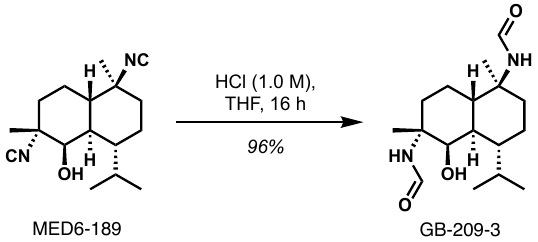

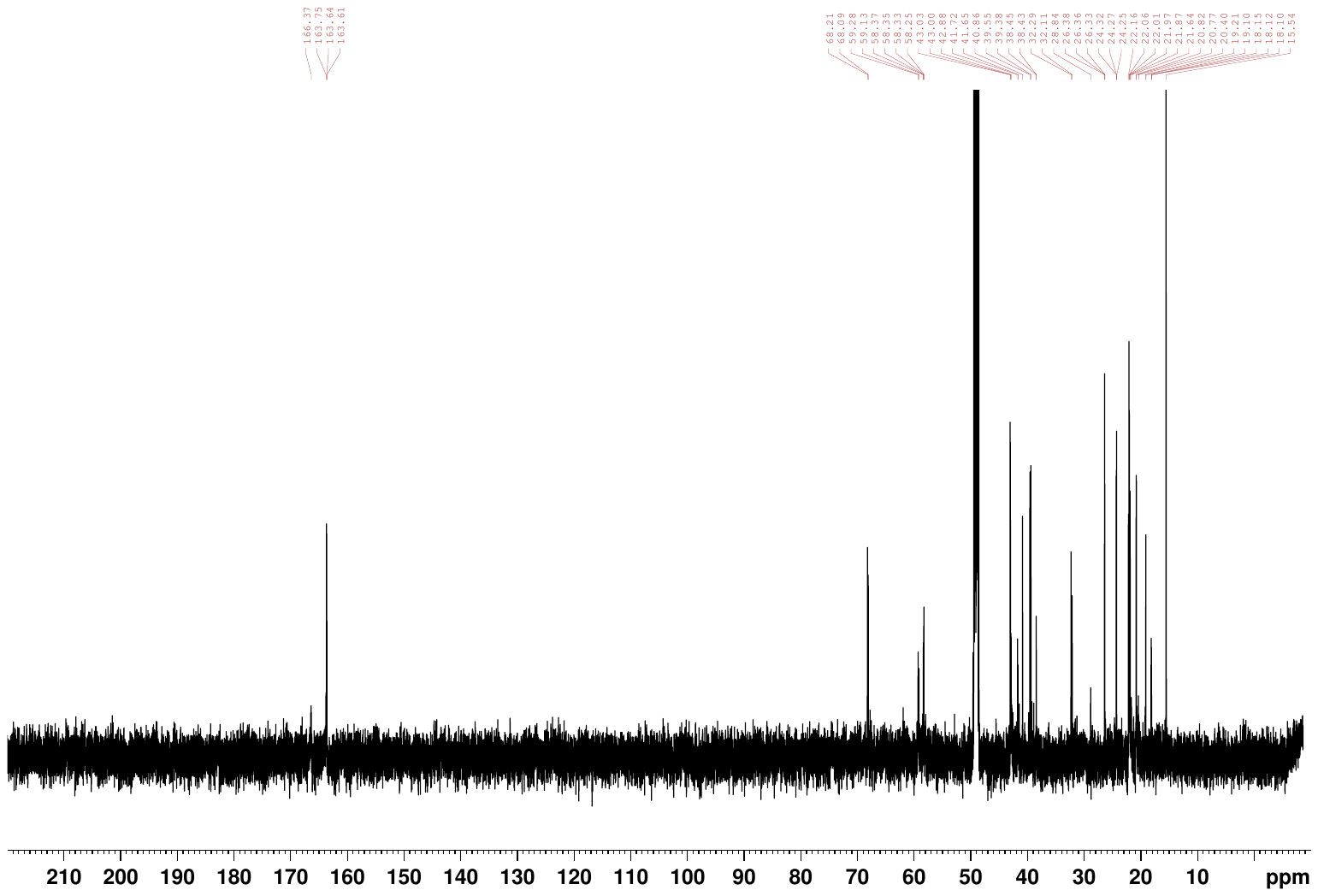
