## Supporting Information S2 for "A Potent Kalihinol Analogue Disrupts Apicoplast Function and Vesicular Trafficking in *P. falciparum* Malaria": Supporting Information_S2_CETSA_Curves_11_6_23.pdf

**CETSA curves plotted for heat challenged samples treated with DMSO (Controls) or MED6-189 (MED6)**

**CETSA curves are presented in order with decreasing euclidian distance (ED) scores for each condition:**

**MED6-189 (red) vs. DMSO (blue)**

Med6 CETSA data plotting_curve fitting

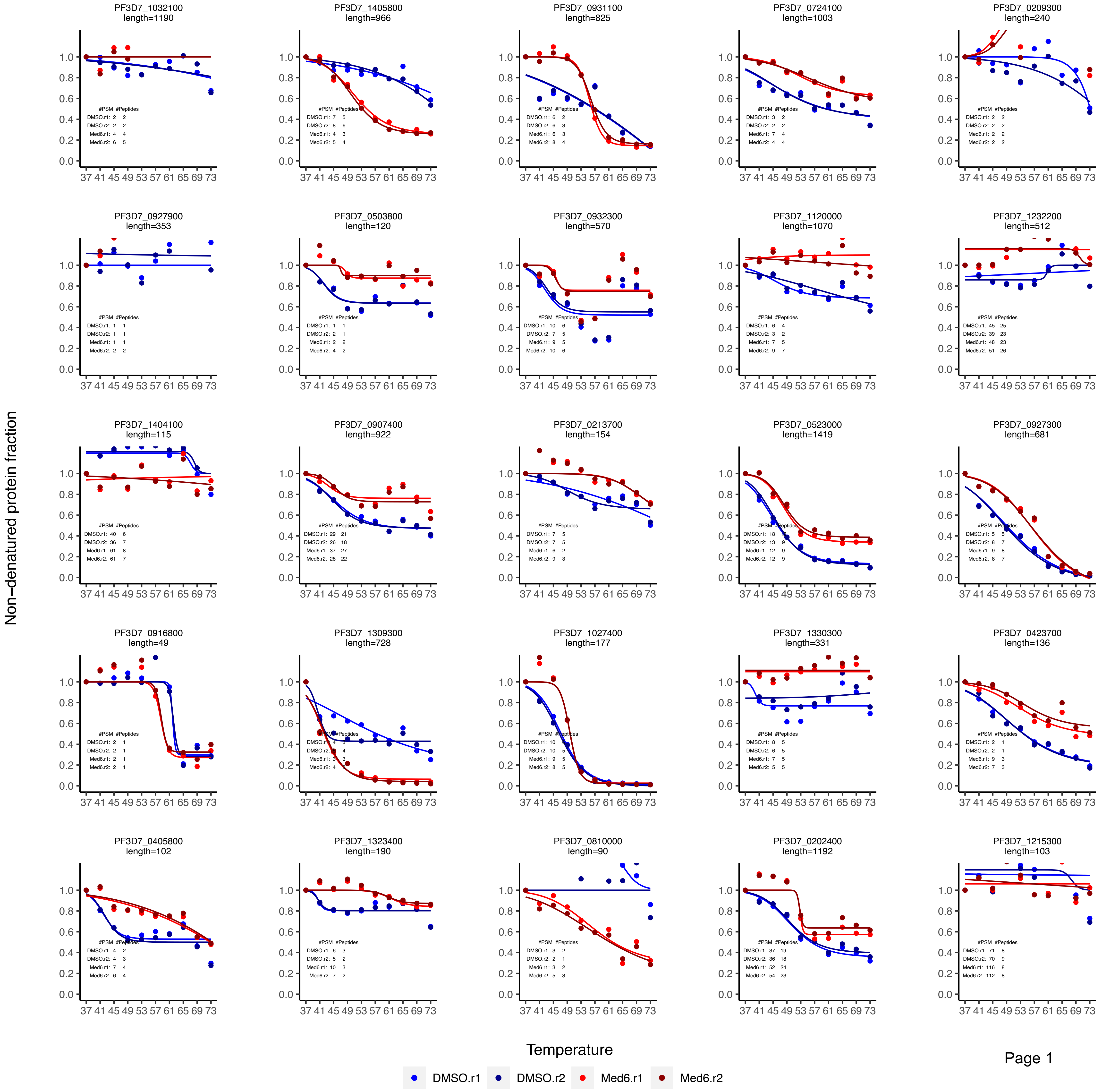

Med6 CETSA data plotting_curve fitting

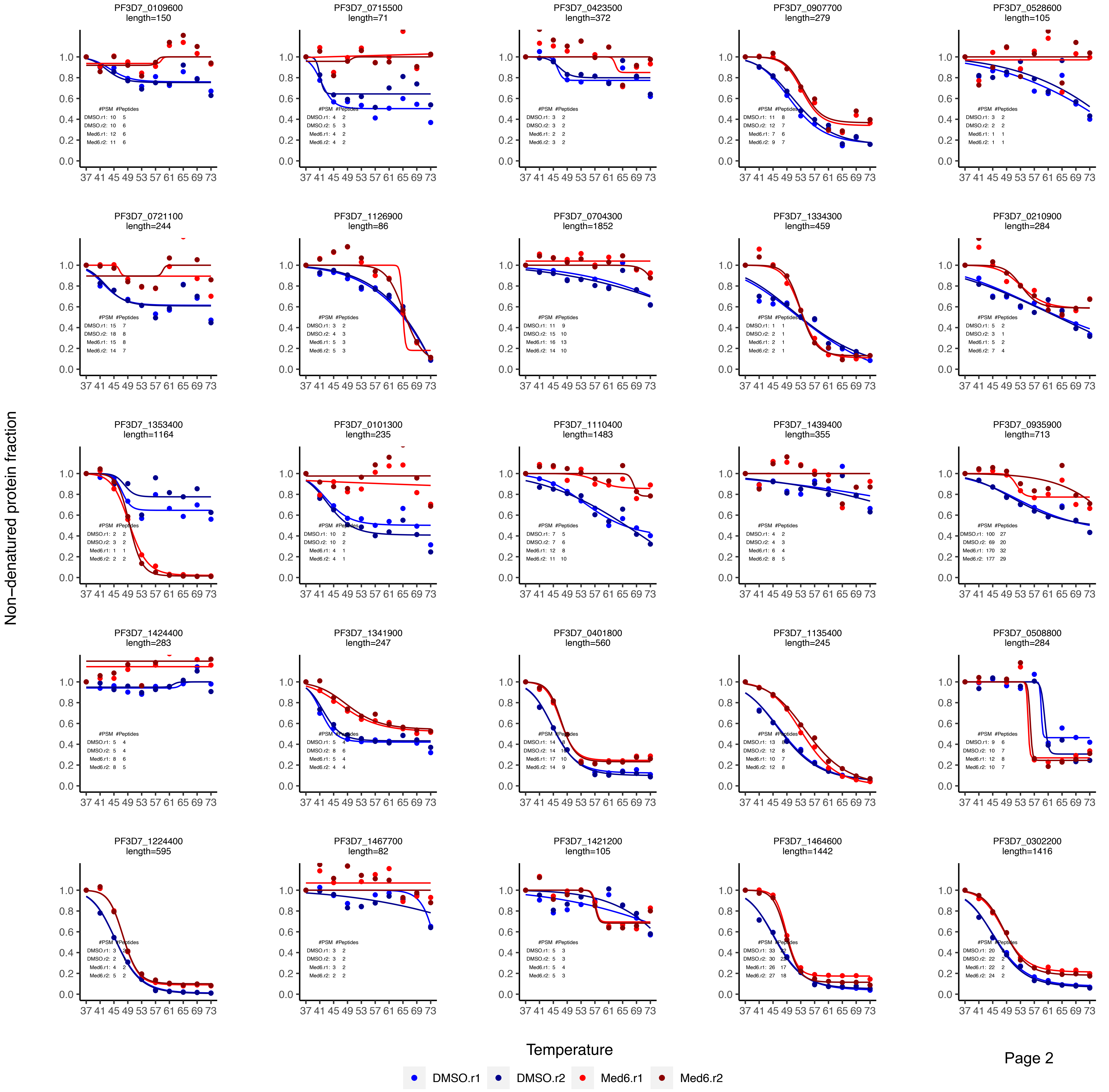


Page 2

Med6 CETSA data plotting_curve fitting

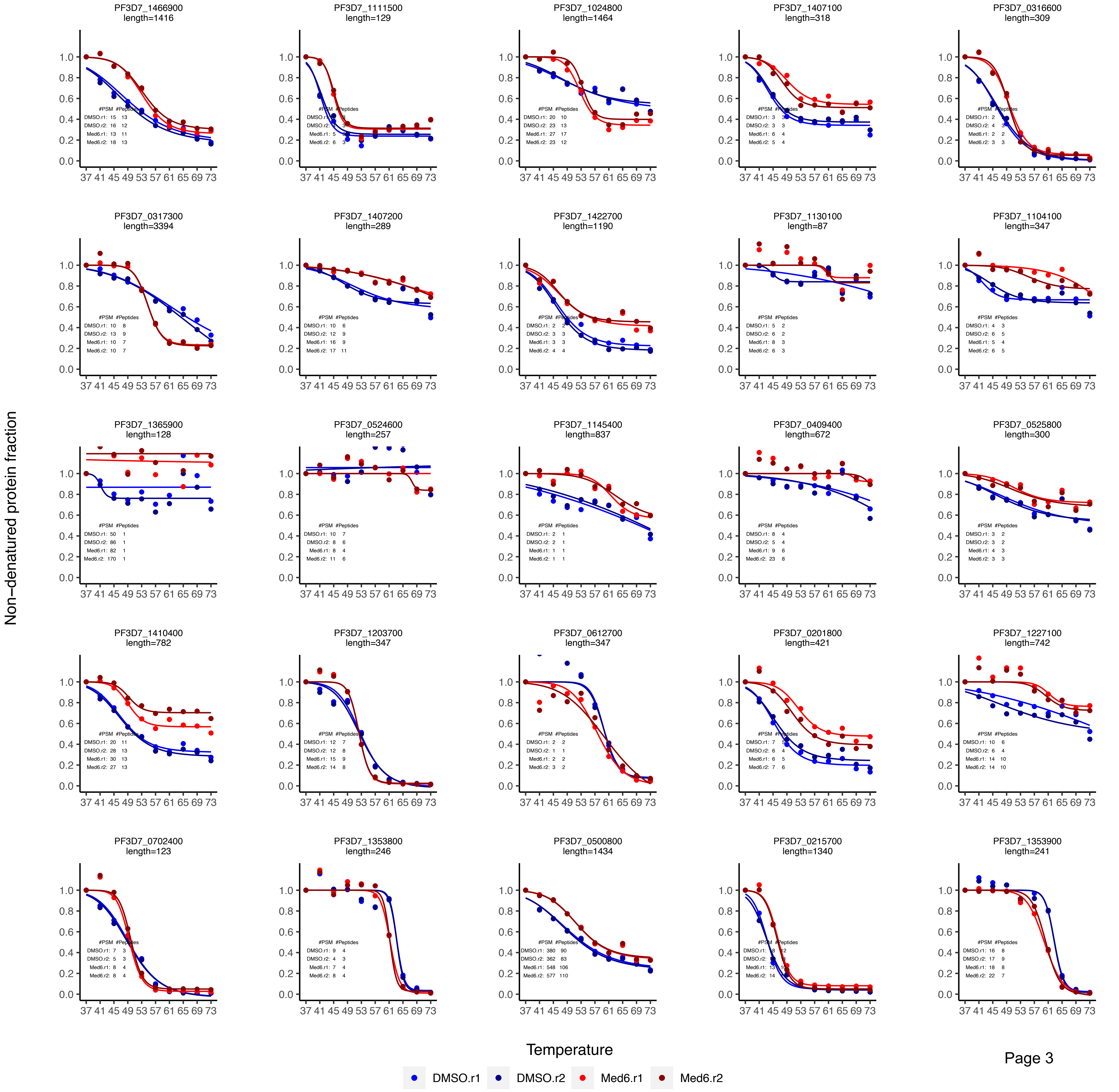

### Med6 CETSA data plotting_curve fitting

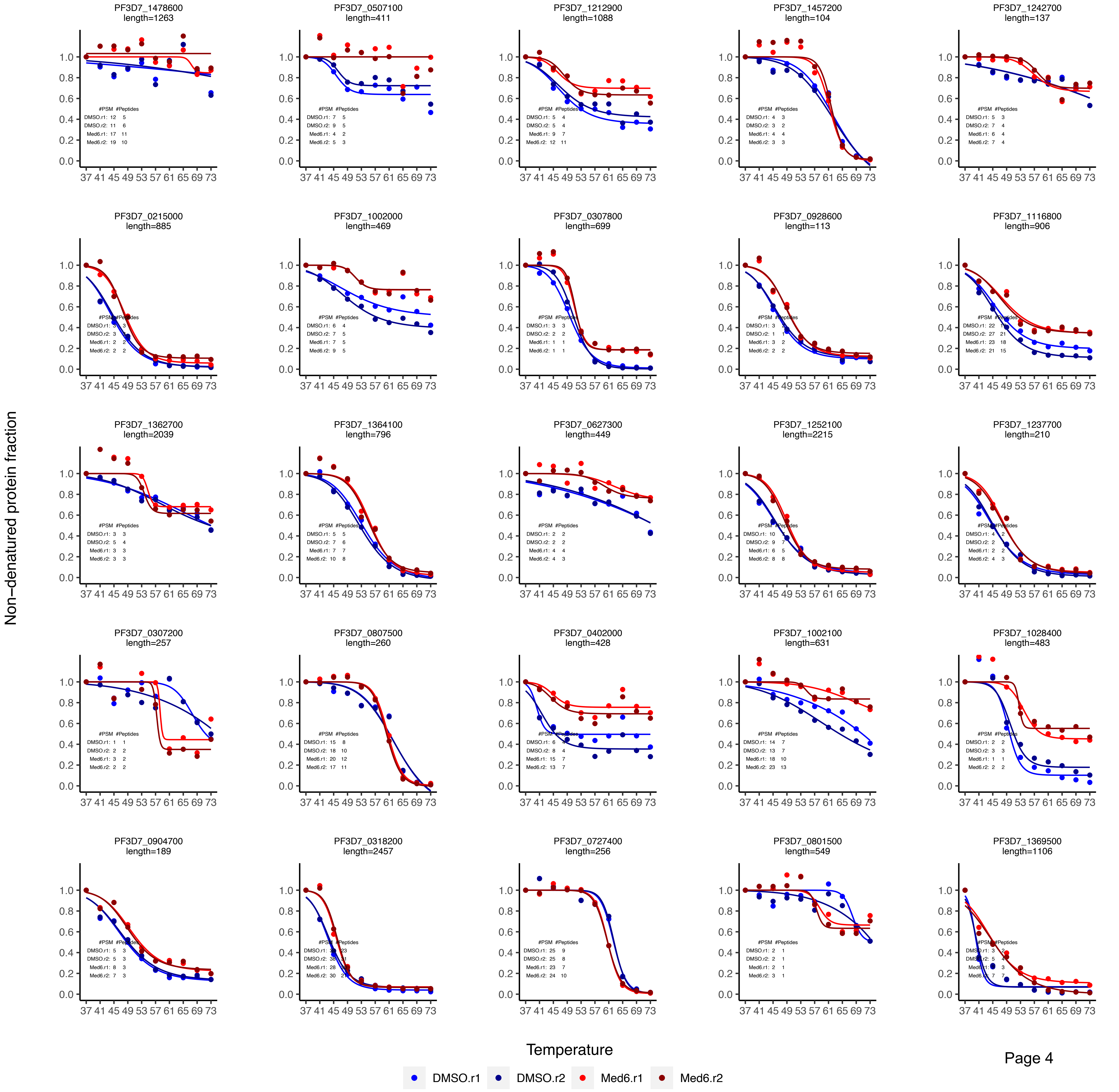

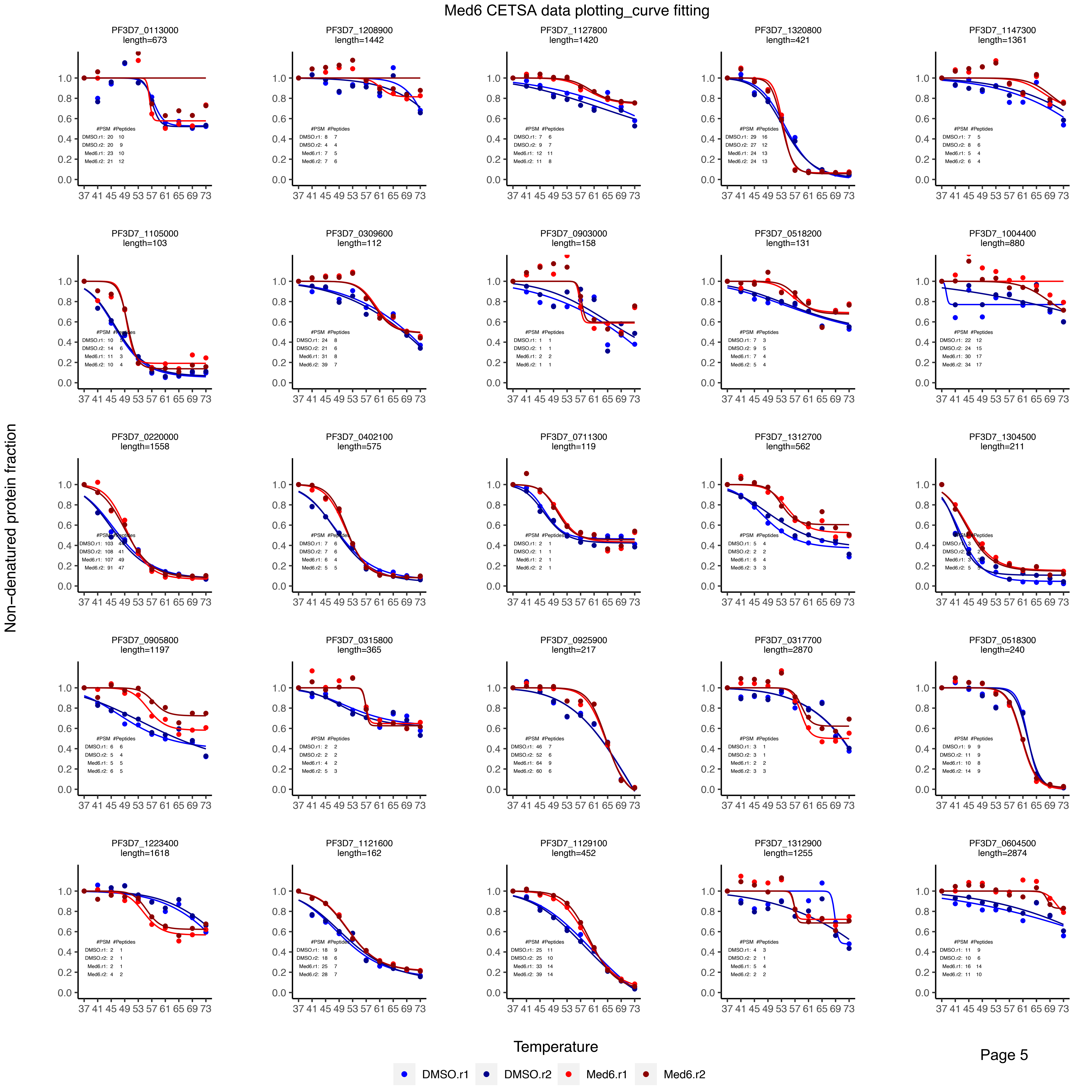

Med6 CETSA data plotting_curve fitting
PF3D7_0113000 length=673
#PSM #Peptides
DMSO.r1: 20 10
DMSO.r2: 20 9
Med6.r1: 23 10
Med6.r2: 21 12
PF3D7_1208900 length=1442
#PSM #Peptides
DMSO.r1: 8 7
DMSO.r2: 4 4
Med6.r1: 7 5
Med6.r2: 7 6
PF3D7_1127800 length=1420
#PSM #Peptides
DMSO.r1: 7 6
DMSO.r2: 9 7
Med6.r1: 12 11
Med6.r2: 11 8
PF3D7_1320800 length=421
#PSM #Peptides
DMSO.r1: 29 16
DMSO.r2: 27 12
Med6.r1: 24 13
Med6.r2: 24 13
PF3D7_1147300 length=1361
#PSM #Peptides
DMSO.r1: 7 5
DMSO.r2: 8 6
Med6.r1: 5 4
Med6.r2: 6 4
PF3D7_1105000 length=103
PF3D7_0309600 length=112
#PSM #Peptides
DMSO.r1: 24 8
DMSO.r2: 21 6
Med6.r1: 31 8
Med6.r2: 39 7
PF3D7_0903000 length=158
#PSM #Peptides
DMSO.r1: 1 1
DMSO.r2: 1 1
Med6.r1: 2 2
Med6.r2: 1 1
PF3D7_0518200 length=131
#PSM #Peptides
DMSO.r1: 7 3
DMSO.r2: 9 5
Med6.r1: 7 4
Med6.r2: 5 4
PF3D7_1004400 length=880
#PSM #Peptides
DMSO.r1: 22 12
DMSO.r2: 24 15
Med6.r1: 30 17
Med6.r2: 34 17
PF3D7_0220000 length=1558
PF3D7_0402100 length=575
PF3D7_0711300 length=119
PF3D7_1312700 length=562
#PSM #Peptides
DMSO.r1: 5 4
DMSO.r2: 2 2
Med6.r1: 6 4
Med6.r2: 3 3
PF3D7_1304500 length=211
PF3D7_0905800 length=1197
#PSM #Peptides
DMSO.r1: 6 6
DMSO.r2: 5 4
Med6.r1: 5 5
Med6.r2: 6 5
PF3D7_0315800 length=365
#PSM #Peptides
DMSO.r1: 2 2
DMSO.r2: 2 2
Med6.r1: 4 2
Med6.r2: 5 3
PF3D7_0925900 length=217
#PSM #Peptides
DMSO.r1: 46 7
DMSO.r2: 52 6
Med6.r1: 64 9
Med6.r2: 60 6
PF3D7_0317700 length=2870
#PSM #Peptides
DMSO.r1: 3 1
DMSO.r2: 3 1
Med6.r1: 2 2
Med6.r2: 3 3
PF3D7_0518300 length=240
#PSM #Peptides
DMSO.r1: 9 9
DMSO.r2: 11 9
Med6.r1: 10 8
Med6.r2: 14 9
PF3D7_1223400 length=1618
#PSM #Peptides
DMSO.r1: 2 1
DMSO.r2: 2 1
Med6.r1: 2 1
Med6.r2: 4 2
PF3D7_1121600 length=162
PF3D7_1129100 length=452
#PSM #Peptides
DMSO.r1: 25 11
DMSO.r2: 25 10
Med6.r1: 33 14
Med6.r2: 39 14
PF3D7_1312900 length=1255
#PSM #Peptides
DMSO.r1: 4 3
DMSO.r2: 2 1
Med6.r1: 5 4
Med6.r2: 2 2
PF3D7_0604500 length=2874
#PSM #Peptides
DMSO.r1: 11 9
DMSO.r2: 10 6
Med6.r1: 16 14
Med6.r2: 11 10
0.0 0.2 0.4 0.6 0.8 1.0
37 41 45 49 53 57 61 65 69 73
Non−denatured protein fraction
Temperature
Page 5
DMSO.r1 DMSO.r2 Med6.r1 Med6.r2

Med6 CETSA data plotting_curve fitting

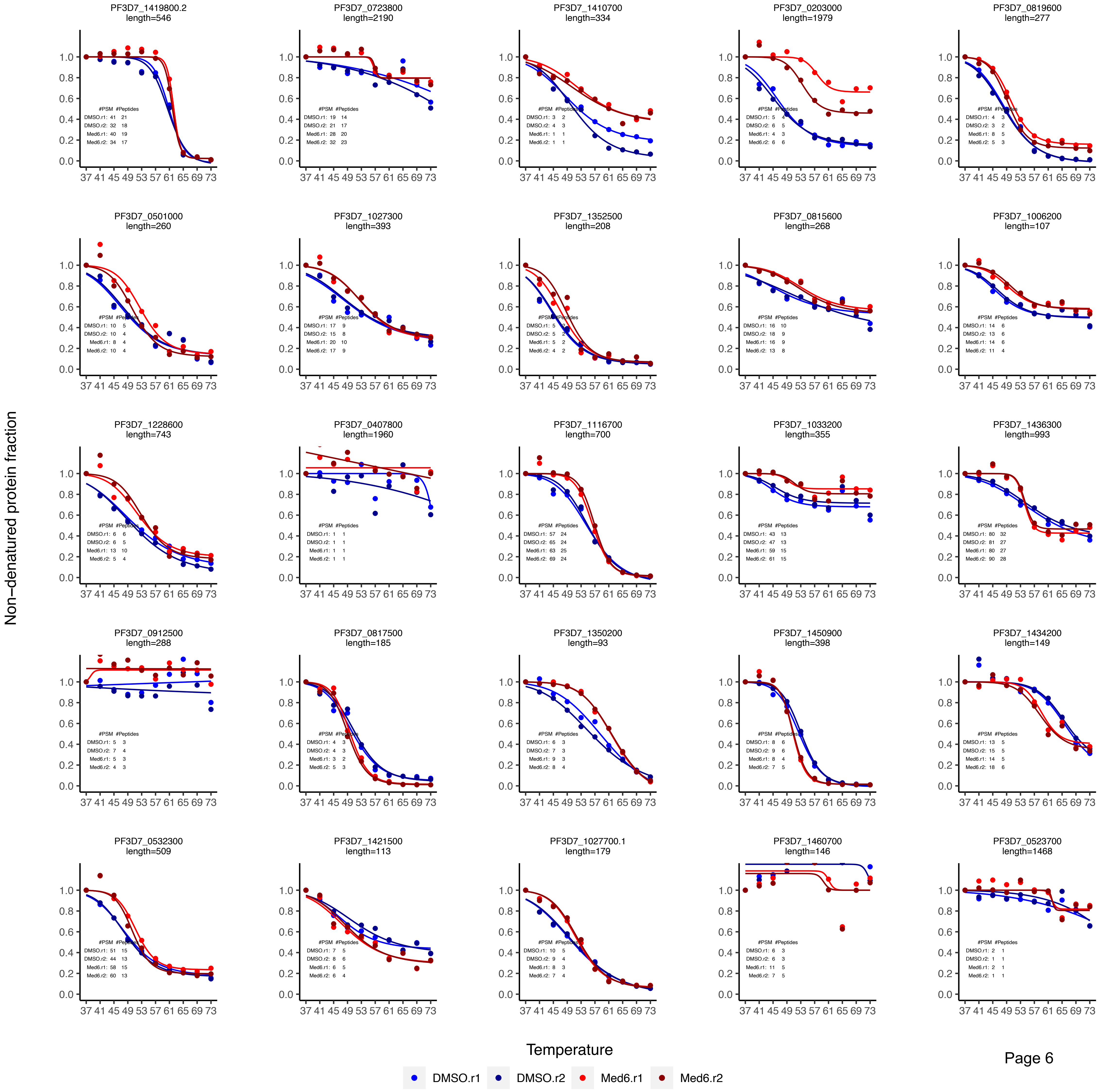

Med6 CETSA data plotting_curve fitting

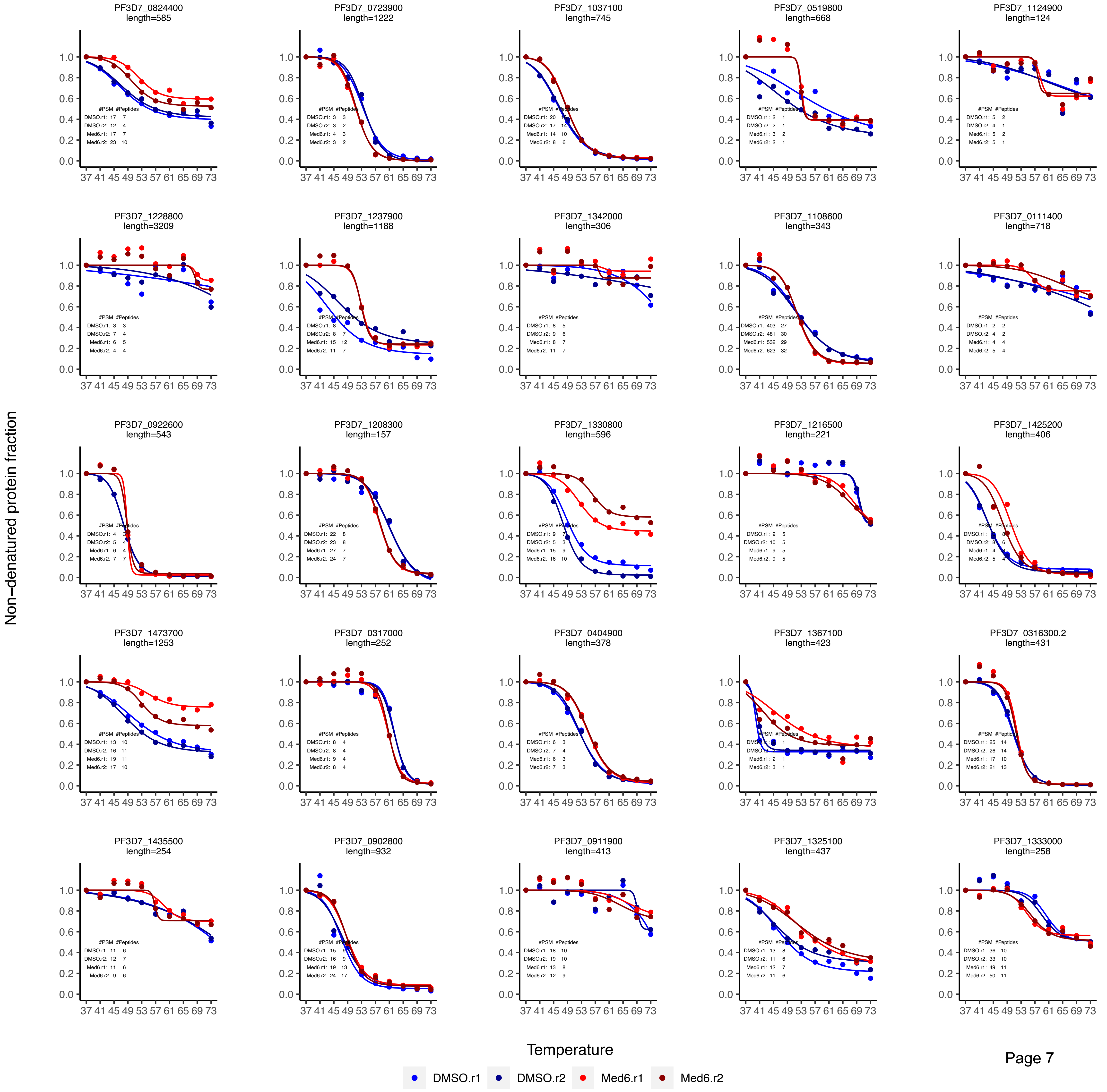

Med6 CETSA data plotting_curve fitting

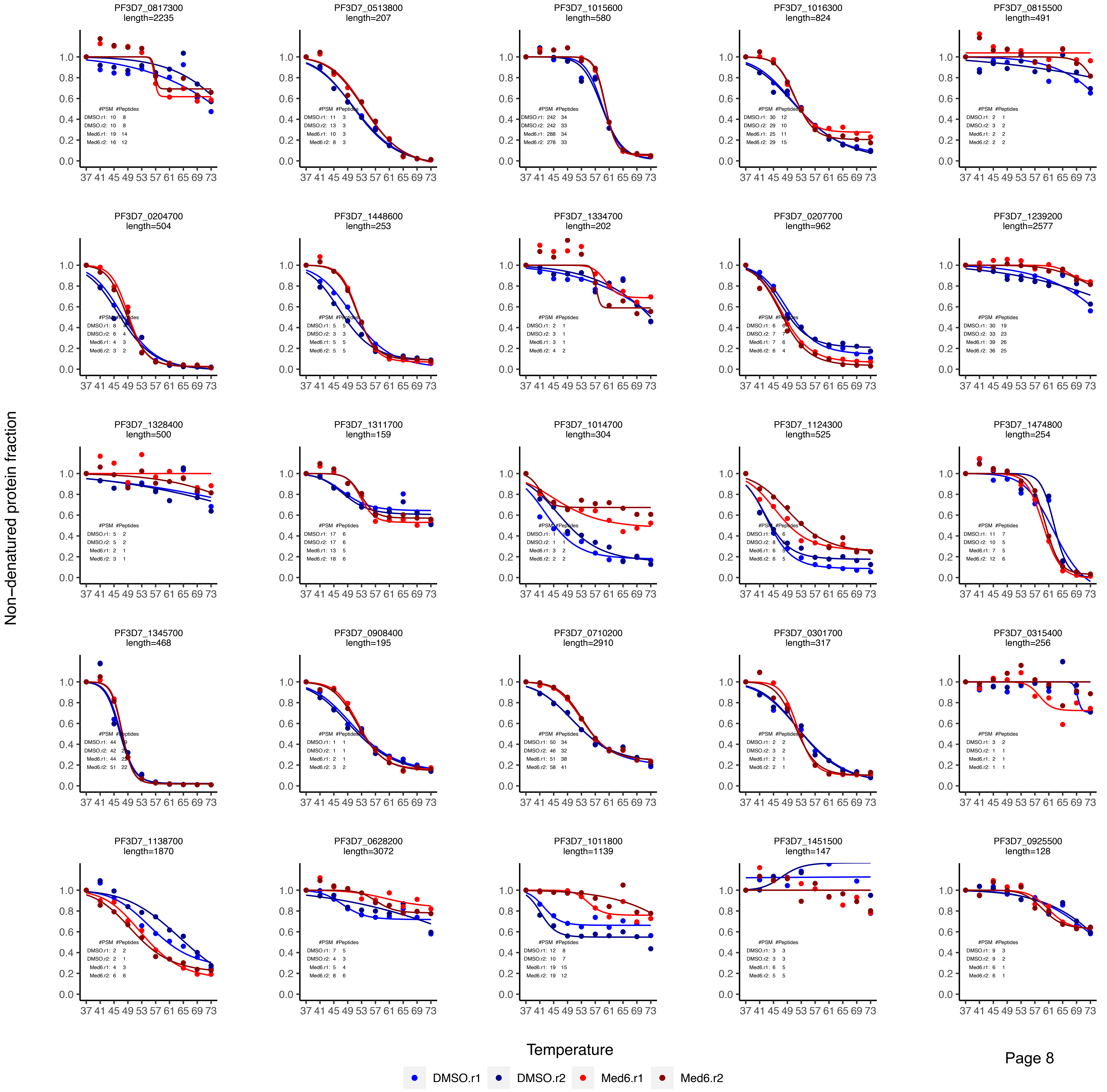

Med6 CETSA data plotting_curve fitting

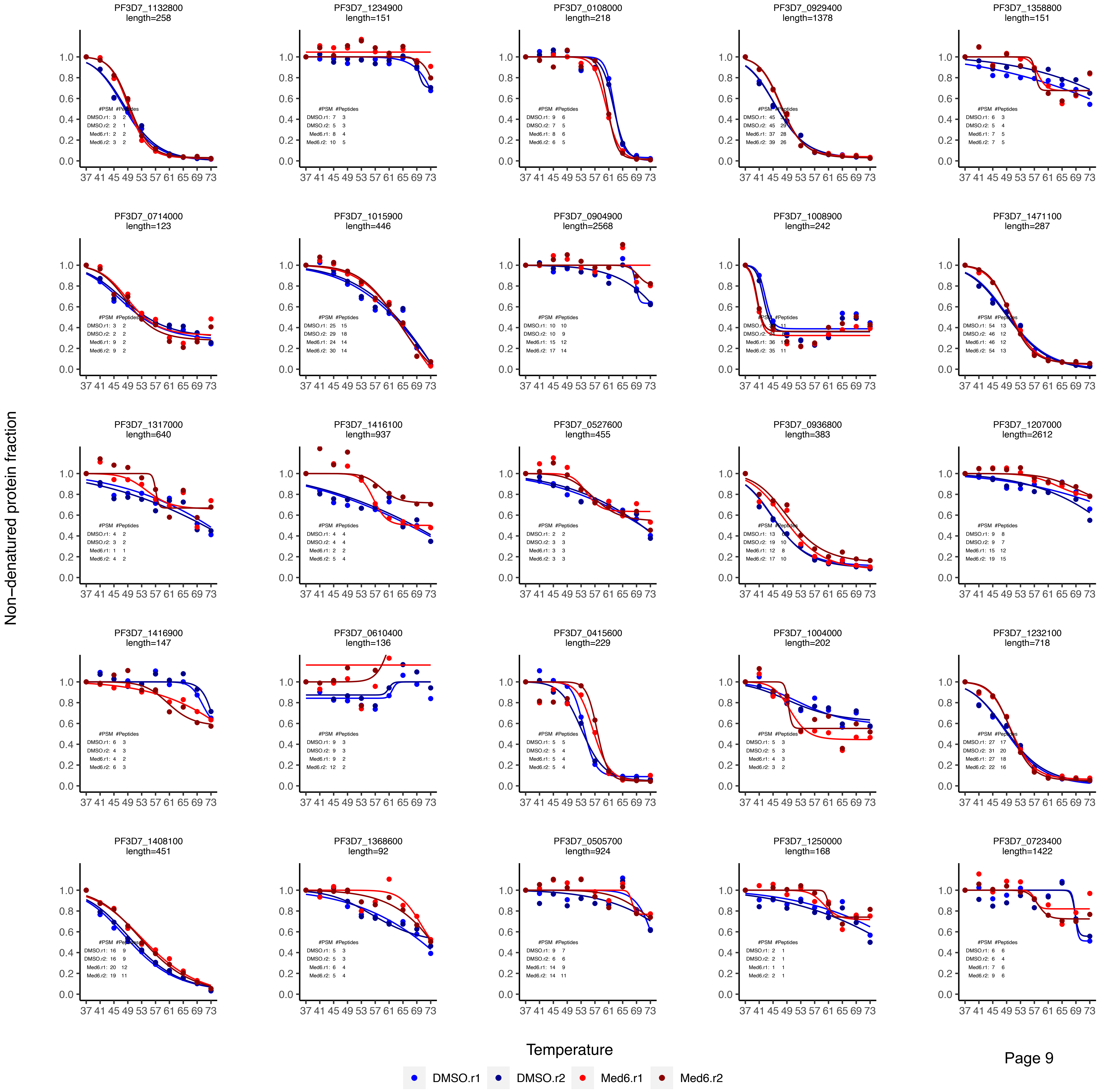

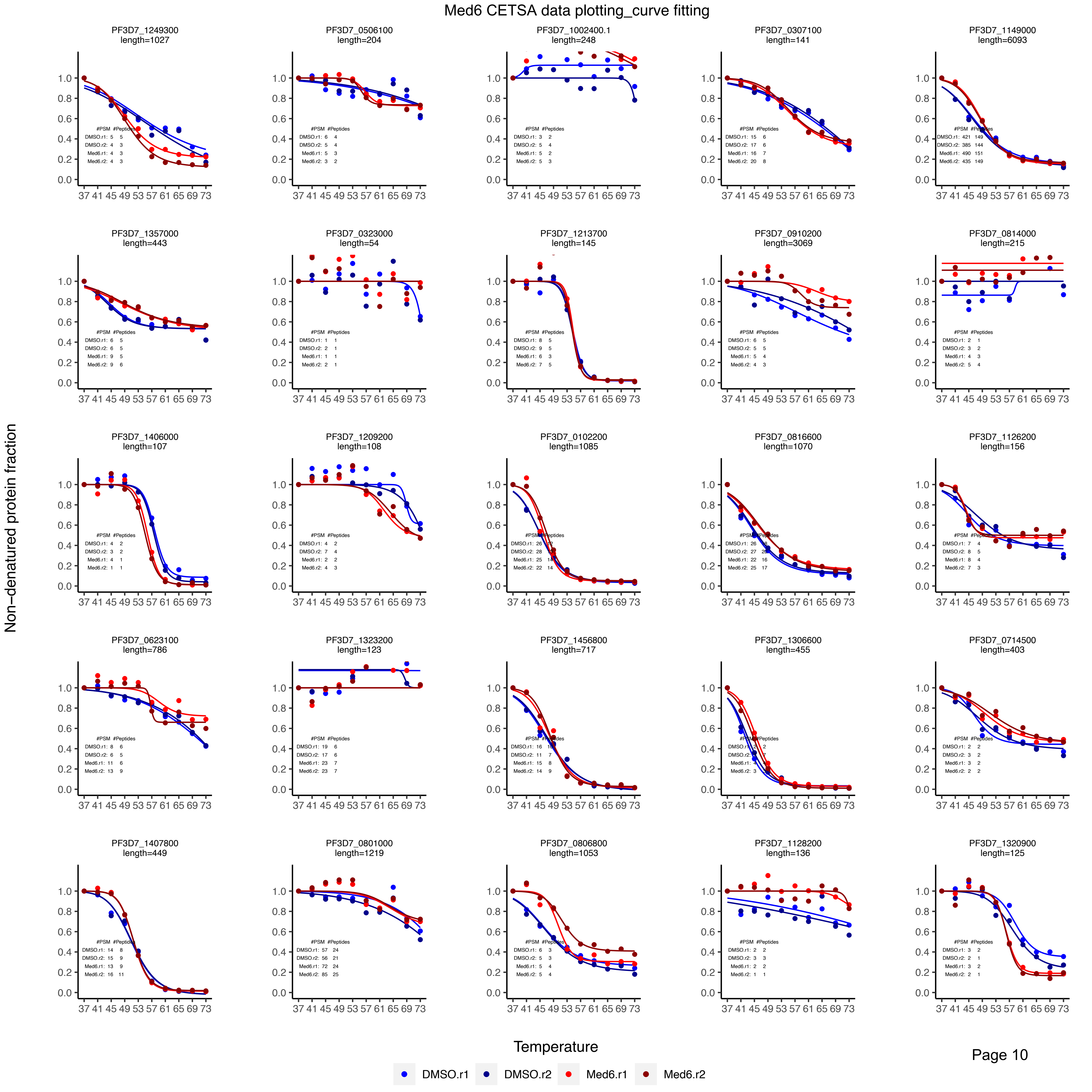
Med6 CETSA data plotting_curve fitting
PF3D7_1249300 length=1027
PF3D7_0506100 length=204
PF3D7_1002400.1 length=248
PF3D7_0307100 length=141
PF3D7_1149000 length=6093
PF3D7_1357000 length=443
PF3D7_0323000 length=54
PF3D7_1213700 length=145
PF3D7_0910200 length=3069
PF3D7_0814000 length=215
PF3D7_1406000 length=107
PF3D7_1209200 length=108
PF3D7_0102200 length=1085
PF3D7_0816600 length=1070
PF3D7_1126200 length=156
PF3D7_0623100 length=786
PF3D7_1323200 length=123
PF3D7_1456800 length=717
PF3D7_1306600 length=455
PF3D7_0714500 length=403
PF3D7_1407800 length=449
PF3D7_0801000 length=1219
PF3D7_0806800 length=1053
PF3D7_1128200 length=136
PF3D7_1320900 length=125
Non-denatured protein fraction
Temperature
DMSO.r1
DMSO.r2
Med6.r1
Med6.r2
Page 10

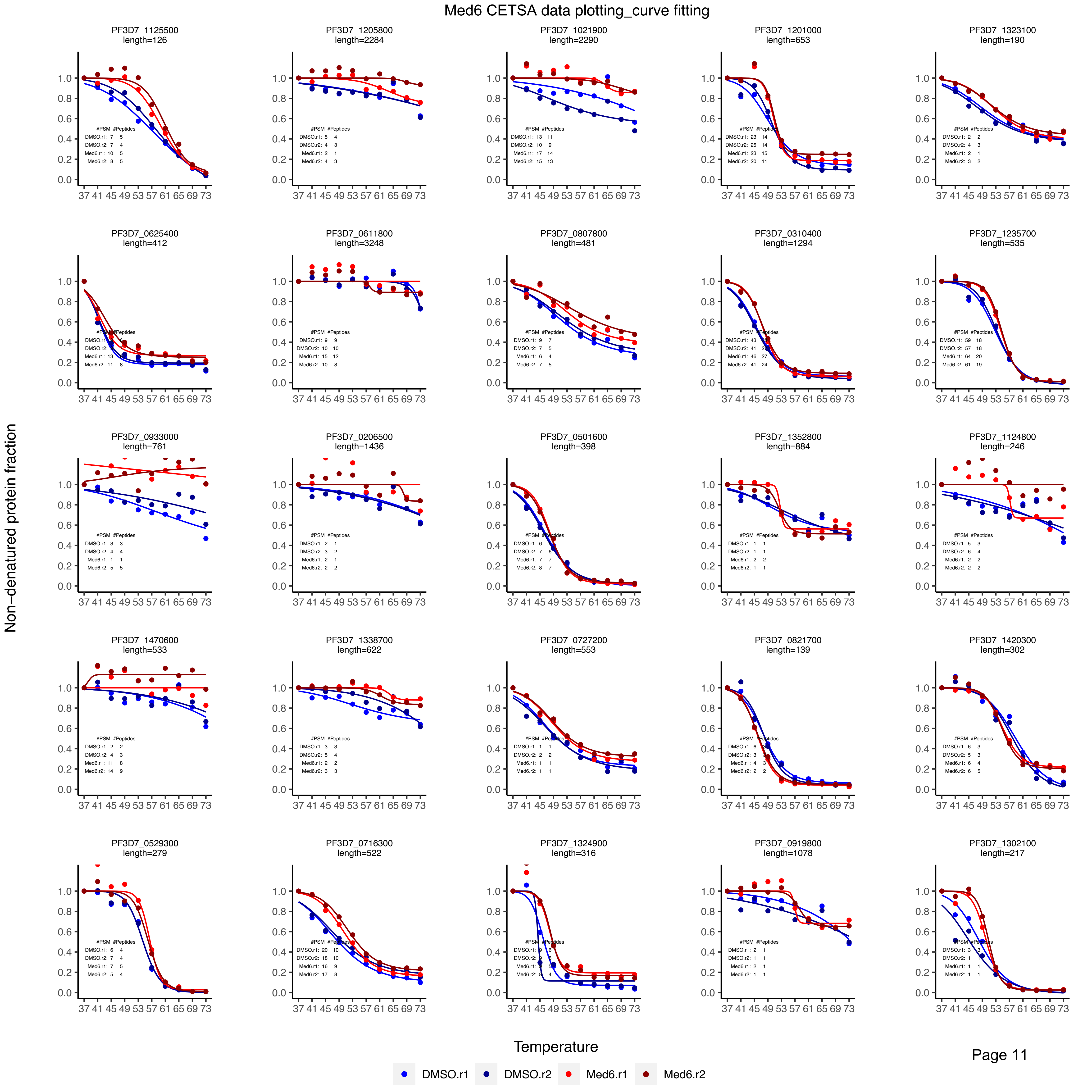

Med6 CETSA data plotting_curve fitting
PF3D7_1125500 length=126
PF3D7_1205800 length=2284
PF3D7_1021900 length=2290
PF3D7_1201000 length=653
PF3D7_1323100 length=190
PF3D7_0625400 length=412
PF3D7_0611800 length=3248
PF3D7_0807800 length=481
PF3D7_0310400 length=1294
PF3D7_1235700 length=535
PF3D7_0933000 length=761
PF3D7_0206500 length=1436
PF3D7_0501600 length=398
PF3D7_1352800 length=884
PF3D7_1124800 length=246
PF3D7_1470600 length=533
PF3D7_1338700 length=622
PF3D7_0727200 length=553
PF3D7_0821700 length=139
PF3D7_1420300 length=302
PF3D7_0529300 length=279
PF3D7_0716300 length=522
PF3D7_1324900 length=316
PF3D7_0919800 length=1078
PF3D7_1302100 length=217
Non−denatured protein fraction
Temperature
Page 11
DMSO.r1
DMSO.r2
Med6.r1
Med6.r2

Med6 CETSA data plotting_curve fitting

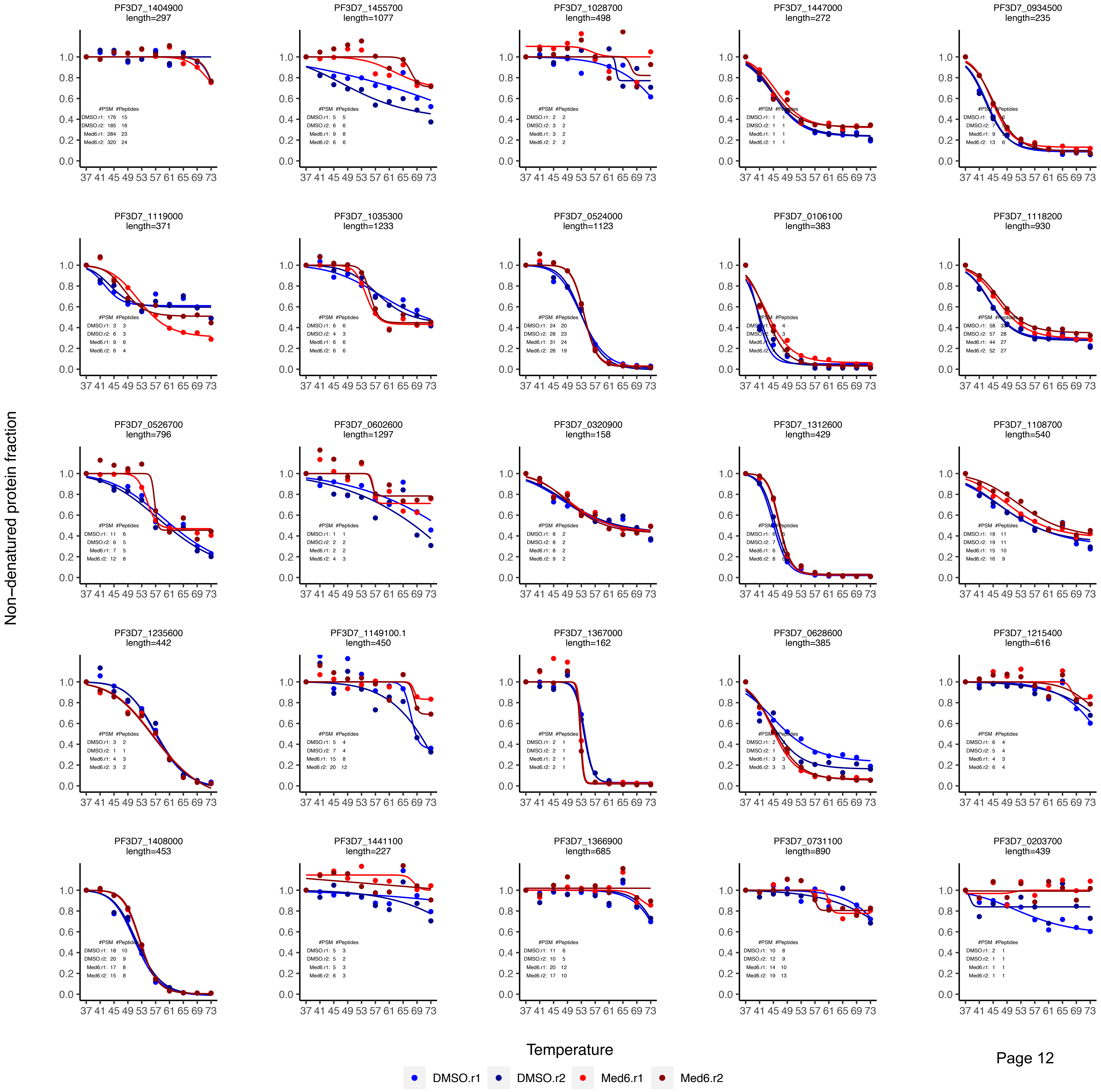

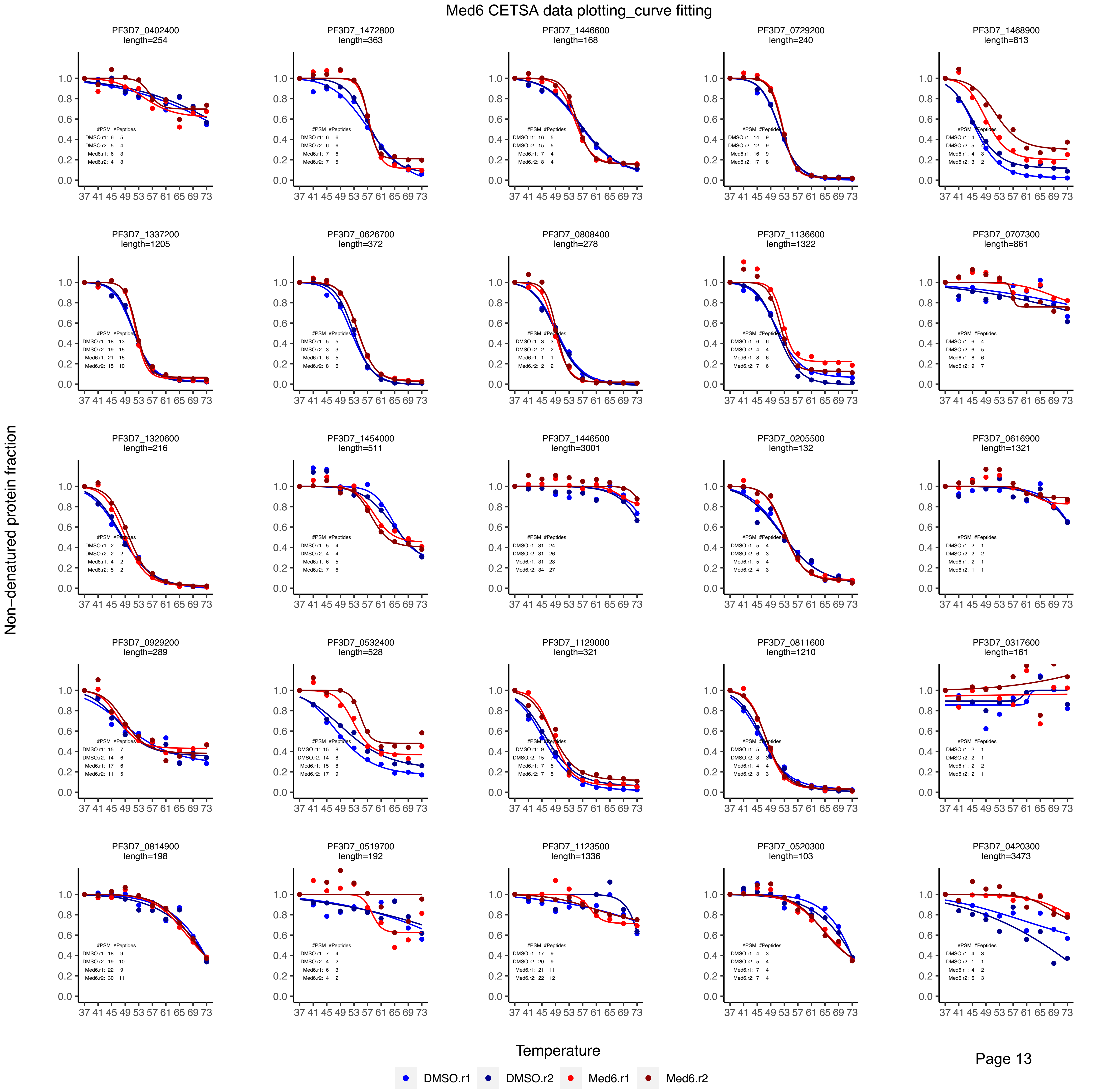
Med6 CETSA data plotting_curve fitting
PF3D7_0402400 length=254
PF3D7_1472800 length=363
PF3D7_1446600 length=168
PF3D7_0729200 length=240
PF3D7_1468900 length=813
PF3D7_1337200 length=1205
PF3D7_0626700 length=372
PF3D7_0808400 length=278
PF3D7_1136600 length=1322
PF3D7_0707300 length=861
PF3D7_1320600 length=216
PF3D7_1454000 length=511
PF3D7_1446500 length=3001
PF3D7_0205500 length=132
PF3D7_0616900 length=1321
PF3D7_0929200 length=289
PF3D7_0532400 length=528
PF3D7_1129000 length=321
PF3D7_0811600 length=1210
PF3D7_0317600 length=161
PF3D7_0814900 length=198
PF3D7_0519700 length=192
PF3D7_1123500 length=1336
PF3D7_0520300 length=103
PF3D7_0420300 length=3473
Non-denatured protein fraction
Temperature
DMSO.r1
DMSO.r2
Med6.r1
Med6.r2

Med6 CETSA data plotting_curve fitting

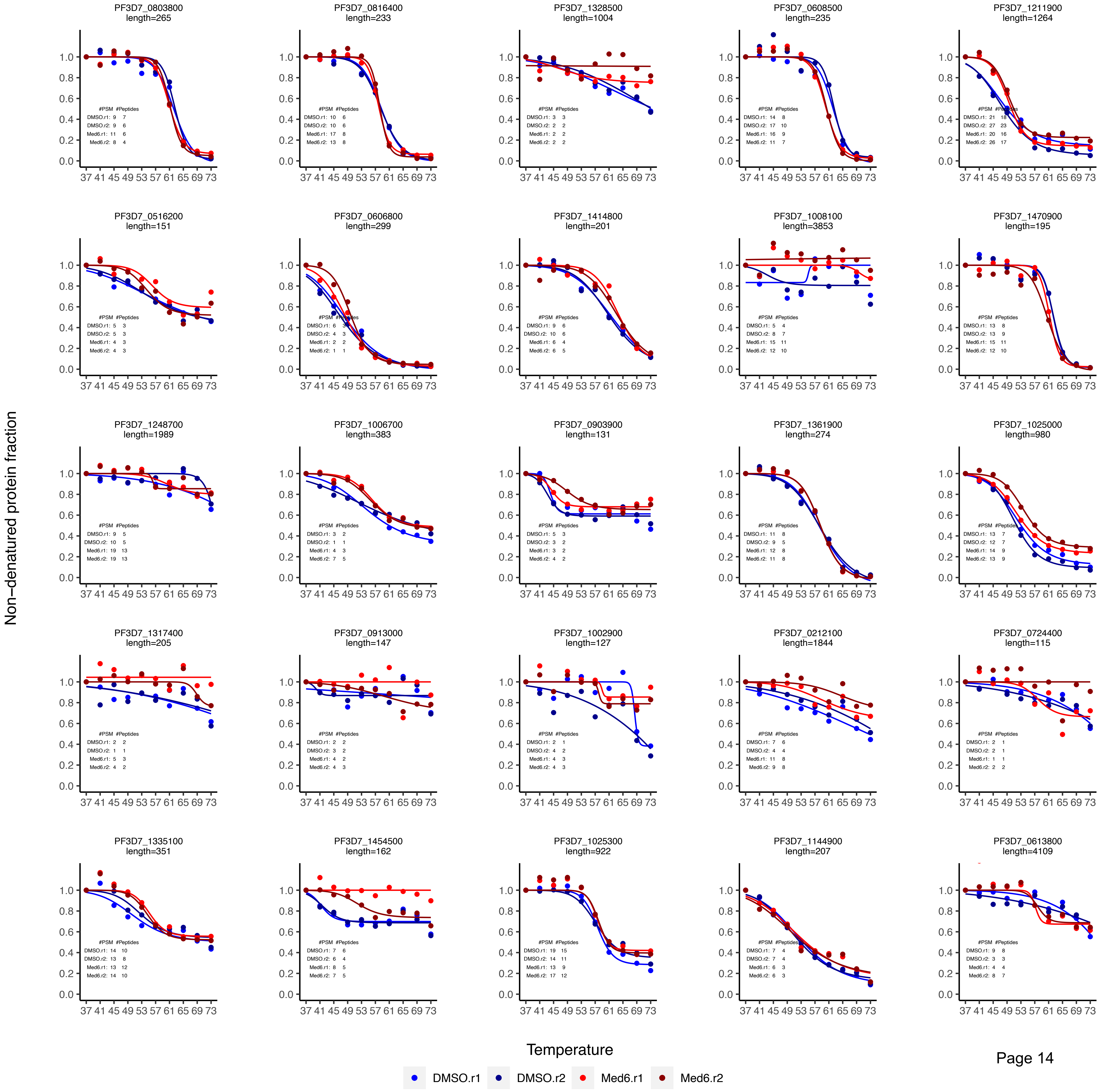

### Med6 CETSA data plotting_curve fitting

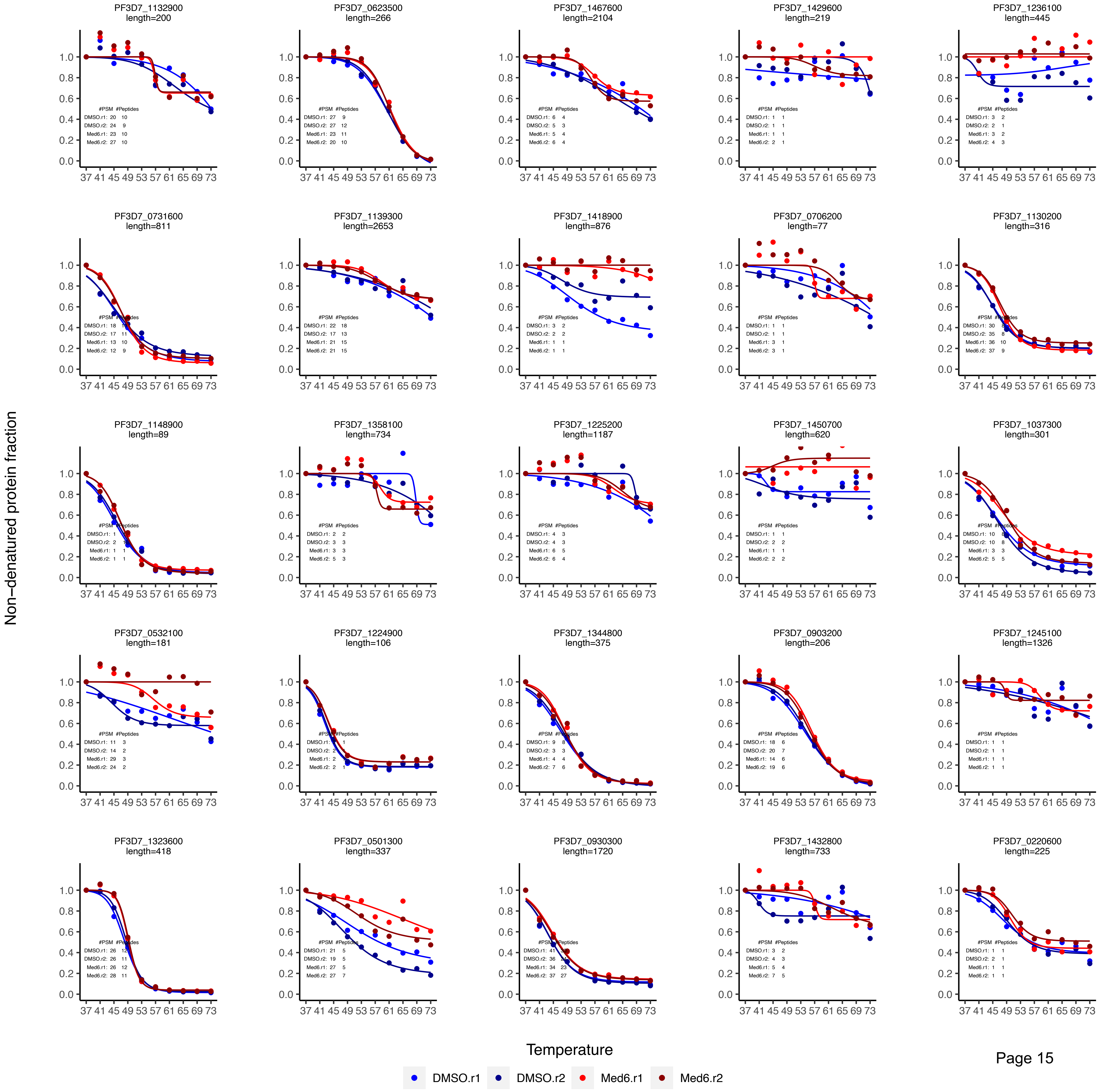

Med6 CETSA data plotting_curve fitting

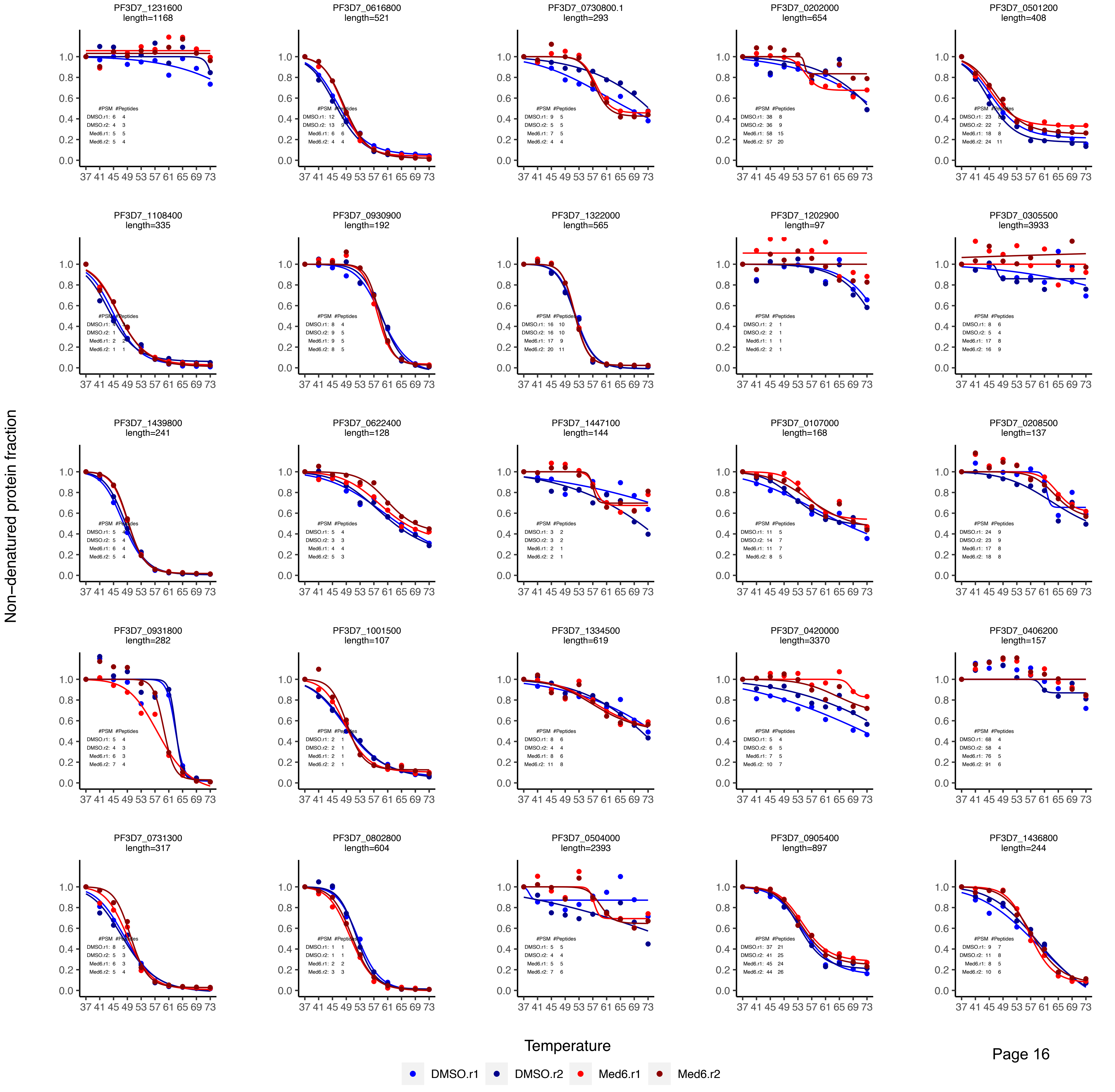

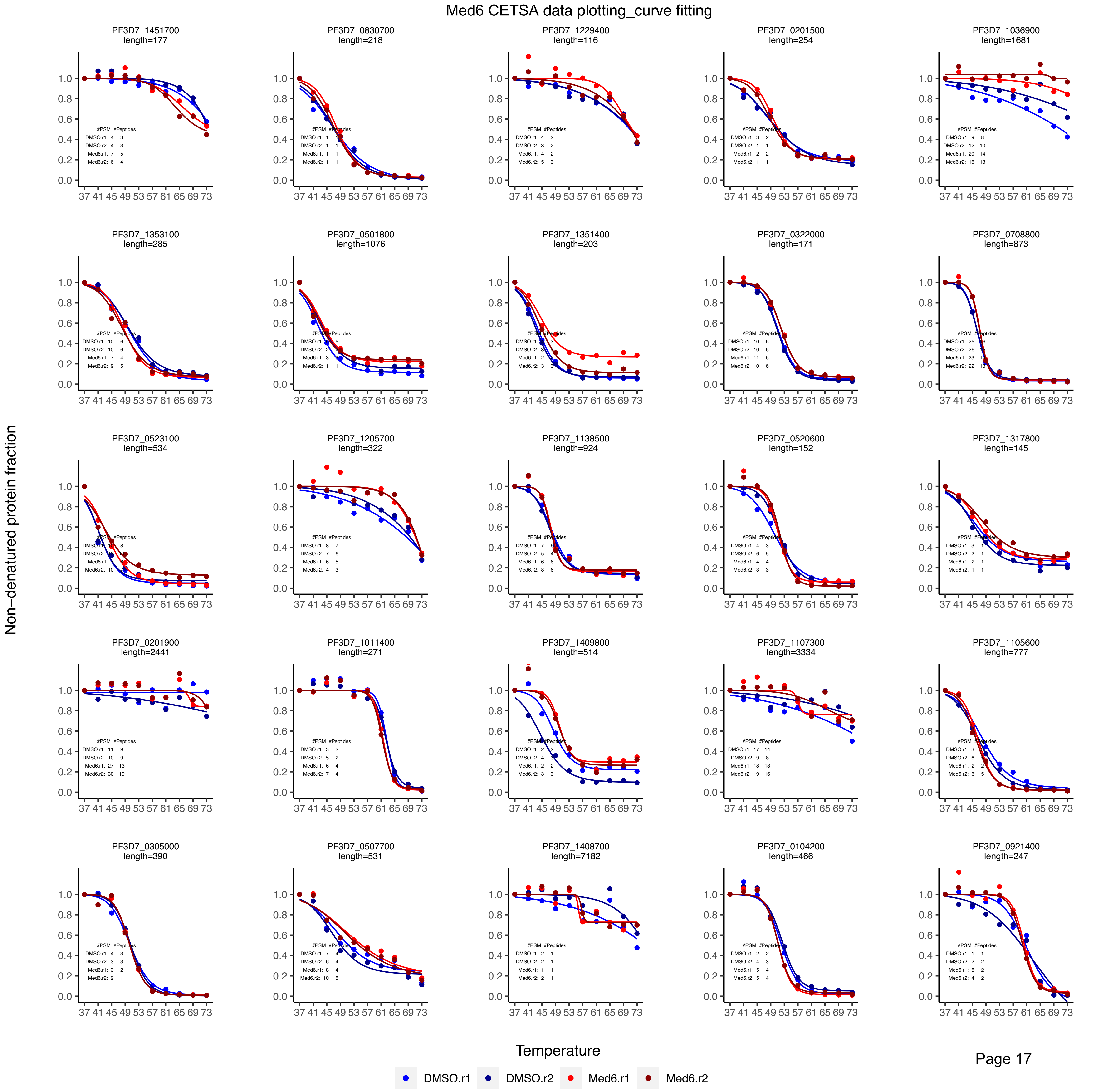
Med6 CETSA data plotting_curve fitting
PF3D7_1451700 length=177
PF3D7_0830700 length=218
PF3D7_1229400 length=116
PF3D7_0201500 length=254
PF3D7_1036900 length=1681
PF3D7_1353100 length=285
PF3D7_0501800 length=1076
PF3D7_1351400 length=203
PF3D7_0322000 length=171
PF3D7_0708800 length=873
PF3D7_0523100 length=534
PF3D7_1205700 length=322
PF3D7_1138500 length=924
PF3D7_0520600 length=152
PF3D7_1317800 length=145
PF3D7_0201900 length=2441
PF3D7_1011400 length=271
PF3D7_1409800 length=514
PF3D7_1107300 length=3334
PF3D7_1105600 length=777
PF3D7_0305000 length=390
PF3D7_0507700 length=531
PF3D7_1408700 length=7182
PF3D7_0104200 length=466
PF3D7_0921400 length=247
Non-denatured protein fraction
Temperature
DMSO.r1
DMSO.r2
Med6.r1
Med6.r2

### Med6 CETSA data plotting_curve fitting

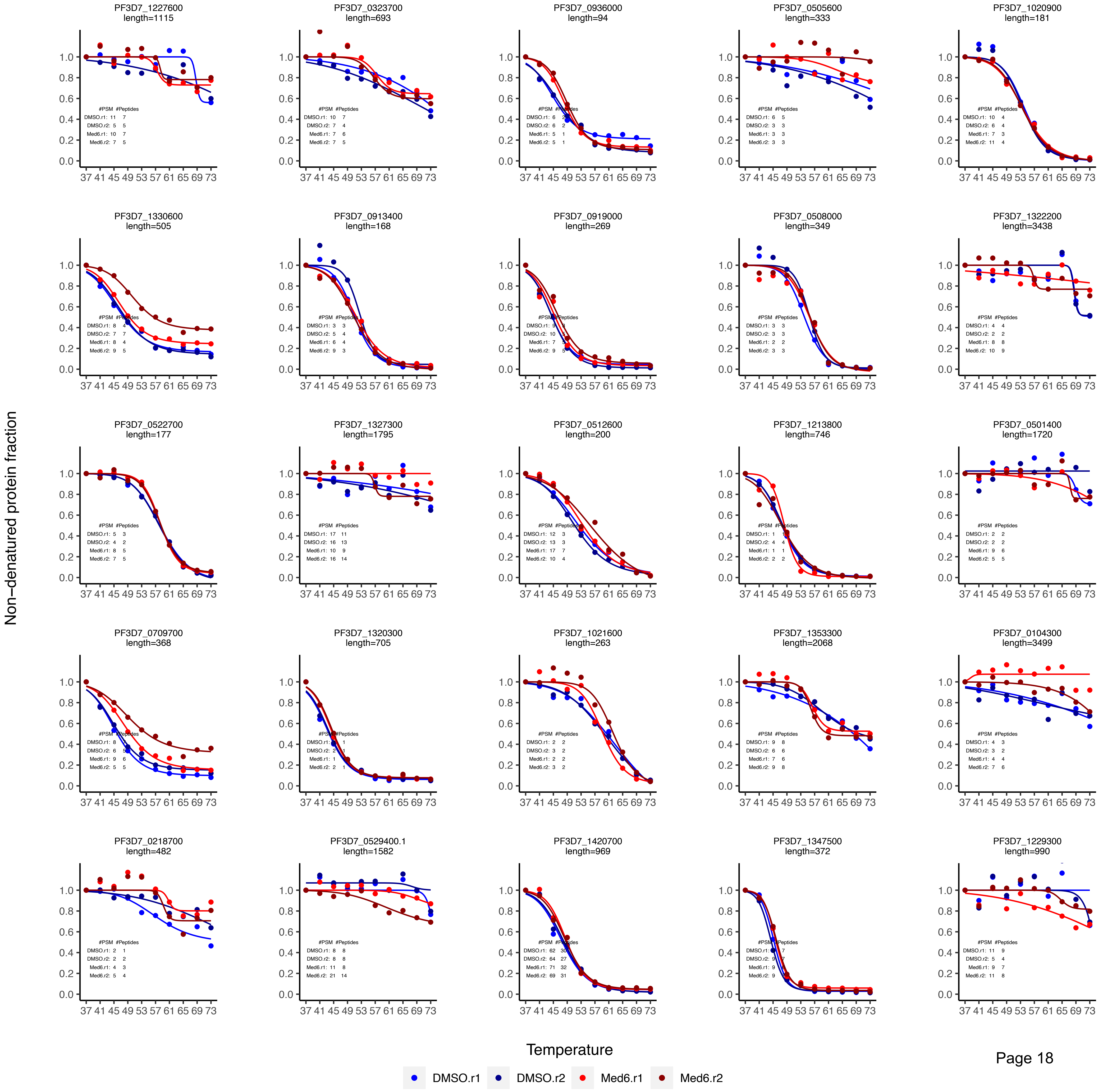

Med6 CETSA data plotting_curve fitting

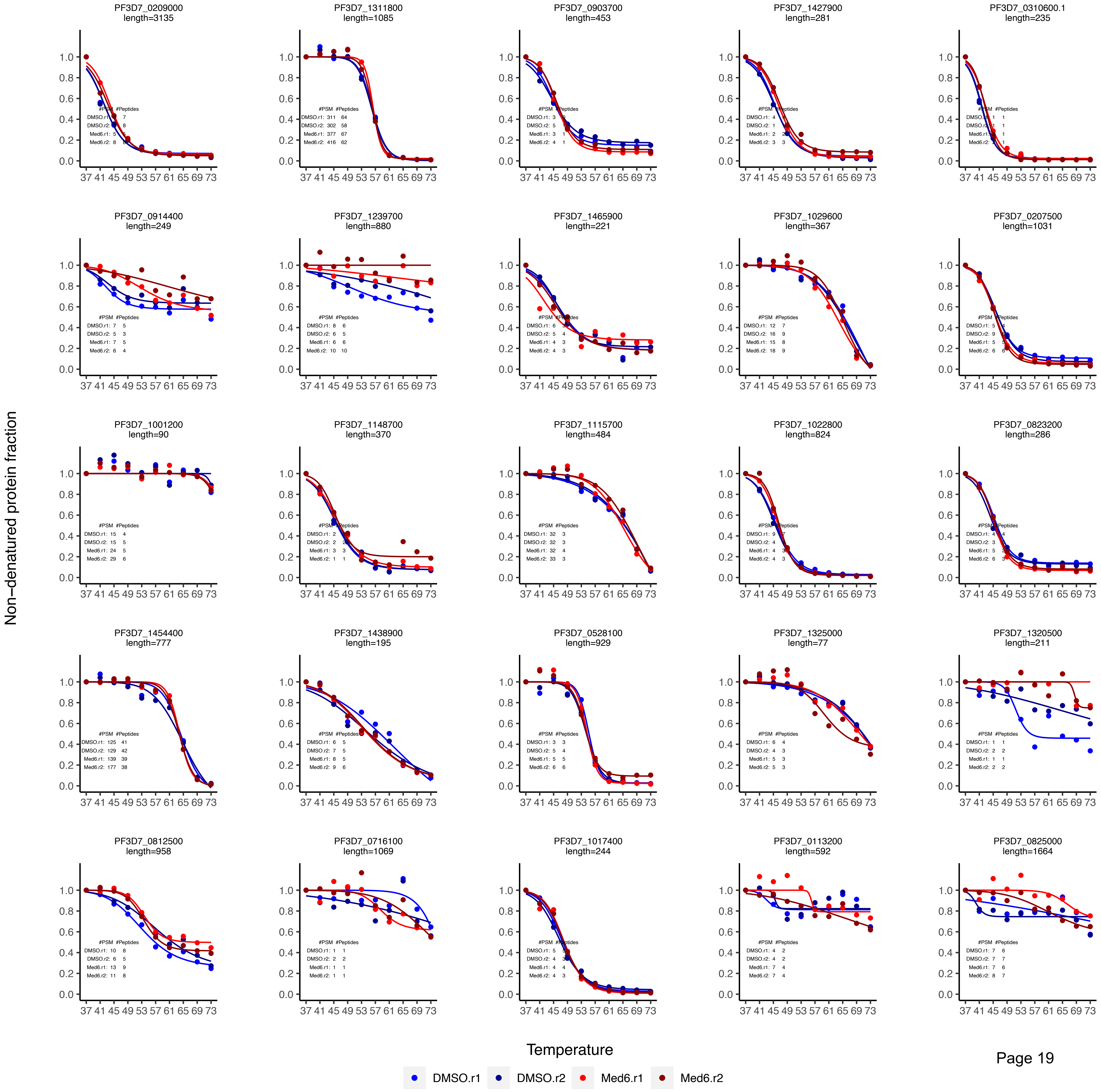

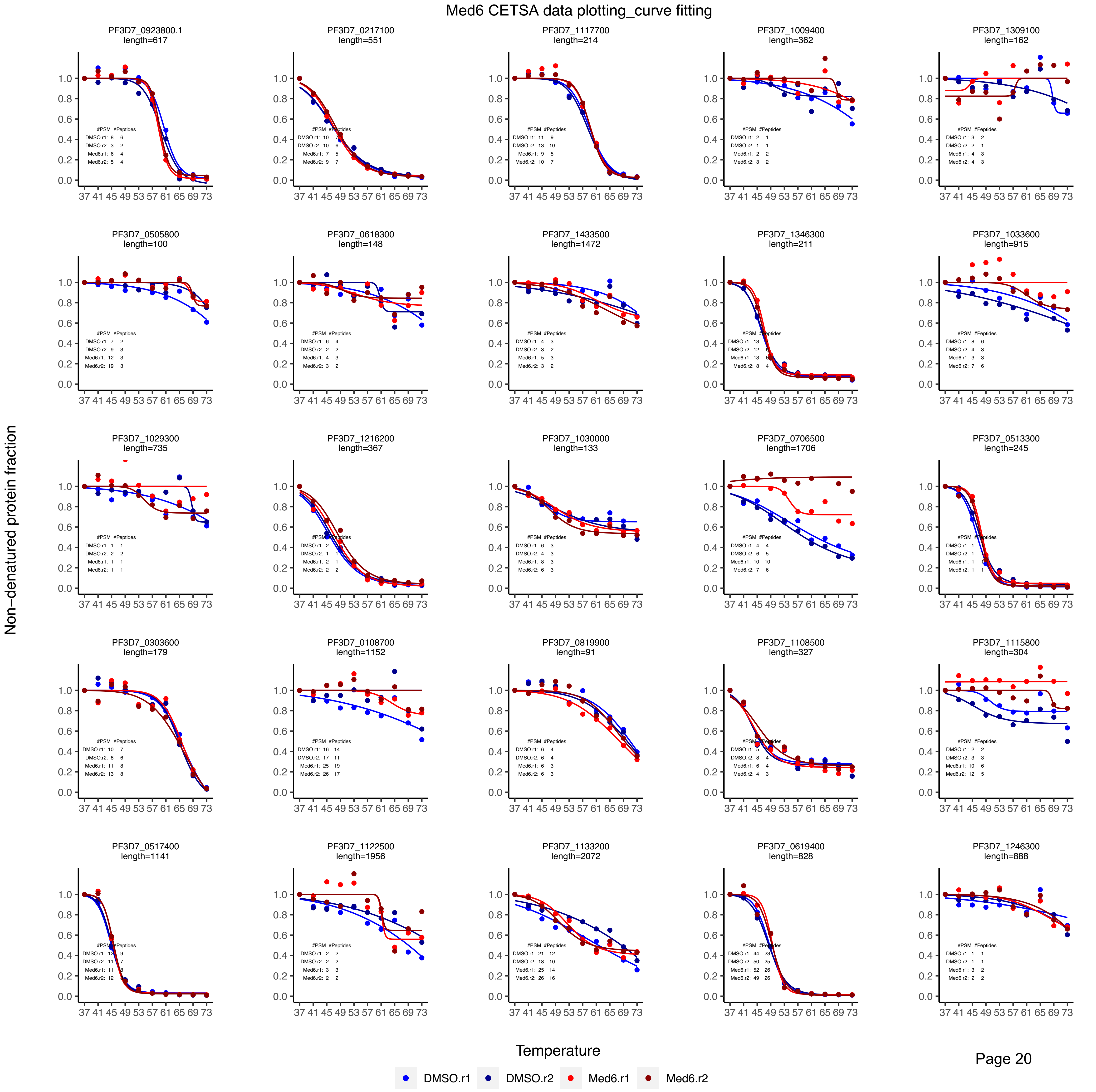
Med6 CETSA data plotting_curve fitting
PF3D7_0923800.1
length=617
PF3D7_0217100
length=551
PF3D7_1117700
length=214
PF3D7_1009400
length=362
PF3D7_1309100
length=162
PF3D7_0505800
length=100
PF3D7_0618300
length=148
PF3D7_1433500
length=1472
PF3D7_1346300
length=211
PF3D7_1033600
length=915
PF3D7_1029300
length=735
PF3D7_1216200
length=367
PF3D7_1030000
length=133
PF3D7_0706500
length=1706
PF3D7_0513300
length=245
PF3D7_0303600
length=179
PF3D7_0108700
length=1152
PF3D7_0819900
length=91
PF3D7_1108500
length=327
PF3D7_1115800
length=304
PF3D7_0517400
length=1141
PF3D7_1122500
length=1956
PF3D7_1133200
length=2072
PF3D7_0619400
length=828
PF3D7_1246300
length=888
Non−denatured protein fraction
Temperature
DMSO.r1
DMSO.r2
Med6.r1
Med6.r2

Med6 CETSA data plotting_curve fitting

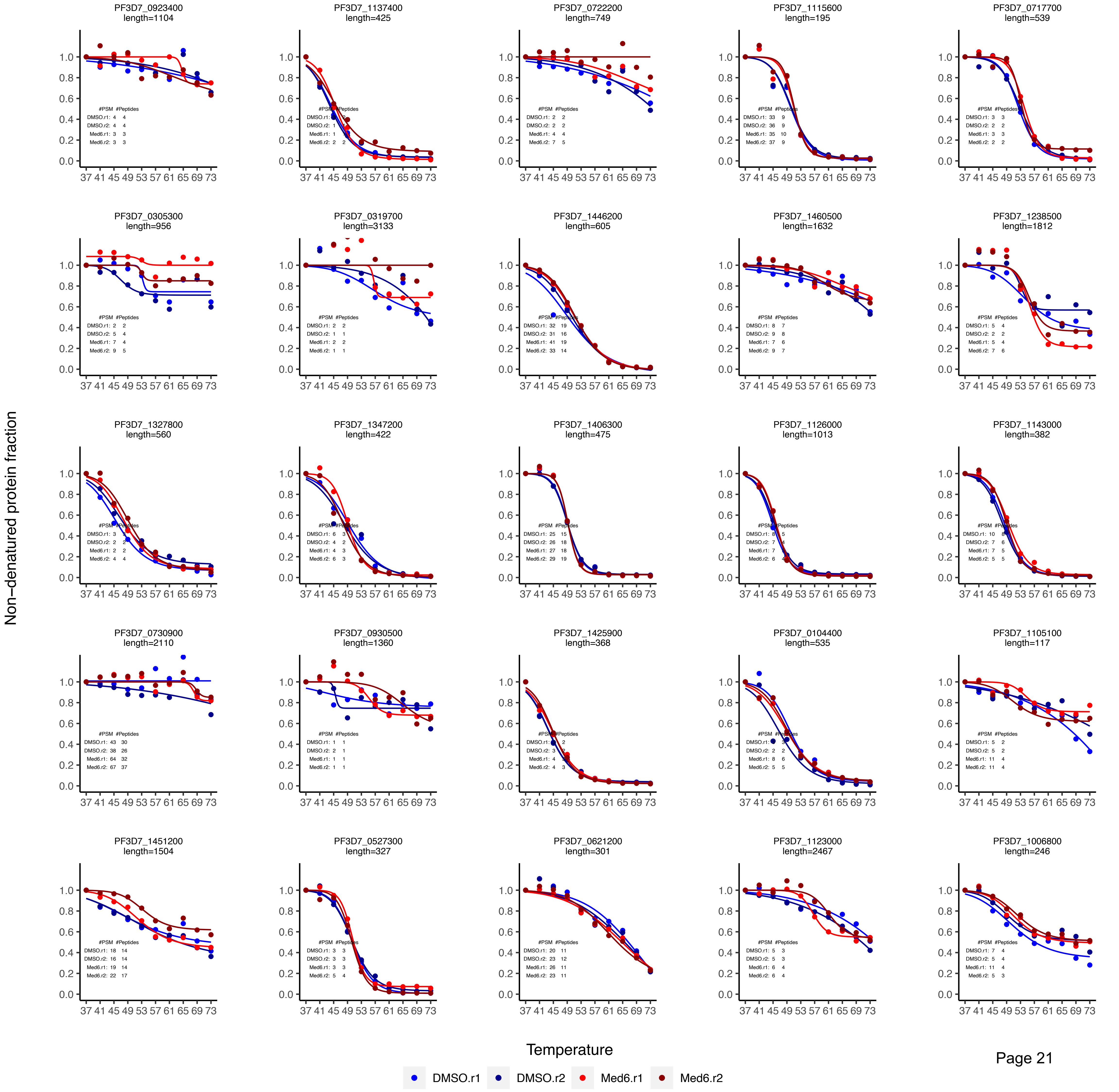

Med6 CETSA data plotting_curve fitting

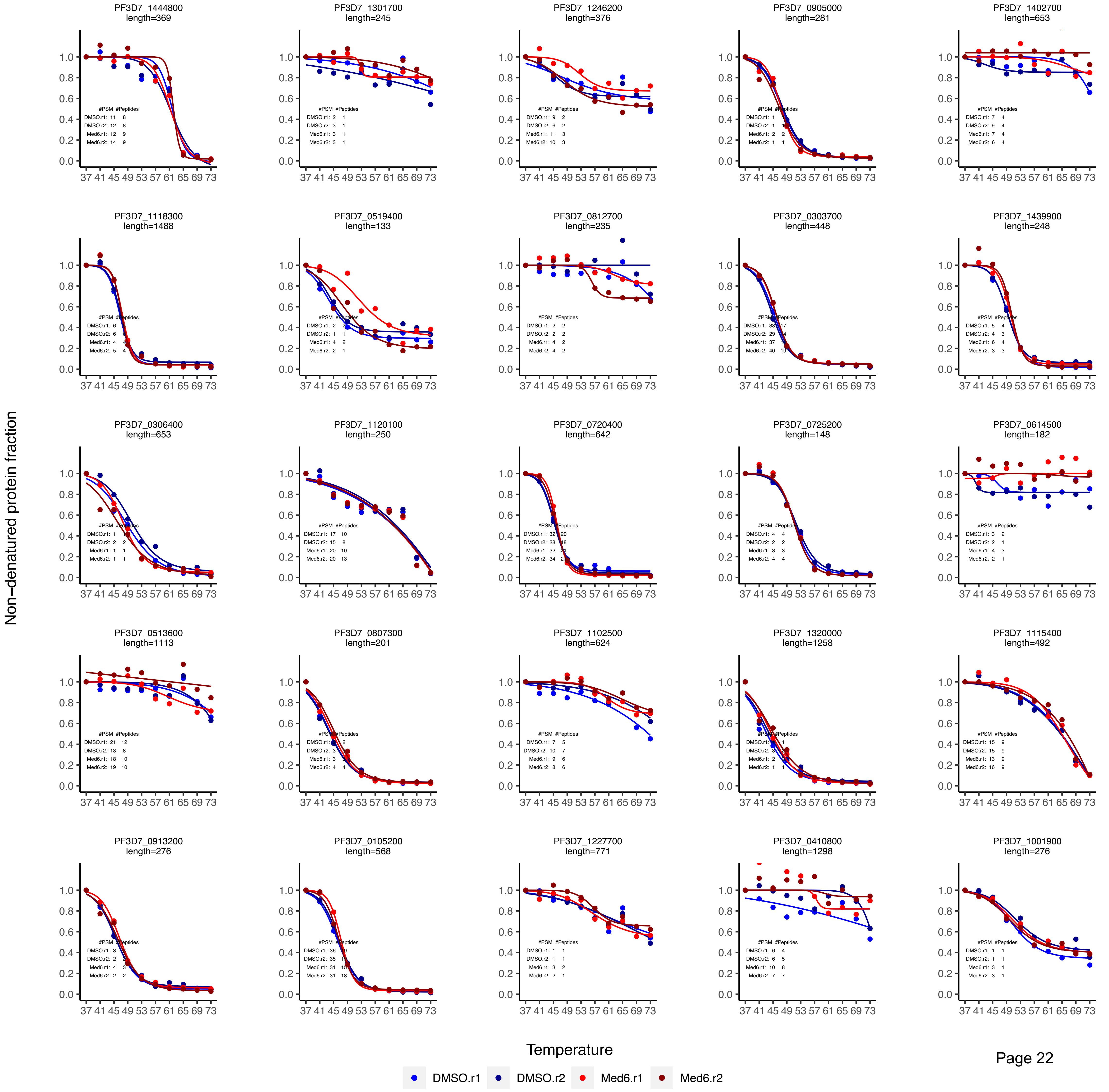

Med6 CETSA data plotting_curve fitting

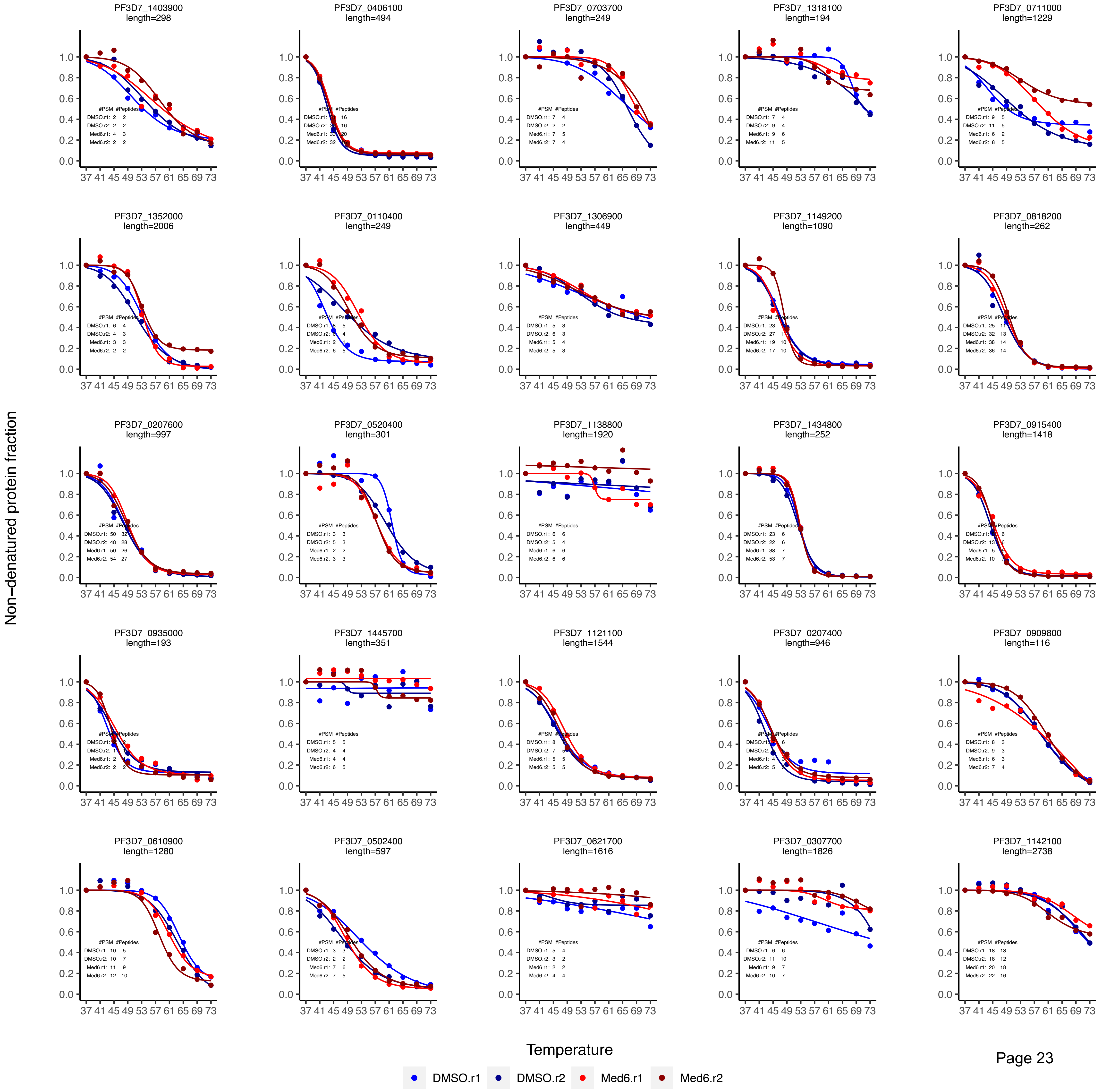

Med6 CETSA data plotting_curve fitting

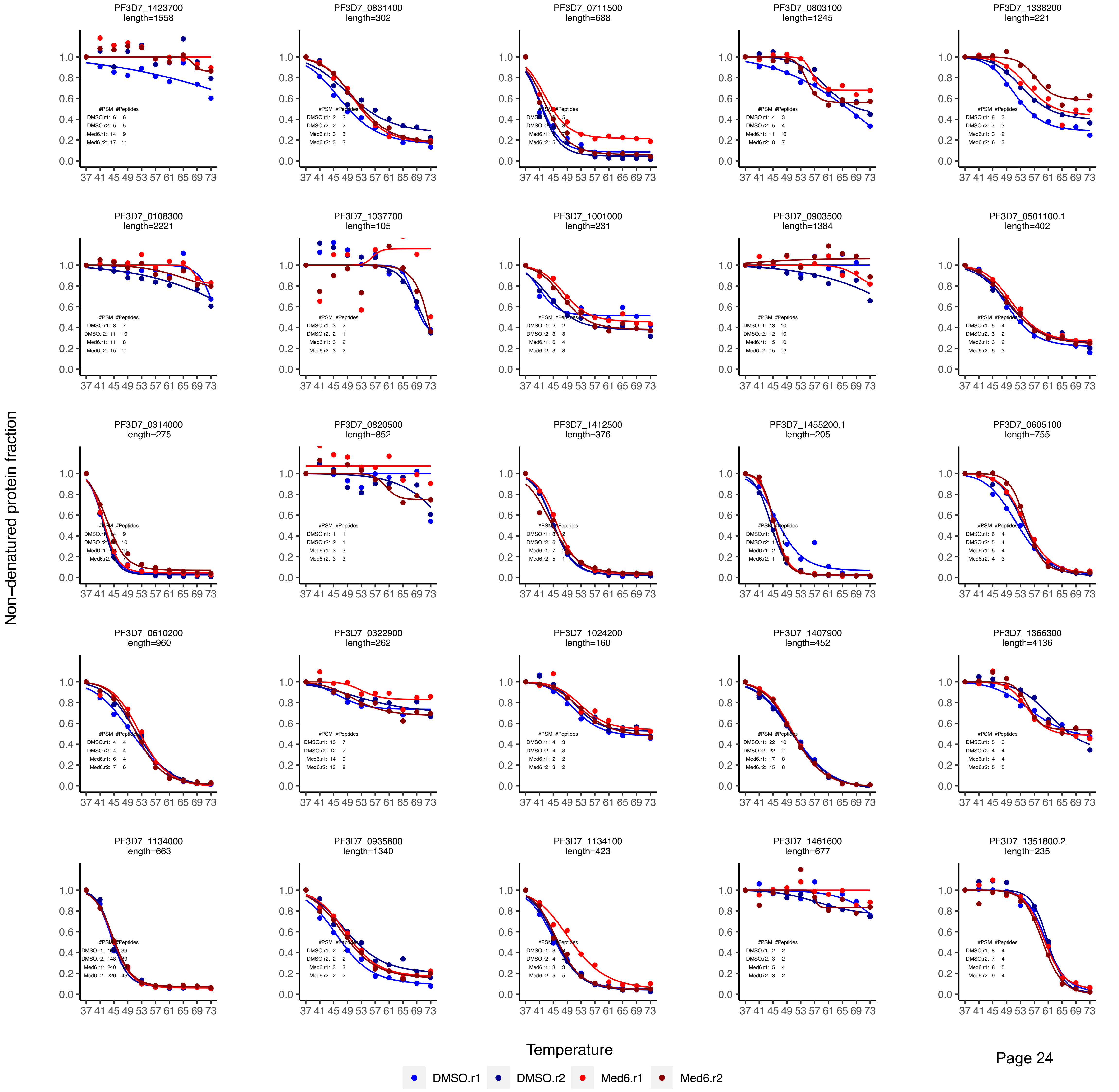

Med6 CETSA data plotting_curve fitting

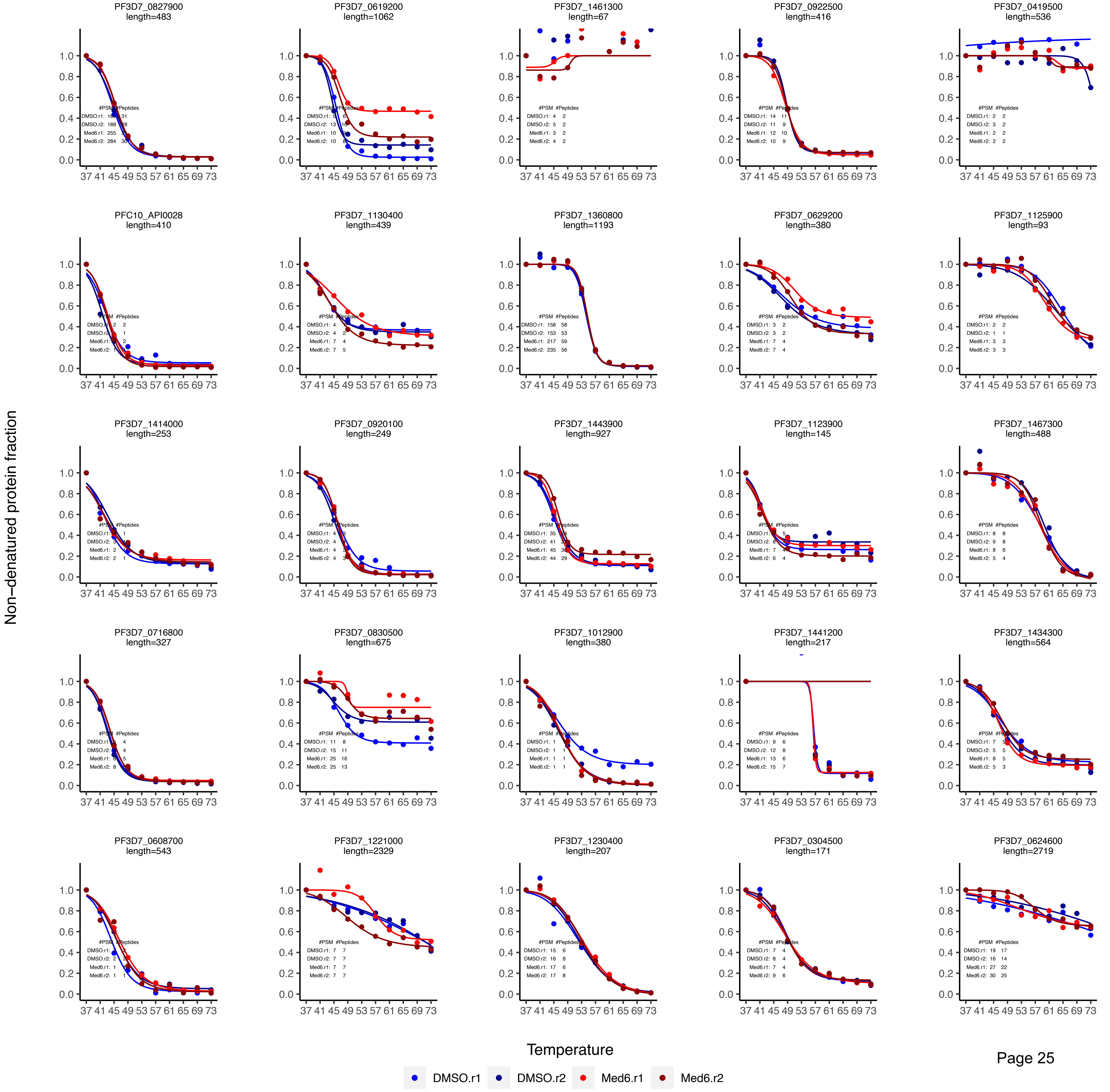

### Med6 CETSA data plotting_curve fitting

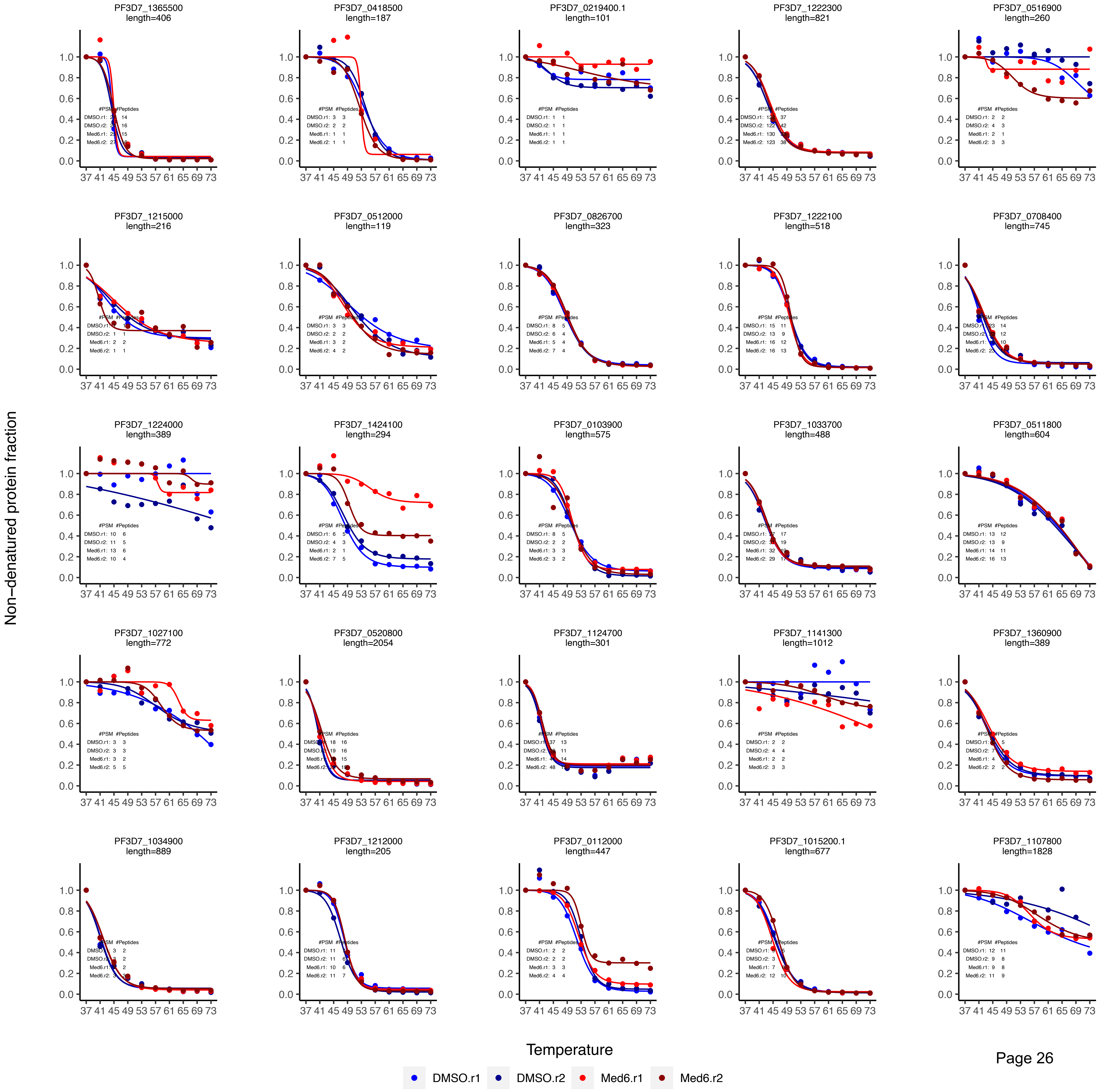

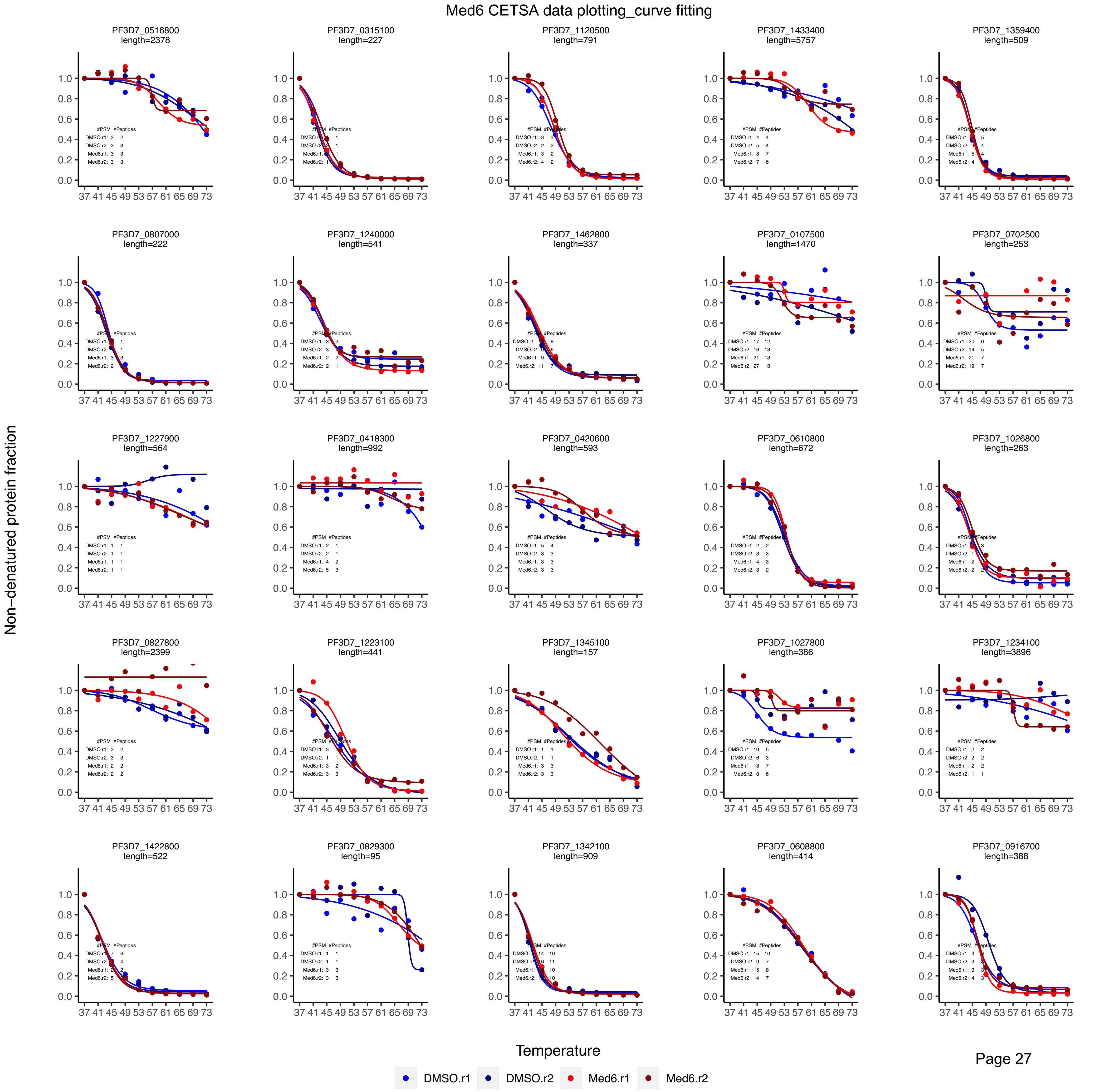
Med6 CETSA data plotting_curve fitting
PF3D7_0516800 length=2378
PF3D7_0315100 length=227
PF3D7_1120500 length=791
PF3D7_1433400 length=5757
PF3D7_1359400 length=509
PF3D7_0807000 length=222
PF3D7_1240000 length=541
PF3D7_1462800 length=337
PF3D7_0107500 length=1470
PF3D7_0702500 length=253
PF3D7_1227900 length=564
PF3D7_0418300 length=992
PF3D7_0420600 length=593
PF3D7_0610800 length=672
PF3D7_1026800 length=263
PF3D7_0827800 length=2399
PF3D7_1223100 length=441
PF3D7_1345100 length=157
PF3D7_1027800 length=386
PF3D7_1234100 length=3896
PF3D7_1422800 length=522
PF3D7_0829300 length=95
PF3D7_1342100 length=909
PF3D7_0608800 length=414
PF3D7_0916700 length=388
Non−denatured protein fraction
Temperature
DMSO.r1
DMSO.r2
Med6.r1
Med6.r2

Med6 CETSA data plotting_curve fitting

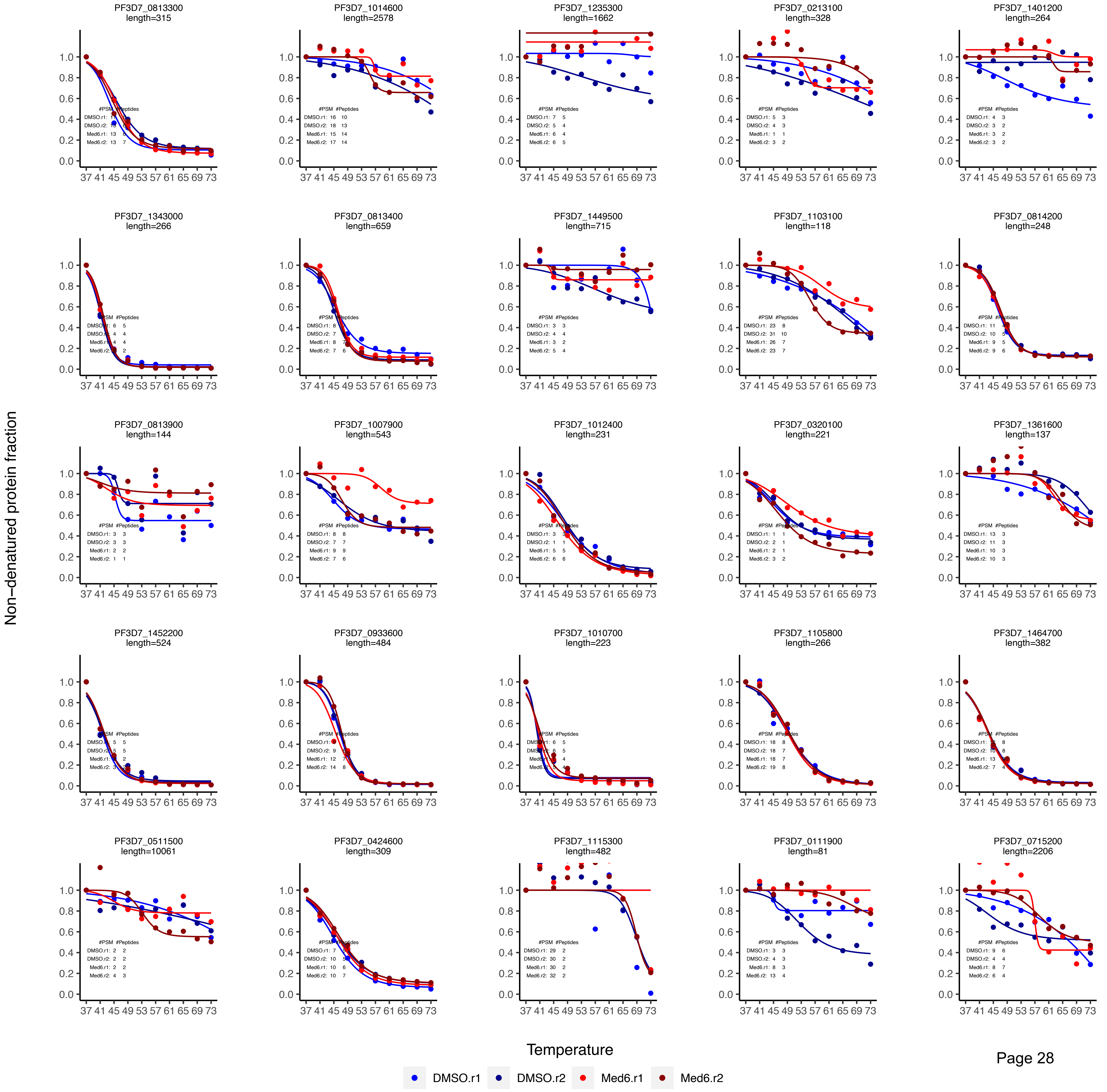

Med6 CETSA data plotting_curve fitting

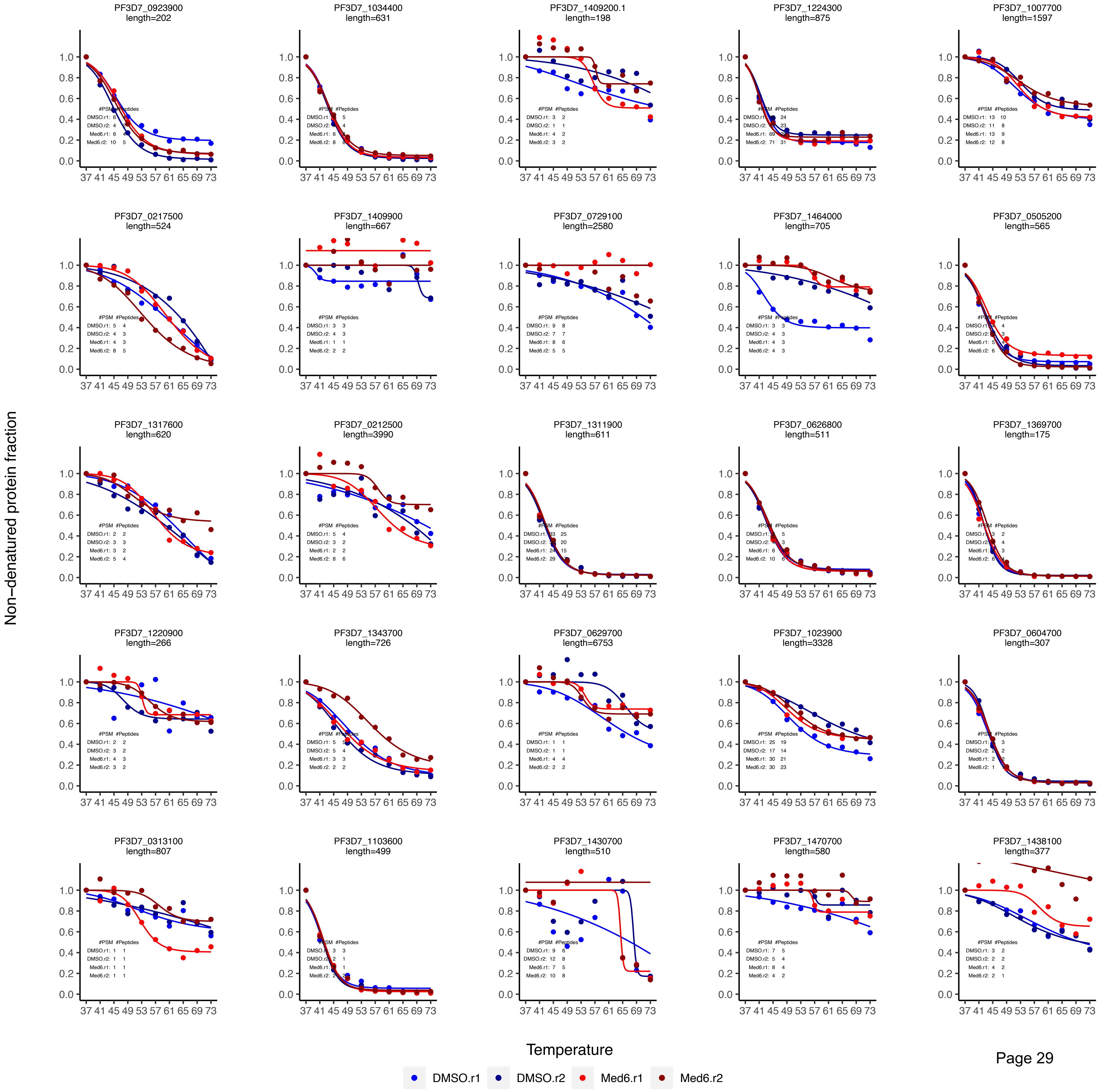

### Med6 CETSA data plotting_curve fitting

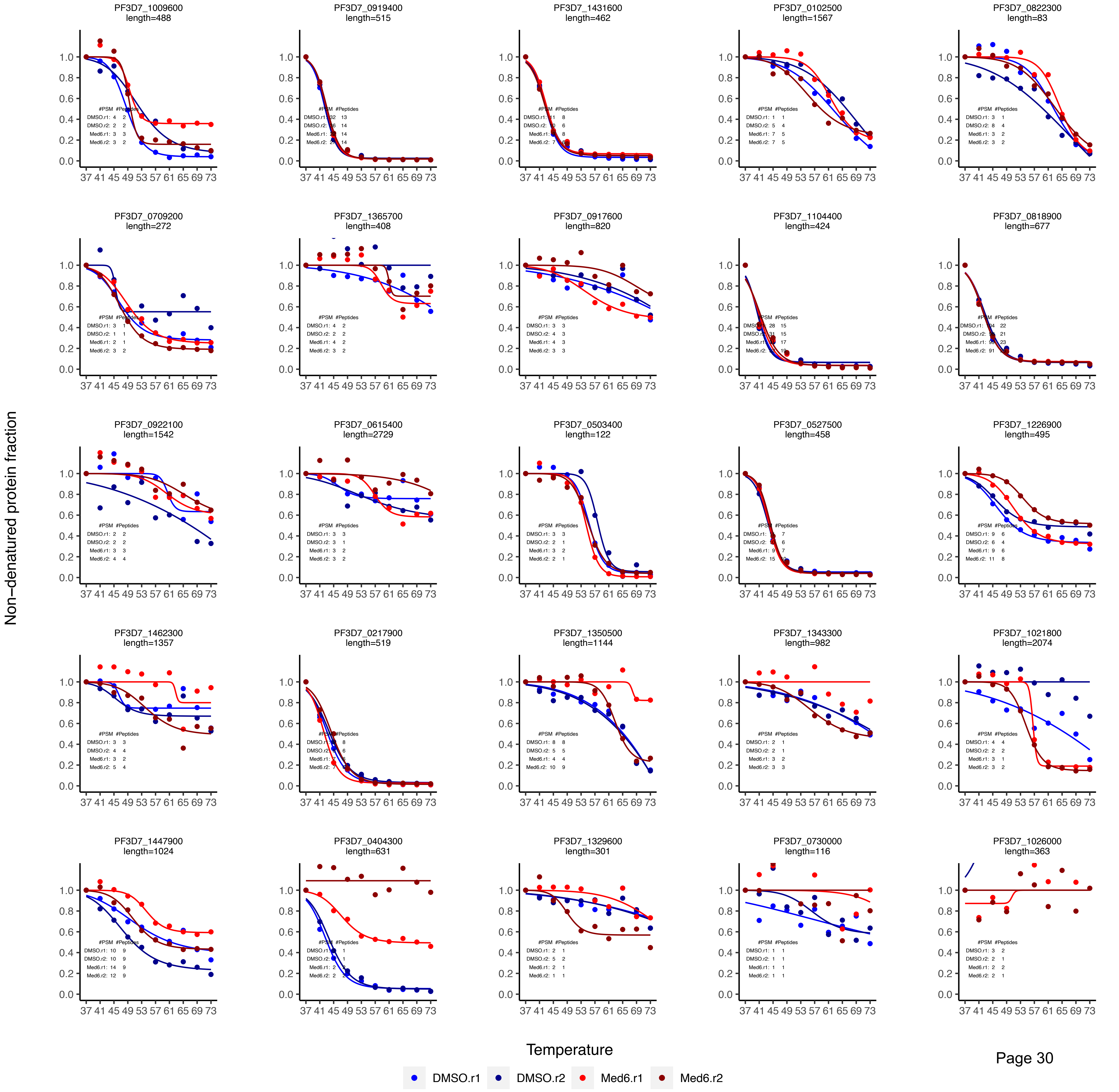

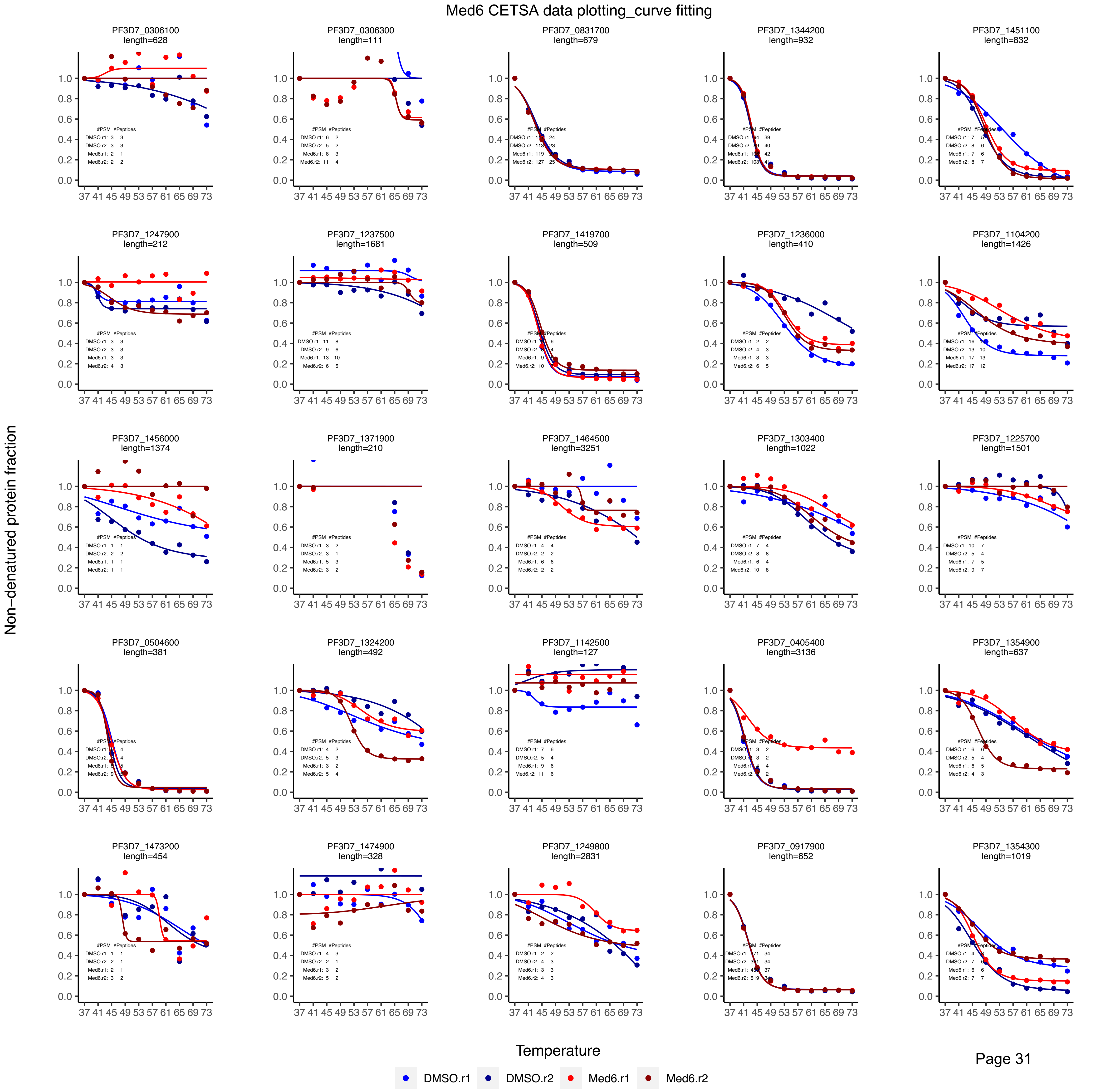
Med6 CETSA data plotting_curve fitting
PF3D7_0306100
length=628
PF3D7_0306300
length=111
PF3D7_0831700
length=679
PF3D7_1344200
length=932
PF3D7_1451100
length=832
PF3D7_1247900
length=212
PF3D7_1237500
length=1681
PF3D7_1419700
length=509
PF3D7_1236000
length=410
PF3D7_1104200
length=1426
PF3D7_1456000
length=1374
PF3D7_1371900
length=210
PF3D7_1464500
length=3251
PF3D7_1303400
length=1022
PF3D7_1225700
length=1501
PF3D7_0504600
length=381
PF3D7_1324200
length=492
PF3D7_1142500
length=127
PF3D7_0405400
length=3136
PF3D7_1354900
length=637
PF3D7_1473200
length=454
PF3D7_1474900
length=328
PF3D7_1249800
length=2831
PF3D7_0917900
length=652
PF3D7_1354300
length=1019
Non−denatured protein fraction
Temperature
DMSO.r1
DMSO.r2
Med6.r1
Med6.r2

Med6 CETSA data plotting_curve fitting

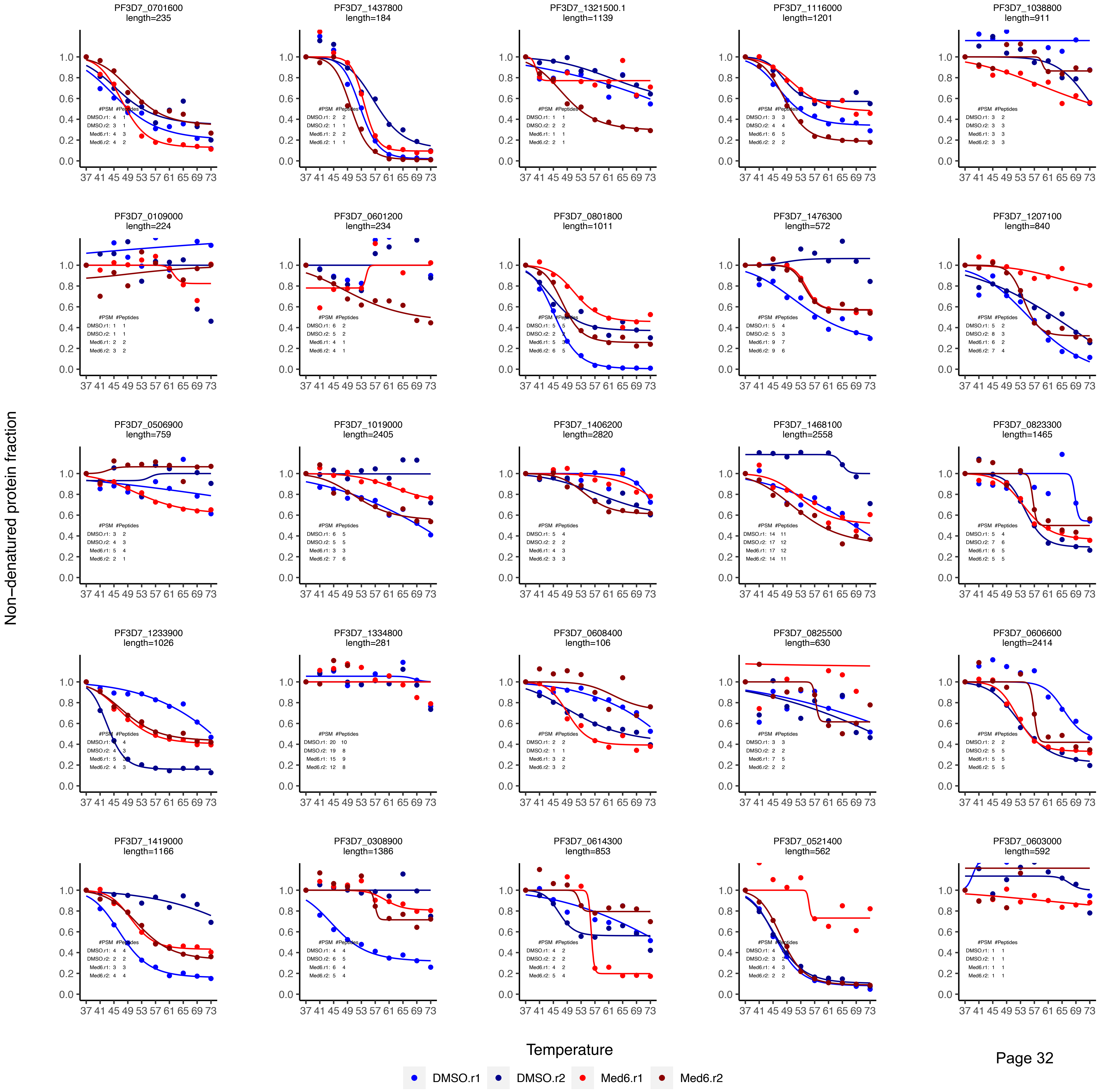

Med6 CETSA data plotting_curve fitting

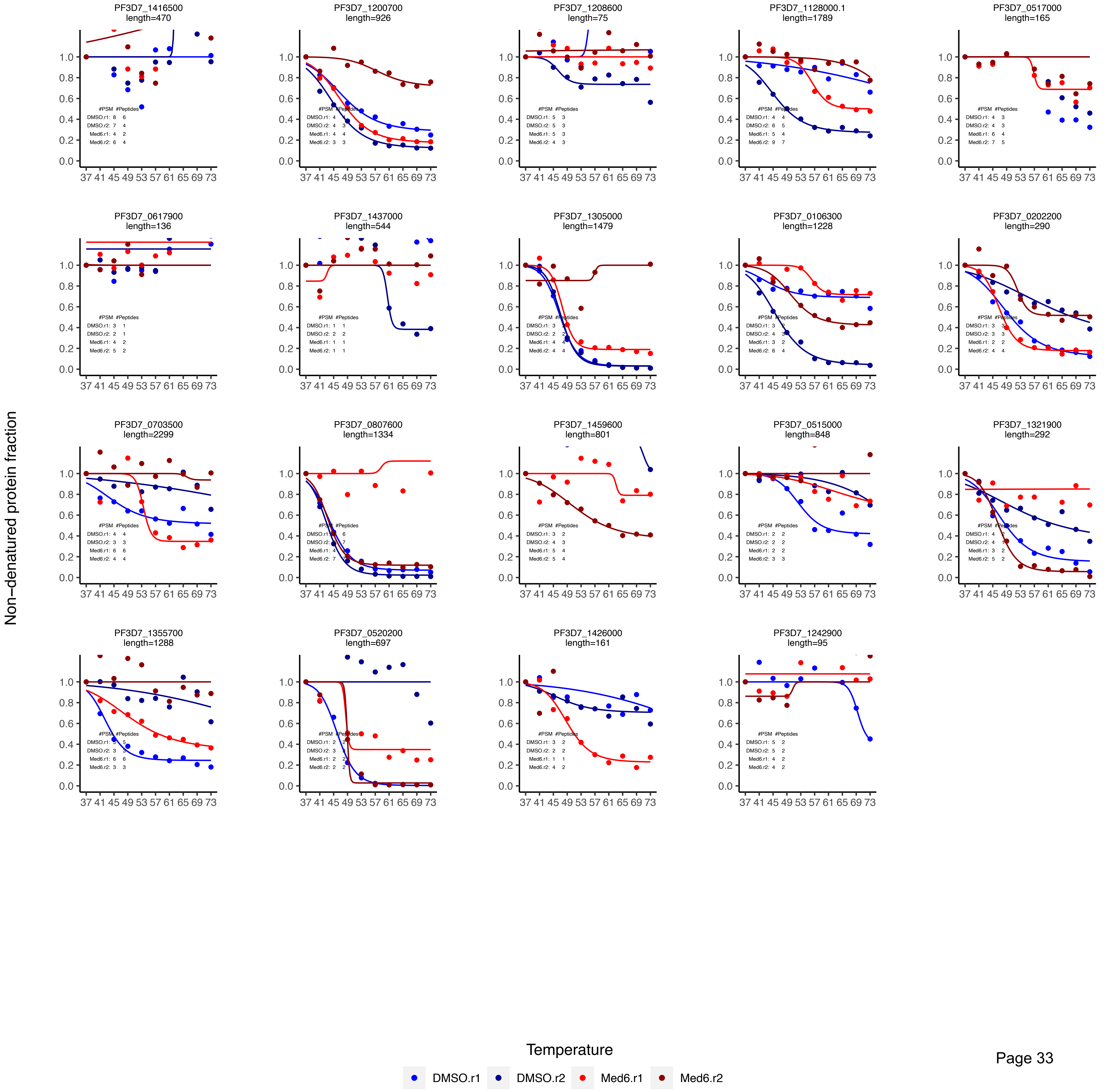
